# AI-enabled rhodopsin design for blue-light enhanced bacterial growth

**DOI:** 10.64898/2026.06.29.735265

**Authors:** Haris Saeed, Michael J. Lewis, Takayoshi Fujiwara, Jek Huang, Masae Konno, Kaho Mori, Susumu Yoshizawa, Keiichi Inoue, Tiezheng Pan, Yun Wang, Aidong Yang, Wei E. Huang

## Abstract

We developed an AI-guided design pipeline that generated and validated non-natural microbial rhodopsins with spectral properties not yet known in nature. The pipeline comprised a three-stage *in silico* design, a genetic algorithm (GA) for sequence generation, a stacked LASSO and XGBoost machine-learning (ML) regressor for spectral prediction and fitness ranking, and a Markov-based sequence plausibility filter to enforce proton pumping like characteristics. Four candidate rhodopsins (APR1, APR2, APR6, and APR7) targeting blue light absorption were designed and AlphaFold3 structural modelling predicted retinal binding pocket architecture consistent with outward proton-pumping function. Experimental characterisation confirmed that all four variants absorbed light at ∼410 nm and significantly promoted the growth of *Cupriavidus necator* under blue light illumination. This study demonstrates that AI-enabled design can engineer proteins with no natural precedent, generating light-harvesting rhodopsins with novel spectral properties while preserving biological function, marking a significant advance in programmable synthetic biology.

## Introduction

Microbial carbon dioxide (CO_2_) fixation through photosynthesis is one of the foundations of the global carbon cycle^1, 2^. Photosynthetic microbes harvest solar radiation to convert CO_2_ and water into organic compounds, contributing around 50% of primary productivity on earth^3^. In nature, almost all known light-harvesting mechanisms in microorganisms are either chlorophyll- or rhodopsin-based systems^4^. Chlorophyll-based photosystems are highly complex, comprising multi-component assemblies and electron transport chains that couple light harvesting to water splitting, ATP generation, and reductant production. In comparison, rhodopsin-based photosystems are much simpler with only a light-activated proton pump generating proton motive force^5^. Despite this simplicity, increasing evidence suggests that rhodopsin phototrophy can support ATP synthesis, reverse electron transfer, and regeneration of reducing equivalents required for CO₂ fixation ^6, 7, 8^. We recently converted a non-photosynthetic bacterium *Cupriavidus necator* (previously *Ralstonia eutropha* H16) into a photoelectrosynthetic organism by engineering *Gloeobacter* rhodopsin (GR) and extracellular electron uptake system (e.g. riboflavin and MtrCBA) ^6^ ^8^. This photoelectrosynthetic *C. necator* H16 is able to grow using light as the primary energy source (coupled with electricity generated by a solar panel) and CO_2_ as the sole carbon source. These findings demonstrate that microbial rhodopsins could serve as modular building blocks for engineering simplified artificial photosynthetic systems^6^ ^8^.

Microbial rhodopsins have attracted broad interest in synthetic biology, bioenergy, and optogenetics ^4, 6, 8, 9, 10, 11, 12, 13, 14^. Microbial rhodopsins are widely used as optogenetic tools in synthetic biology and have been successfully expressed in model bacteria such as *Escherichia coli*^14^, *C. necator*^6, 8^ and *Shewanella oneidensis*^15, 16^. Many studies demonstrated that the engineered bacteria with rhodopsin can convert light energy into intracellular chemical energy^6^ ^8, 13, 17, 18^. Engineering rhodopsins with altered photophysical properties, such as blue-and red- shifted absorption spectra, could expand their broader potential by enabling spectral multiplexing, improving compatibility with defined illumination regimes and increasing the flexibility of artificial photosynthetic platforms. Spectral engineering of rhodopsins remains a challenge. First, accurate prediction of quantum yield (QY), photocycle kinetics and absorption often requires intensive and expensive computational simulations. Hybrid quantum mechanics and molecular mechanics approaches such as Automatic Rhodopsin Modeling (ARM) ^19^ can achieve high predictive accuracy, but often require days to weeks of computation per sequence, making large scale sequence space exploration difficult. Second, blue- shifted proton pumping rhodopsins are rare in nature. Within a curated set of 884 collected rhodopsins ^20^ ^21^, only a small fraction (about 3.5%) of proton pumps occupy the blue-absorbing region, making the design of extreme blue-shifted phenotypes difficult. Random mutagenesis is inefficient because most mutations are deleterious and searches rapidly plateau before reaching rare and functional rhodopsin variants with extreme spectral properties.

Artificial intelligence (AI) ^22, 23, 24, 25^ and machine learning (ML) ^20, 21^ ^26, 27, 28^ provide a potential route to accelerate rhodopsin engineering by enabling large-scale *in silico* screening. However, ML trained predominantly on naturally occurring rhodopsin sequences often generalise poorly when extrapolating toward rare or extreme spectral phenotypes. Previous ML-guided rhodopsin engineering studies have largely remained within the natural spectral range and did not efficiently access blue-shifted variants ^27, 28, 29, 30^. Accessing highly blue-shifted proton-pump-like rhodopsins therefore requires not only a fast spectral predictor, but also a sequence-search strategy that can explore rare regions of sequence space while maintaining structural and functional plausibility.

To address this limitation, we developed an integrated AI-guided evolutionary framework for targeted rhodopsin design. The platform comprises three integrated computational stages: a genetic algorithm (GA) for sequence generation and optimisation, a λ_max_ predictor based on stacked LASSO and XGBoost models for fitness ranking, and a Markov-model-based sequence filter that constrains the search to aligned sequences predicted to retain proton-pump-like functionality and stability. This design - score - filter workflow enables efficient exploration of rare spectral regions while maintaining biophysical plausibility (Figure 1).

**Figure 1.**
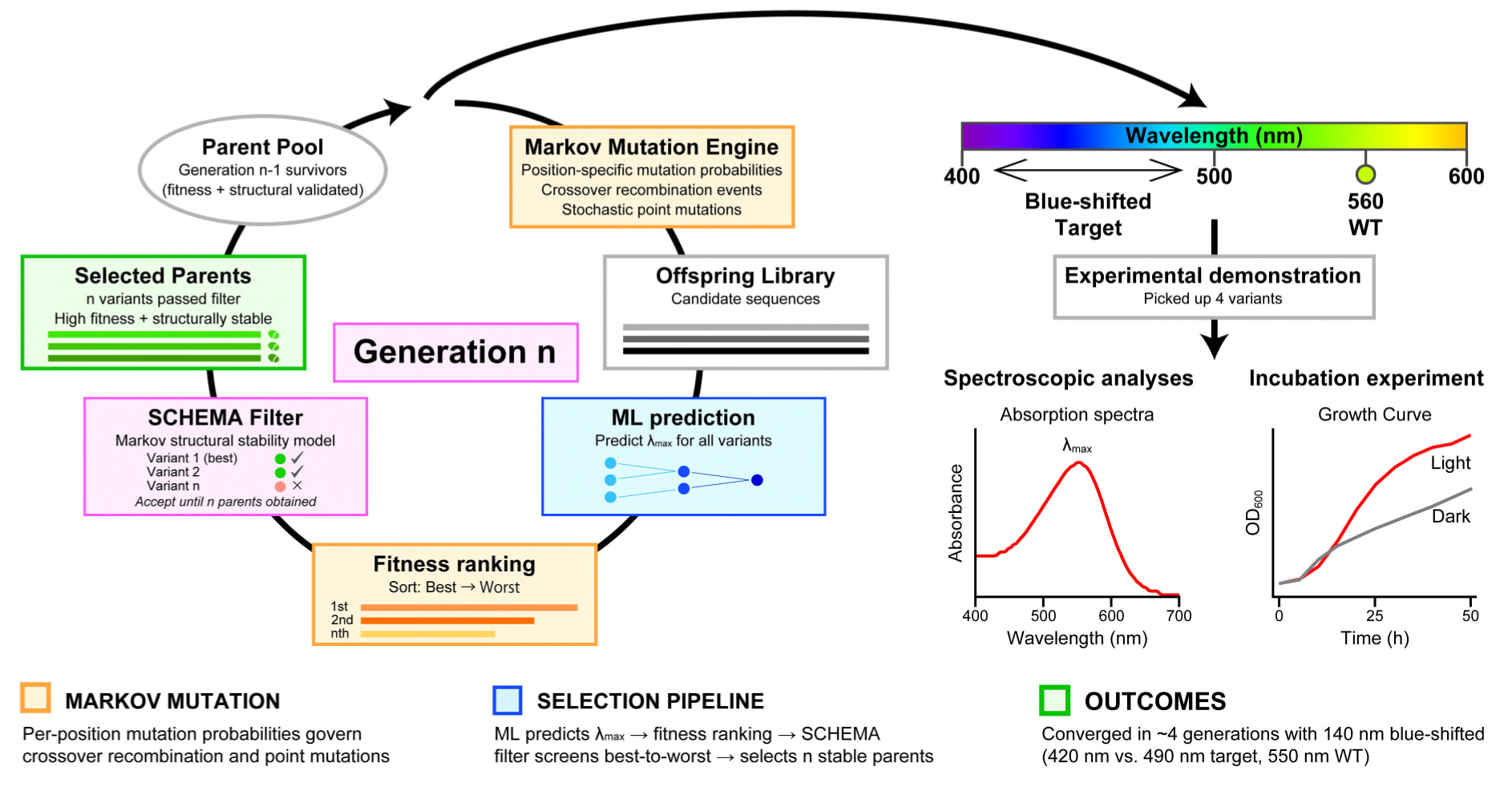
AI-guided directed-evolution pipeline. A GA proposes variants, an ML regressor scores predicted λ_max_, and a Markov model rejects sequence-implausible candidates before they can seed later generations. Selected candidates are then assayed experimentally.

Here, we apply this framework to design and biologically validate blue-shifted microbial rhodopsins. We present a ML predictor for λ_max_ optimised for computational efficiency (< 10 ms per prediction), enabling real-time guidance during GA search. Our sensitivity analysis shows the MI-weighted mutation prior accelerates convergence (median 11 vs 23 generations with inverse-MI prior at target 490 nm, an ∼2-fold improvement) and candidate quality. Experimental validation of top ranked candidates confirms this framework has successfully generated four blue-shifted, functional proton pump rhodopsins, which enhanced growth activity in *C. necator* H16 under the blue light. Together, these results establish a scalable strategy for data-driven rhodopsin engineering and provide a foundation for the development of next-generation artificial photosynthetic systems and optogenetic tools.

## Results

### Computational design of rhodopsins

The detailed methods and algorithms of computational design of rhodopsins are provided in Computational supplementary information.

### Phylogenetic analysis of rhodopsins

Blue-shifted proton pump design is constrained by the natural diversity in microbial rhodopsins (Figure 2). Analysis of 884 rhodopsin sequences ^20^ ^21^ across six functional classes shows that wavelength distributions are separated by function (Figure 2). Proton-pump rhodopsins are well represented in the dataset (n = 310; 35.1% of all sequences), but they are rare in the blue region: only 11 of 310 proton-pump sequences (3.5%) have a peak absorption wavelength below 500 nm, and blue-extreme proton pumps below 470 nm account for <1% of all samples. The median *λ*_max_ of proton-pump rhodopsins is 548 nm, by contrast, channelrhodopsins (n = 37) have a median *λ*_max_ of 480 nm, representing a 68 nm blue-shift relative to proton-pump rhodopsins (Figure 2c). Thus, blue-shifted absorption is physically achievable within microbial rhodopsins but is concentrated outside the proton-pump lineage.

**Figure 2.**
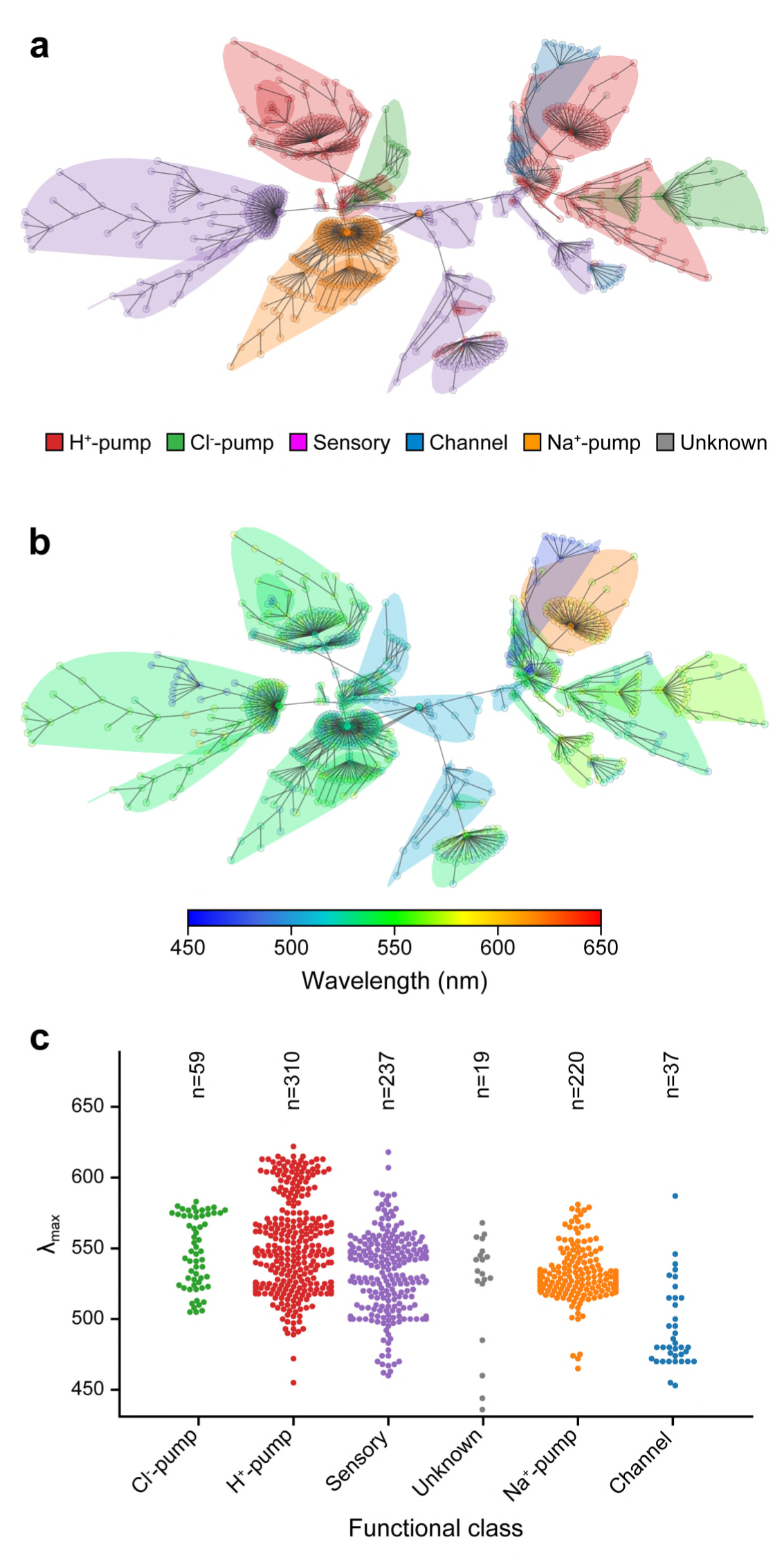
Sequence similarity, structural conservation, and spectral distribution of rhodopsins. (a) Primary-sequence MST coloured by functional class; pump sequences form the dominant cluster. (b) Same MST coloured by λ_max_; most pump sequences absorb in the green (520-570 nm), while blue-absorbing clusters are sparse. (c) λ_max_ across functional classes; only 3.5% of pump sequences absorb below 500 nm. Secondary-structure MSTs and distance distributions are in Computational Supplementary Information Figs. S3 and S4.

A Minimum Spanning Tree (MST) over pairwise Levenshtein distances shows that functional classes occupy distinct regions of sequence space, with the proton-pump core well-separated from blue-absorbing clusters (Figure 2a, b). Green proteorhodopsin (GPR, 126 sequences) and bacteriorhodopsin (BR, 97 sequences) together account for >70% of the proton-pump cluster, reflecting the evolution paths on these two constructs.

Despite their shared function, proton-pump rhodopsins are highly divergent at the sequence level. Within the 310 proton-pump cluster, position-by-position aligned-sequence identity (including gap columns) has a median of 8.84% (mean 29.43%). The mean is right-skewed by clusters of near-duplicate wildtype variants (e.g. the GPR and BR mutational series), while the low median reflects the large fraction of aligned positions that are gaps for any given pair. As most of this variation is concentrated in tail regions; the mean secondary-structure identity was 74.03%, and 95.8% of sequences have at least one close homolog (E < 0.01) in the dataset (Biological Supplementary information Table S1). Within-class pairwise Levenshtein distances show internal conservation of the proton-pump cluster relative to channelrhodopsins (Computational Supplementary information Figure CS1).

These analyses make evident a spectral and sequence-space gap between blue absorption and proton-pump function in rhodopsins.

### Rhodopsin wavelength prediction and sequence-plausibility filtering

#### Wavelength prediction

To bridge this gap, we designed two components: a sequence based *λ*_max_ regressor (Computational Supplementary Information methods), and a sequence-plausibility filter to constrain evolutionary search within the proton-pump lineage while permitting exploration of blue-shifted variants. A stacked LASSO-XGBoost regressor using physicochemical Feature Map encoding achieved R^2^ = 0.840, RMSE = 12.64 nm, and MAE = 8.45 nm under shuffled 10-fold cross-validation (Figure 3a), outperforming the best linear model (R^2^ = 0.783; ΔR^2^ = 0.057; Computational Supplementary Information Table CS10), and alternative encodings, including one-hot (R^2^ = 0.812) and rank-based ordinal (R^2^ = 0.750) (Computational Supplementary Information Table CS10). Because the genetic algorithm only propagates sequences that pass the proton-pump plausibility filter, shuffled cross-validation provides a direct estimate of in-lineage interpolation performance. However, the training set contains dense clusters of closely related variants from the GPR and BR mutational series (Computational Supplementary Information Figure CS20), which can inflate shuffled-CV estimates for more remote candidates. Thus, we consider spectral-tail validation, where the extreme 5% of wavelengths from each end of the distribution are held out (R^2^ = 0.648, RMSE = 33.95 nm; Figure 3d), and wildtype-grouped CV, in which all variants sharing the same parent scaffold are forced into the same fold (RMSE 10.1-29.1 nm across groups Computational Supplementary Table CS11). Inference takes < 10 ms per sequence on a single CPU core, fast enough for in-loop GA scoring.

**Figure 3.**
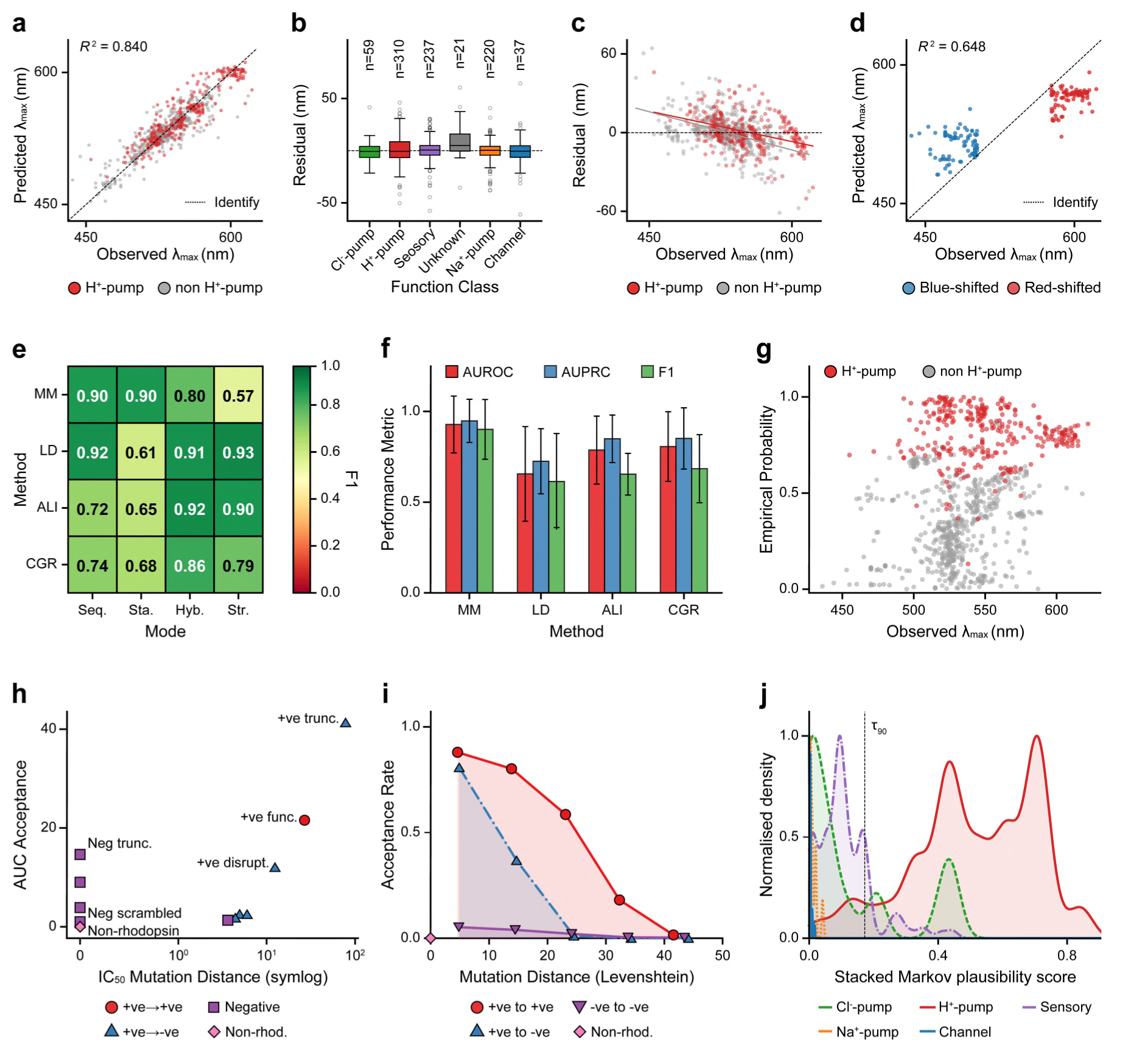
Wavelength predictor diagnostics and sequence-plausibility filter evaluation. (a) Predicted vs observed λ_max_ under 10-fold cross-validation (R^2^=0.840, MAE =8.45 nm); dashed line is identity. (b) Residuals by functional class; distributions are approximately zero-centred. (c) Residuals vs observed λ_max_; the negative trend (slope ≈ −0.17, p<0.001) shows regression-to-the-mean compression. (d) Spectral-tail variants; blue-shifted variants are overestimated by up to 80 nm, consistent with regression-to-the-mean compression at the distributional tails. (e) F1 scores across filter method-mode combinations; sequence-aware and stacked models perform best, while structure-only Markov performs poorly (median F1 = 0.618), consistent with conserved rhodopsin folds. (f) Stacked classification performance; Markov gives the highest median subfamily AUROC among the stacked filters. (g) Filter score shows no systematic correlation with λ_max_ in proton pumps (R^2^=0.002). (h) IC_50_ mutation distance versus integrated AUC acceptance; functional variants are accepted at larger mutation distances, while non-rhodopsins are rejected near distance zero. (i) Acceptance rate by mutation distance; positive-parent/positive-child mutants remain accepted across all distances, whereas destructive controls decline; targeted sub-classes are in Supplementary Fig. S21. (j) Stacked Markov plausibility score by functional class; proton pumps (green) score near 1 while all other subfamilies score near 0, with the τ_90_ threshold (dashed) providing clean separation.

Spectral-tail validation is the most relevant diagnostic for the GA’s blue-shifted-target regime and shows lower accuracy, retained useful triage capacity. The predictor consistently compressed predictions toward the training mean, overestimating *λ*_max_ for blue-shifted variants; a 490 nm prediction corresponded to a measured value bluer than 490 nm by up to 80 nm. SHAP (SHApley exPlanation) analysis identified the retinal-binding pocket as the primary prediction driver (Computational Supplementary Information Figure CS7): the top five residues accounted for > 45% of mean absolute SHAP values, with Ala 124 (as in alignments shown in Computational Supplementary Information Figure CS4) carrying the largest individual contribution (importance = 0.088).

#### Sequence-plausibility filtering

We evaluated four key models: Markov chains, Levenshtein distance, alignment-based, and Chaos Game Representation (CGR) (Figure 3e). For each of these 4 models we considered four data cases: sequence-only, structure-only, stacked (multiplying probabilities), or hybrid (concatenating input data) modes, each representing how primary (sequence) and secondary (structural) data are used. Using extant proteins, tests covered two cases of class separation: (1) rhodopsins (n=884) versus non-rhodopsin bacterial ion transporters of similar length (n=10,000), and (2) proton pumps (n=310) versus phylogenetically related subfamilies (chloride pumps and sensory rhodopsins; n=296). All filters performed well at level (1): Levenshtein sequence achieved median AUROC = 1.000 and F1 = 0.989, while stacked Markov achieved median AUROC = 0.999 and F1 = 0.980) (Figure 3e and f).

At level (2), the stacked Markov model achieved median AUROC = 1.000 and F1 = 0.984 under random sequence-level cross-validation; Levenshtein baselines performed comparably. Structure-only features failed to discriminate subfamilies (Markov structure median AUROC = 0.489, F1 = 0.618), consistent with the conserved secondary-structure fold across microbial rhodopsin subfamilies (Figure 3e and f).

While these results are promising, AUROC on natural sequences does not directly reflect the GA’s operating regime, where the filter scores mutants of seed sequences rather than unseen natural relatives. Thus, to better reflect this operation mode we tested on mutants from wild type sequences. Considering two primary cases: (1) functional (physicohemically similar amino acid substitutions) and (2) disruptive (physicochemically dissimilar mutations). Using the same model and framework, trained solely on known proton pumps, we assessed each mutant category across a range of mutagenesis distances. Functional variants of proton-pump scaffolds remained accepted out to large mutation distances, while disruptive(positive-to-negative) substitutions, negative-parent controls, and non-rhodopsin baselines are rejected even at small distances (Figure 3h and i). Levenshtein and BLAST baselines fail here: synthesised mutants always sit close to their training-set parents, so nearest-neighbour metrics accept them regardless of whether the substitutions are conservative or destructive showing the primary benefit of probabilistic models that learn mutagenesis rules.

A recurrent critical failure of filters is the introduction of systematic bias due to overreliance on homology. Figure 3g demonstrate the low correlation between wavelength and empirical probability from 10-fold CV. Thus, the Markov filter serves to reject most non-functional sequences while preserving the mutational diversity required for directed evolution (Figure 3). Together, the wavelength regressor and sequence-plausibility filter equipped the GA with the scoring and constraint functions required for directed evolution toward blue-shifted targets. A complete breakdown of all filter architectures tested, as well as their failure modes is presented in Computational Supplementary Information Figure CS2.3.

### Genetic algorithm (GA) for the generation of mutants

The GA minimised |*λ̂* _max_ − 490 nm|subject to Markov sequence-plausibility acceptance with priors guided from sequence conservation. Per-generation dynamics are illustrated for a representative single-seed run (Run 789); aggregate statistics and plots over all seeds are in Computational Supplementary. With a mutual information guided prior, and population setting as detailed in methods, said run reached the 490 nm target by generation 5 (sequence with predicted *λ*_max_ ≤ 490). A Markov filter threshold of 0.2 (as outlined with null set behaviour outlined in Computational Supplementary Information Table CS5) limited drift from proton-pump-like sequence space by allowing only threshold-passing candidates to remain in the selectable population.

LDA (Linear Discriminant Analysis) projection onto rhodopsin classifier axes (Figure 4a) shows the accepted best-sequence trajectory remaining within the proton-pump region throughout; filter-rejected candidates scatter toward the boundaries of adjacent classes (channels, chloride pumps, sodium pumps, sensory rhodopsins) but are excluded from the parent pool. The filter bounded this drift (Figure 4d): at generation 5 the best sequence had a Markov score of −2.72 (p≈0.27). Markov scores fell as mutants accumulated substitutions and partially recovered as implausible candidates were rejected. Quantitatively, for this run, the converged best sequence (generation 15, *λ̂*_max_ =489.8 nm) had a minimum Levenshtein distance of 40 to training-set proton pumps; the three closest sensory-annotated sequences (distance 41-42) are bacteriorhodopsin variants with pump-like sequence character, and the nearest genuinely non-pump sequence (channel rhodopsin, distance 210) is 5.3× further, confirming that the accepted mutants remained within the proton-pump sequence neighbourhood.

**Figure 4.**
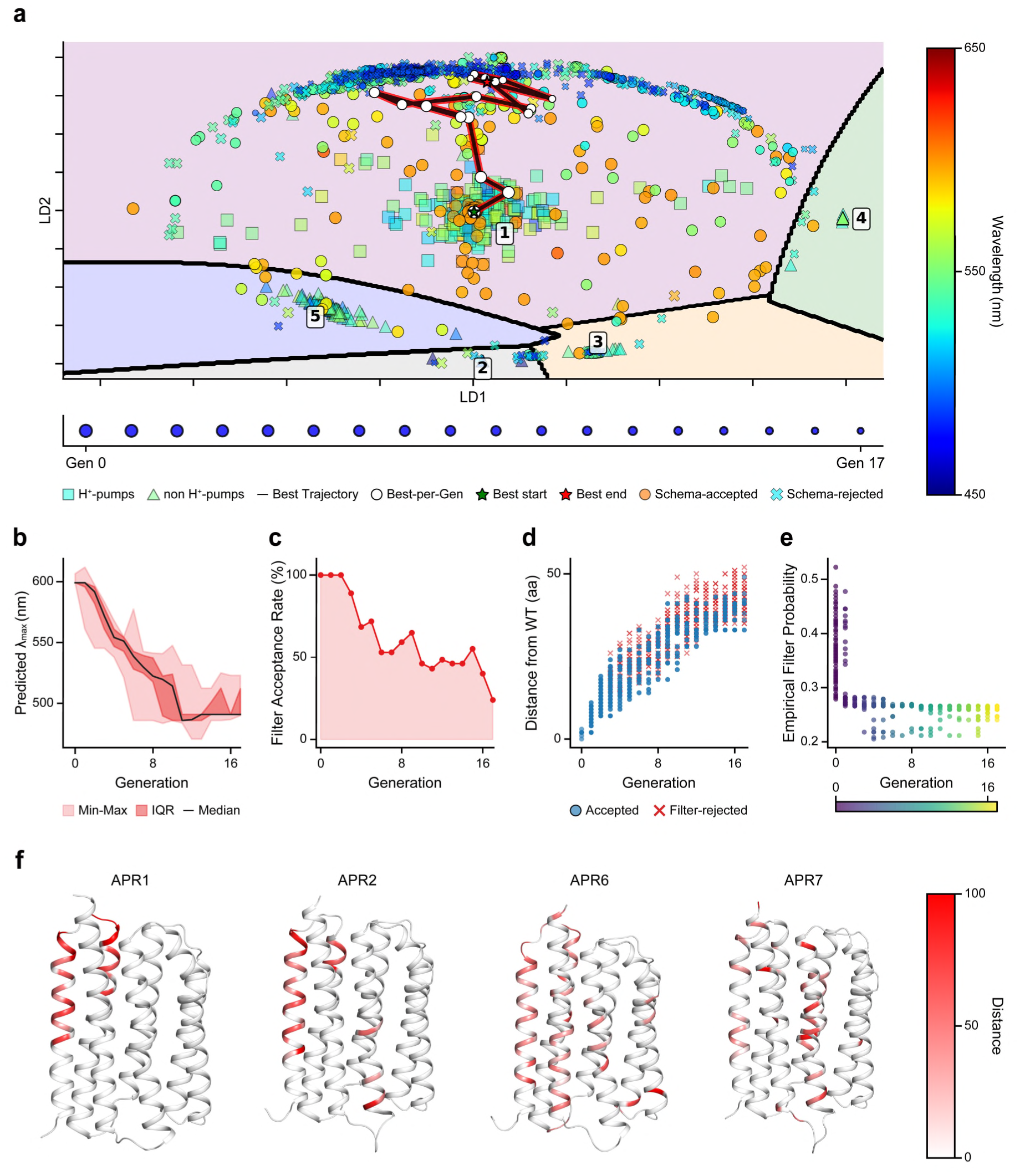
(a) LDA projection with SVC decision regions: 1 = proton pumps, 2 = channels, 3 = chloride pumps, 4 = sodium pumps, 5 = sensory rhodopsins. Circles are accepted variants; crosses are rejected variants. (b) Trace of predicted λ_max_ for children across generations decreases toward the 490 nm target. (c) Filter acceptance rate across generations, (d) Levenstein distance from closest wild type homolog accepted are blue circles, rejected are red crosses (e) Empirical filter probability p; the predicted target is reached at G5 (p≈0.27). (f) AlphaFold 3 structures of APR1, APR2, APR6, and APR7 coloured by per-residue BLOSUM62 distance from the closest wild-type proton pump; substitutions cluster in the retinal-binding pocket and extracellular loops.

Four designs were selected for experimental characterisation. AlphaFold 3 structural models (Figure 4f), coloured by residue-level BLOSUM distance from the closest wild-type proton pump, show that accepted substitutions cluster in the retinal-binding pocket and extracellular loops, consistent with the SHAP-identified prediction drivers.

Ablation experiments varying mutation prior, filter strength, and target wavelength confirmed these dynamics at scale. MI-weighted priors reached the 490 nm target in a median of 7 generations versus 15 for inverse-MI priors; all priors achieved ≥94% success at permissive filter strengths (0.05-0.10). At the hardest 410 nm target, strict thresholds (≥0.30) blocked convergence entirely, while at 490 nm success remained above 90% up to threshold 0.25. Without filtering, 98.8% of runs converged on the scalar wavelength objective but produced designs below the 20th-percentile plausibility threshold-confirming that the Markov filter acts as an in-loop sequence-identity constraint rather than a post-hoc screen. Full hyperparameter interactions and per-prior convergence distributions are in Computational Supplementary Information Figure CS2.4.

### Experimental characterisation of the designed rhodopsins

#### AI-designed rhodopsins are proteorhodopsins absorbing blue light

From filter-passing 490-nm-target candidates across ten seed runs, we selected the seven with the highest Levenshtein distance from the wild-type scaffold. This selection tested retention of distant proton-pump-like sequences by the plausibility filter, not ranking of closely spaced wavelength predictions. AlphaFold 3 structural predictions then ranked these seven by pTM score, and the top five were taken forward for cloning, transformation, and expression. Four of the five were successfully cloned into plasmid pLO11a-GR ^6^, transformed and expressed in *E. coli* and *Cupriavidus necator* H16 (previously *Ralstonia eutropha* H16) for further tests, the fifth design did not yield viable transformants and was not assayed further. The four designed rhodopsins were designated Artificial Proteorhodopsins (APRs): APR1, APR2, APR6, and APR7 (Biological Supplementary Information Table S3).

To infer the spectroscopic and functional characterisations of four AI-designed rhodopsin mutants, their peptide sequences were aligned with representative microbial rhodopsins. Phylogenetic analysis indicates that four mutants belong to Proteorhodopsin (PR) clade (Figure 5a), and the multiple sequence alignment reveals that the key amino acid residues in the APRs correspond to either DTE or DTD motif (Figure 5b). These motifs match up Asp-97, Thr-101, and Glu-108 (DTE motif) in green-absorbing PRs found in the SAR86 group (GPR), and the Asp-85, Thr-89, and Asp-96 (DTD motif) of *Halobacterium salinarum* bacteriorhodopsin (HsBR) ^31^.

**Figure 5.**
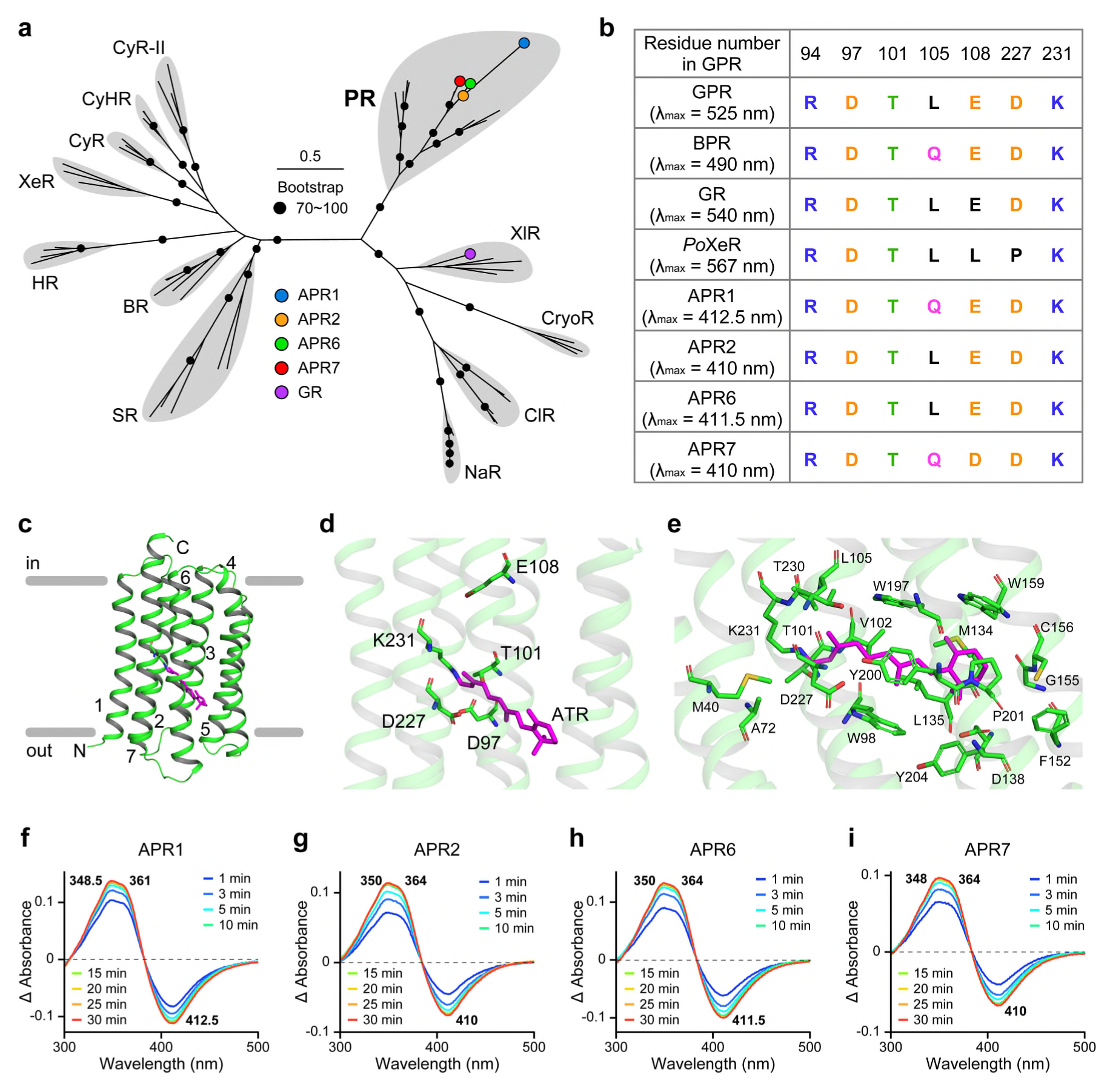
Phylogenetic, predicted structural, and spectroscopic characteristics of APRs. (a) Phylogenetic tree of selected microbial rhodopsins. Bootstrap probabilities (≥ 70%) are indicated by black circles. Abbreviations: CyHR: Cyanobacterial halorhodopsin, CyR: Cyanorhodopsin, CyR-II: Cyanorhodopsin-II, XeR: Xenorhodopsin, SR: Sensory rhodopsin, HR: Halorhodopsin, BR: Bacteriorhodopsin, NaR: Na+-pumping rhodopsin, ClR: Cl--pumping rhodopsin, XlR: Xanthorhodopsin-like rhodopsin, CryoR: CryoRhodopsin, PR: Proteorhodopsin. (b) Key residues for the function of microbial rhodopsins (orange, acidic; blue, basic; black, aromatic; green, -OH bearing; magenta, asparagine and glutamine residues; gray, leucine residue). (c-e) The predicted structures of APR6, the highest predicted Template Modeling scores among all APRs (0.85). A stick model of all-trans retinal (ATR) is shown in magenta. Structural predictions were conducted using AlphaFold 3. (f-i) Time-series absorbance changes of membrane fractions containing APR1, APR2, APR6, and APR7, caused by incubation with hydroxylamine, respectively.

Alphafold structural modelling predicts that the APRs have seven transmembrane helix architectures (Figure 5c and Biological Supplementary Information Figure BS1), and form retinal binding pockets close to the conserved lysine residue (Lys-231 in GPR), which covalently binds to retinal chromophore (Figure 5d and 5e), similar to other proton-pumping rhodopsins ^32, 33, 34^. In PR and BR, two conserved negatively charged residues, equivalent to Asp-97 and Asp-227 in GPR, generate a strong electrostatic attraction and lead to outward proton transport ^35, 36^, consistent with the mechanism of outward proton-pumping rhodopsins. The colour tuning in PRs is strongly influenced by the residue at position 105 of GPR; where leucine is associated with green-absorption (λ_max_: 525 nm) and glutamine with blue-absorption (λ_max_: 490 nm) ^11^. Based on the sequence feature, it was predicted that APR1 and APR7 are blue-absorbing, and APR2 and APR6 are green-absorbing (Figure 5b).

We analysed the maximum absorption wavelength of rhodopsin by monitoring the change of absorbance spectrum before and after bleaching with hydroxylamine (HA). HA hydrolyses the Schiff base bond, which connects the retinal chromophore to the lysine residue ^20, 21, 37^. The membrane fractions from rhodopsin-expressed *Escherichia coli* cells were incubated with HA, resulting in distinct absorption spectra with a negative peak around 410 nm. The λ_max_ of APR1, APR2, APR6, and APR7 were 412.5, 410, 411.5, and 410 nm, respectively (Figure 5f-i). Concomitant with these negative peaks, the positive peaks at 350-370 nm appeared, indicating the formation of the retinal oxime. These data suggest that all APRs bound the retinal chromophore and absorbed violet-purple light region. Notably, APR2 and APR6, which were predicted to be green-absorbing rhodopsins based on peptide sequence features, also shows blue-shifted absorption (Figure 5).

#### Rhodopsin-mediated phototrophic growth in engineered *Cupriavidus necator*

All APR variants cloned in *Cupriavidus necator* were induced by L-arabinose. Due to the blue shift in absorption of APRs, induced APR-expressing cells displayed a distinctly yellow-orange colouration, contrast to green light absorbing GR-containing cells, which presented a pink colour ^6, 8^ (Biological Supplementary Information Figure BS2).

Similar to the previous finding that *Gloeobacter* rhodopsin (GR) expression enhanced the growth of *C. necator* strain under the green light ^38^, we assessed whether AI-designed APRs also confer a growth advantages to APR-expressing *C. necator* variants using formate as the sole carbon source under blue light. APR-expressing *C. necator* variants (Biological Supplementary Information Table S3) were grown in batch culture using Chi.bio™ reactors, enabling continuous OD₆₀₀ monitoring under defined illumination conditions. All cultures were normalised to OD_600_ = 0.1 at the start of the experiment and initially grown in darkness for the first 8 h. Before assessing strains under illumination, the growth profiles of induced cultures in the dark were first examined across all strains (Biological Supplementary Information Figure BS3). As growth profiles were comparable in the absence of light, the pooled dark-induced condition was compared with uninduced controls grown under light or dark conditions (Biological Supplementary Information Figure BS4). These controls follow similar trajectories throughout the experiment, with only modest divergence at later time points, which was small relative to the effects subsequently observed in light-exposed strains. The dark-induced condition was therefore used as the principal controls for subsequent comparisons.

Compared with the dark controls, divergence between APR-expressing *C. necator* variants emerged after approximately 20-30 h (Figure 6a). Growth trajectories were compared using an ordinary least squares (OLS) regression model incorporating time (linear and quadratic terms) and strain × time interactions. This analysis revealed a significant difference in growth kinetics between strains (likelihood ratio test, *p =* 5.18 × 10⁻⁸). Relative to the dark-induced control, Figure 6a shows that all APR strains under blue light exhibited significantly steeper increases in OD_600_ over time (APR1: *β* = 0.0168, *p* = 2.12 × 10⁻¹⁸; APR2: *β* = 0.0162, *p* = 9.50 × 10⁻¹²; APR6: *β* = 0.0110, *p* = 4.39 × 10⁻¹⁰; APR7: *β* = 0.0153, *p* = 1.08 × 10⁻¹⁵). In contrast, GR expressing *C. necator* also had a significant but smaller increase in growth rate (*β* = 0.0058, *p* = 0.005) (Figure 6a), indicating a weaker growth promoting effect relative to the APR variants. No major differences were observed in the curvature of the growth trajectories, as indicated by the absence of significant strain × time² interaction terms (Biological Supplementary Information Table S4 and S5). In contrast, significant strain × time interaction terms for all strains indicate that the observed differences were driven primarily by variation in exponential-phase growth rate.

**Figure 6.**
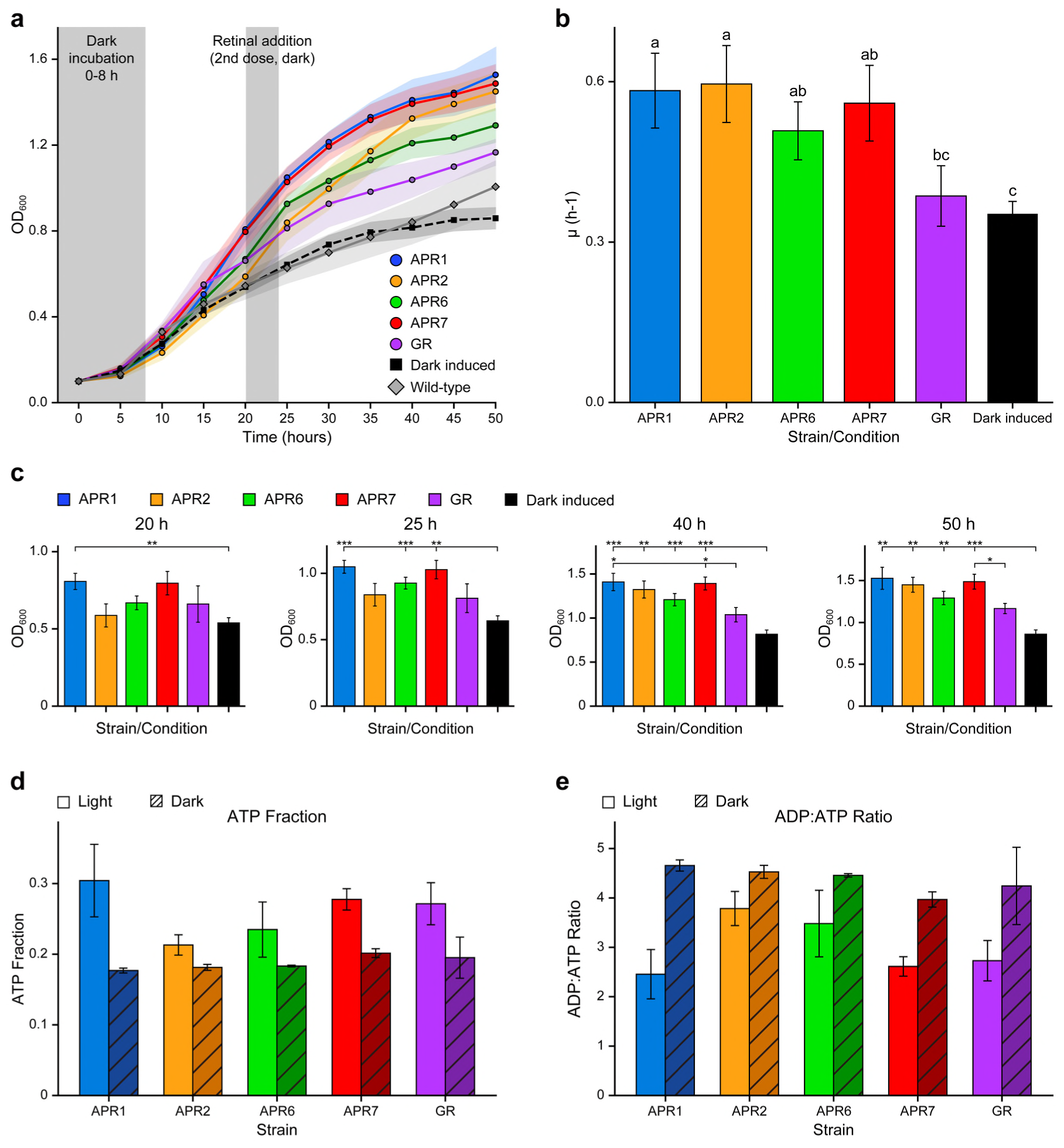
Biological validation of the designed APR functions for enhanced growth of *C. necator* H16 under blue light. (a) Growth of induced light-exposed strains relative to the pooled dark controls. Mean OD_600_ ± SEM for induced light-exposed cultures expressing APR1 (n=10), APR2 (n=6), APR6 (n=12), APR7 (n=10), or GR (n=8), shown together with the pooled dark control (n=52) and Wild-type combined uninduced control (n=14). Grey shaded regions indicate periods of dark incubation. The second retinal addition is also indicated by the vertical shaded region. (b) Exponential-phase growth rate across strains. Average specific growth rate (μ, h⁻¹) during the exponential phase (10–40 h) for each induced light strain and the dark induced control, shown as mean ± SEM (n = 6–12 biological replicates per strain; n = 52 pooled dark induced control). Different letters indicate statistically significant differences between groups (Welch one-way ANOVA followed by pairwise Welch t-tests with Benjamini–Hochberg correction, p < 0.05). (c) Time-point snapshots of strain-specific OD_600_. Mean OD_600_ by strain at 20 h, 25 h, 40 h and 50 h. These snapshots illustrate the emergence and persistence of differences observed in the full growth-curve analysis, *** = p<0.001, ** = p<0.01, * = p<0.05. Bars represent mean ± SEM (n = 6–12 biological replicates per strain; n = 52 pooled control measurements). (d) ATP fractions of APR expressing *C. necator* H16 in the light is higher than those in the dark, calculating as ATP / (ATP + ADP), for each strain under the indicated conditions. (e) ADP/ATP ratio for the same samples of (d). Data are shown for induced light and induced dark cultures after 20 hours, with values normalised per plate. Bars represent mean ± SEM across biological replicates (n = 3 plates), with Light and Dark conditions paired within each plate.

To determine the timing of strains divergence, pairwise comparisons at each time point were performed using Welch’s two-sample t-tests, with p-values adjusted using the Benjamini-Hochberg procedure to control the false discovery rate. No significant differences were detected during early growth before 20 h, indicating similar initial behaviour across all strains. At 20 h, APR1 was significantly higher than the dark-induced control (*p* = 0.00653) (Figure 6b and Biological Supplementary Information Table S4). By 25 h, APR1, APR6, and APR7 were significantly higher than the dark-induced control (*p* = 1.85 × 10⁻⁵, 2.47 × 10⁻⁴, and 1.01 × 10⁻³, respectively), whereas APR2 and GR did not differ significantly at this stage (Figure 6b). By 40 h, this pattern had broadened: all APR variants were now significantly elevated relative to the dark-induced control (APR1: *p* = 5.63 × 10⁻⁴; APR2: *p* = 6.26 × 10⁻³; APR6: *p* = 5.63 × 10⁻⁴; APR7: *p* = 5.25 × 10⁻⁵), and APR1 and APR7 were also significantly higher than GR (*p* = 0.0259 and *p* = 0.0166, respectively), indicating a greater growth advantage for these variants during mid-to-late growth (Figure 6c). These differences were maintained at 50 h, where all APR strains remained significantly above the control (APR1: *p* = 1.87 × 10⁻³; APR2: *p* = 1.55 × 10⁻³; APR6: *p* = 1.16 × 10⁻³; APR7: 2.47 × 10⁻⁴). APR7 also continued to exceed GR (*p* = 0.0232), while other APR-GR comparisons were not significant after correction (Figure 6c and Biological Supplementary Information Table S5).

#### Growth-rate analysis showed a stronger and more sustained light-dependent kinetic advantage in APR expressed mutants

Time-resolved growth rates (μ) were calculated to examine how growth dynamics evolved over the 50 h experiment (Biological Supplementary Information Figure BS5). All strains exhibited an initial increase in growth rate during the early exponential phase, which peaked between 10-15 h and declined thereafter as cultures approached the stationary phase. Under illumination, APR-expressing strains consistently exhibited higher growth rates than the dark-induced control during the early-to-mid exponential phase (10-25 h). For example, APR1 reached a peak growth rate of *μ* = 0.139 ± 0.013 h⁻¹ at 15 h under light, compared with *μ* = 0.085 ± 0.007 h⁻¹ under dark conditions at the same time point. Similar illumination-dependent increases were observed for APR7 (*μ* = 0.136 ± 0.023 h⁻¹ under light vs *μ* = 0.061 ± 0.011 h⁻¹ under dark at 15 h) and APR6, which reached *μ* = 0.132 ± 0.013 h⁻¹ at 10 h under light compared with *μ* = 0.119 ± 0.010 h⁻¹ under dark. APR2 showed a more modest but still detectable light-dependent effect, with peak growth rates of *μ* = 0.118 ± 0.019 h⁻¹ at 10-15 h but sustained higher values under illumination at later time points (*μ* = 0.073 ± 0.012 h⁻¹ at 25 h under light vs *μ* = 0.037 ± 0.004 h⁻¹ under dark). In contrast, GR displayed a comparatively weak and inconsistent response to illumination. Although GR reached a slightly higher peak under light (*μ* = 0.148 ± 0.020 h⁻¹ at 10 h) than under dark (*μ* = 0.119 ± 0.010 h⁻¹), this difference was not sustained, and growth-rate profiles converged thereafter (*μ* ≈ 0.041-0.052 h⁻¹ at 20-25 h under light vs *μ* ≈ 0.037-0.044 h⁻¹ under dark). Growth rates were therefore averaged across the exponential phase (10-40 h) to provide a summary measure for each strain (Fig. 5). Mean growth rates were highest for APR2 (*μ* = 0.0596 ± 0.0072 h⁻¹), APR1 (*μ* = 0.0583 ± 0.0070 h⁻¹), and APR7 (*μ* = 0.0560 ± 0.0071 h⁻¹), differing by less than 0.004 h⁻¹. APR6 exhibited a lower mean growth rate (*μ* = 0.0508 ± 0.0054 h⁻¹), while GR showed the lowest growth rate among the light-exposed strains (*μ* = 0.0386 ± 0.0057 h⁻¹).

These differences were statistically significant (Welch one-way ANOVA, *F₅,₂₂.₇* = 6.14, *p* = 0.001). Pairwise Welch t-tests followed by Benjamini-Hochberg correction showed that all APR strains exhibited higher growth rates than the dark-induced control (adjusted *p* < 0.05), whereas GR did not differ significantly from the dark-induced condition (Biological Supplementary Information Table S4 and S5). In addition, GR was lower than APR1 and APR2 (adjusted *p* < 0.05), while differences among APR variants were not significant.

#### APR strains exhibit an improved cellular energy state under illumination

To determine whether the enhanced growth observed in APR strains was associated with changes in cellular energy status, ATP and ADP levels were quantified and expressed as ATP fraction and ADP/ATP ratio (Figure 6d and 6e). Illumination significantly altered both measures of cellular energy status across all strains. ATP fraction increased from 0.186 ± 0.004 under dark conditions to 0.252 ± 0.015 under illumination (two-way linear model, main effect of condition: *F₁,_19_* = 18.49, *p* = 0.0004), while the ADP/ATP ratio decreased from 4.42 ± 0.11 to 3.16 ± 0.24 (*F_1_,_19_* = 25.62, *p* = 0.0001). Although all strains exhibited this light-dependent effect, the magnitude of the response did not differ significantly between strains (Genotype × Condition interaction: *F₄,_19_* = 1.28, *p* = 0.314 for ATP fraction; *F₄,_19_* = 1.18, *p* = 0.351 for ADP/ATP ratio). Within-strain comparisons between light and dark conditions were not statistically significant (Welch two-sample t-tests with Benjamini-Hochberg correction, all adjusted p > 0.05). Taken together, these results demonstrate a consistent light-dependent growth advantage in APR-expressing strains, accompanied by a shift toward a more energy-rich cellular state.

## Discussion

This study demonstrates that an integrated AI-guided design pipeline can generate non-natural microbial rhodopsins with strongly blue-shifted features while retaining key sequence signatures of proton-pumping rhodopsins. The experimental validation of APR1, APR2, APR6, and APR7 represents a meaningful advance in AI-guided protein design. It demonstrates extrapolative rather than interpolative design in a functionally constrained protein family where spectral tuning and proton pumping feature are typically phylogenetically coupled. Natural proton-pumping rhodopsins are largely confined to the green-orange spectral window, a bias reflecting evolutionary history rather than physical necessity. The ∼410 nm absorption of the APR variants lies well outside the characterised proton-pump spectral distribution and has not yet been found in nature (Figure 5). This work establishes that machine-learning-guided genetic algorithms can potentially navigate into genuinely novel regions of protein sequence space without sacrificing the target biological function.

By combining a genetic algorithm for sequence generation, a stacked LASSO/XGBoost regressor for spectral prediction and fitness ranking, and a Markov-based plausibility filter for proton-pump-like sequence, the pipeline explored regions of rhodopsin space that are difficult to access by conventional rational design or mutagenesis (Figure 1). The resulting APR variants contained 26–50 substitutions relative to their closest relative in the model training set, yet retained the conserved proton-relay motif and retinal-binding lysine, indicating that extensive sequence diversification can be compatible with preservation of core rhodopsin architecture. In the absence of a sequence plausibility constraint, the genetic algorithm tended to move sequences away from the proton-pump clade towards blue-shifted but functionally distinct rhodopsin families. The Markov filter therefore played an essential role by limiting this drift and maintaining proton-pump-like sequence statistics. This constraint was not simply a phylogenetic filter: it helped preserve sequence features associated with ion specificity and proton-pumping architecture while allowing spectral exploration beyond the densely sampled natural proton-pump region.

The biological validation in *C. necator* H16 indicates that the AI-designed APRs are not only spectroscopically altered but also biologically active in cells. Figure 5 shows that all APRs are blue-light activated proton pumps. Under defined blue-light illumination, these APR-expressing *C. necator* H16 strains show enhanced growth relative to the dark-induced control, and some APR variants outperformed the established GR-based system (Figure 6). The increase in cellular ATP fraction supports a model in which APR-mediated light capture contributes to proton-motive-force generation and ATP synthesis (Figure 6).

These findings advance rhodopsin engineering in two ways. First, they show that AI-guided sequence exploration can move beyond natural spectral distributions while preserving a conserved membrane-protein fold. Previous ML-guided rhodopsin engineering has largely operated within or near the natural spectral range, where training data are dense and prediction errors are smaller. Here, the model was deliberately used at the edge of the training distribution, where extrapolation is difficult but potentially more interesting. Second, the work illustrates the importance of combining performance-driven optimisation with biological plausibility constraints. Spectral prediction alone drove sequences towards non-proton-pump lineages, whereas the Markov filter allowed the search to remain within a proton-pump-compatible region of sequence space.

Several limitations remain. The hydroxylamine bleaching assay provides evidence of retinal incorporation and retinal-associated absorbance changes, but it does not directly measure proton-pumping rate, photocycle kinetics or quantum yield. The current predictor provides coarse directionality rather than precise wavelength control, and monotonic ranking among closely spaced predictions has not yet been established. The relationship between membrane-fraction spectra, purified-protein spectra and in vivo phototrophic performance also remains unresolved. Future work should therefore combine action spectroscopy, native-membrane time-resolved measurements, direct proton-transport assays and QM/MM-based modelling of retinal excited states.

Overall, this study establishes a generalisable strategy for AI-guided protein engineering: use generative search to explore sequence space, predictive models to enrich for desired phenotypes, and structural or evolutionary plausibility filters to preserve biological function. Applied to microbial rhodopsins, this strategy generated artificial proteorhodopsins with strongly blue-shifted retinal-associated spectral features and light-dependent growth-promoting activity in *C. necator*. These results provide a foundation for programmable rhodopsin-based light-harvesting modules and future artificial photosynthesis systems.

## Methods

### Computational pipeline

All analyses used Python 3.10.12 on Ubuntu 22.04 LTS workstations. Data analysis used NumPy (1.23.5), SciPy (1.10.1), scikit-learn (1.2.2), XGBoost (1.7.5), NetworkX (3.0), and igraph (0.10.4) (Matplotlib 3.7.1, Seaborn 0.12.2 for visualisation). Jupyter notebooks, container definitions (Singularity/Docker), and fixed random seeds (seed=42; the same seed was applied independently to the GA, ML training, and train/test split steps) are described in the Data Availability section below. Full details of sequence curation, phylogenetic analysis, wavelength prediction, Markov filtering, and genetic algorithm design are provided in the Supplementary Methods; the key specifications are summarised below.

#### Sequences, Alignment, and Features

The curated dataset comprises 884 microbial rhodopsin sequences with functional annotation and measured; all 884 carry wavelength labels and were used for regressor training and cross-validation. Sequences were aligned with MAFFT (L-INS-i) against a curated rhodopsin reference. Gaps were retained as a 21st character, giving a final alignment length of *L* = 555 residues. For the regressor, each aligned residue was encoded with an 18-component Feature Map: five continuous physicochemical descriptors (Kyte-Doolittle hydrophobicity, side-chain volume, Grantham polarity, isoelectric point, Chou-Fasman helical propensity), each standardised to zero mean and unit variance over the alignment, plus 13 binary indicator channels (chemical class, charge sign, aromaticity, gap). One-hot and rank-ordinal encodings were retained for comparison.

λ_max_ Regressor

The ML objective is supervised prediction of experimentally measured *λ*_max_ from aligned sequence features. The predictor stacks an L1-penalised LASSO with an XGBoost residual learner. LASSO *α* was tuned over a logarithmic grid *α*∈[10^−4^,10^1^] (50 points) by inner 5-fold CV; XGBoost hyperparameters: max_depth= 6, n_estimators= 300, learning_rate= 0.05, subsample= 0.8, colsample_bytree= 0.8. The LASSO prediction was concatenated with the original feature vector and passed to XGBoost; both stages were refit on the same outer fold to avoid leakage. We report shuffled 10-fold CV as the primary metric because the downstream GA operates within the proton-pump clade enforced by the plausibility filter; leave-family CV and leave-wildtype CV are reported as out-of-distribution diagnostics. SHAP values were computed with the TreeExplainer on the XGBoost residual learner.

#### Markov Sequence-Plausibility Filter

The filter used is a position-aware first-order Markov (bigram) model fit on the annotated proton-pump subfamily, stacking aligned-sequence transitions with secondary-structure-context transitions and combining the two log-likelihoods by mean. Transition counts were Laplace-smoothed (pseudocount = 1) to handle unseen bigrams. The filter score is the average log-likelihood per aligned position. A candidate is accepted if its score exceeds a threshold *τ* set at the *q*-th percentile of training-set scores; *q* is the “filter strength” reported in ablations. This threshold defines the GA’s feasible set: among the finite set of aligned sequences of length *L* = 555 over the 21-character residue/gap alphabet, only candidates with Markov score ≥ *τ* can contribute to subsequent generations. Production runs used *q* = 0.20 (approximately one standard deviation below the mean training score; chosen from the F1/TPR–FPR calibration sweep). Candidates failing the filter are generated and scored within a generation but excluded from the parent pool for the next generation; if fewer than *n*_parents_ filter-passing children survive, the deficit is back-filled with the best filter-passing parents from the previous generation.

#### Genetic Algorithm

Each generation maintains a population of *P* = 50 sequences. The top *n*_parents_ = 15 by fitness are selected by truncation selection, and *n*_elite_ = 5 elite parents are copied unchanged into the next generation. The remaining *P* − *n*_elite_ = 45 slots are filled by crossover-then-mutation: a single key parent is drawn uniformly from the parent pool, the number of crossover splice events is Poisson(*λ*_xo_ = 1.0), and each splice segment length is drawn from a truncated normal over alignment positions (mean *μ*_seg_ = 6, std *σ*_seg_ = 6, truncated at [1, *L*]). Each child is then mutated: the per-sequence number of mutations is Poisson(*η* = 0.01 ⋅ *L*) and mutation sites are sampled from a position-wise prior *π*, with substituted residue drawn uniformly from the 20 amino acids plus gap. We compare seven priors *π*: uniform, sequence entropy, mutual information with *λ*_max_ (MI), inverse-entropy, inverse-MI, regressor-importance, and inverse-regressor. The regressor-importance prior is a normalised combination of LASSO coefficient magnitudes and per-position SHAP values from the XGBoost stage. All non-uniform priors are blended with a 10% uniform floor (*π* ← 0.9*π* + 0.1/*L*) so every position retains non-zero mutation probability. The optimisation objective is

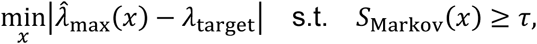

implemented as fitness = −|*λ̂*_max_ − *λ*_target_| for filter-passing candidates; sequences failing the filter are assigned the worst fitness and cannot seed later generations. Termination is by early stopping when the best fitness fails to improve by more than *ε* = 0.01 for 5 consecutive generations, or by a generation cap of *G* = 40 (hyperparameter ablation), *G* = 120 (prior-vs-filter schema test), and *G* = 120 (production runs). Each production run uses an independent random seed; ten seeds were run for the 490 nm target.

### Selection of designed rhodopsins for blue light harvesting

The genetic algorithm generated candidates targeting *λ*_max_= 490 nm. Candidates with Markov log-likelihood below the filter threshold were rejected before ranking. From ten independent GA runs, each with a different random seed, we selected the seven filter-passing candidates with the greatest Levenshtein distance from the wild-type scaffold. AlphaFold 3 (https://alphafoldserver.com) predictions (default model, three seeds per sequence, prediction with retinal cofactor disabled) were then used to rank these seven by pTM score; the top five (pTM 0.78) were taken forward experimentally and the remaining two were discarded. Four designs were successfully cloned, transformed, and expressed in *C. necator* (APR1, APR2, APR6, and APR7).

Three-dimensional structures of four microbial rhodopsin (APR) variants, APR1, APR2, APR6, and APR7 were modeled as homopentamers using the folding entry on the biodesign platform (https://biodesign.top). Each variant was represented as an oligomer of five identical polypeptide chains. Prior to the inference, MSAs were generated independently for each single-chain sequence. All four APR proteins belong to the rhodopsin family and contain a conserved lysine that forms a Schiff base with all-trans-retinal. In the input, this post-translational modification (ptm) was specified using the Chemical Component Dictionary (CCD) code LYR (N6-(all-trans-retinyl)-L-lysine, i.e., retinylated lysine). Modification sites were assigned to the lysine (K) within the conserved “VNK” motif near the C-terminal transmembrane region (1-based indexing): APR1 K234, APR2 K231, and APR7 K233. APR6 carries “LDVTTK” rather than the VNK motif at the homologous position but retains a lysine at residue 231; LYR was therefore assigned at this homologous site. Each polypeptide chain carried one LYR modification, yielding five LYR residues per homopentamer. We used Prodenovo (https://prodenovo.ai/protein-design) to visualise the predicted rhodopsin structures and *in-silico* analyse the protein features.

### Bacterial strains, culture conditions and plasmid construction

All bacterial strains, plasmids, and primers used in this study are shown in Biological Supplementary Information Table S3. *C. necator* cultures were grown overnight in Lysogeny Broth (LB) supplemented with 15 µg mL⁻¹ tetracycline under shaking conditions (250 rpm) at 30 °C. Overnight cultures were diluted 1:10 into fresh LB medium containing 80 mM sodium formate, 5 µg mL⁻¹ all-trans-retinal, and 0.2% (w/v) L-arabinose to induce rhodopsin expression. Cells were incubated overnight under the same shaking and temperature conditions.

Cells were centrifuged at 2800 x g for 10 min, washed three times with minimal medium (composition below), and resuspended in 10 mL of the same medium. The minimal medium (per litre of distilled water) contained: 6.8 g Na₂HPO₄, 1.5 g KH₂PO₄, 1.0 g (NH₄)₂SO₄, 0.2 g MgSO₄·7H₂O, 0.02 g FeSO₄·7H₂O, 4 mg CaSO₄·2H₂O, and 0.1 mg thiamine hydrochloride (C₁₂H₁₈Cl₂N₄OS). Phosphate and ammonium salts were sterilised by autoclaving, while magnesium, iron, calcium, and thiamine components were sterilised by filtration. A 100x trace element solution (0.1 mL L⁻¹ final concentration) was added, prepared as follows: 10% (v/v) 5 M HCl containing 15 g FeCl₂·4H₂O, 1.9 g CoCl₂·6H₂O, 1.0 g MnCl₂·4H₂O, 0.7 g ZnCl₂, 0.62 g H₃BO₃, 0.36 g Na₂MoO₄·2H₂O, and 0.16 g CuSO₄ per litre.

### *C. necator* growth assays

Cells were diluted 1:20 in the supplemented minimal medium, with and without L-arabinose, and incubated overnight at their respective growth temperatures. Cultures were normalised to an optical density of 0.1 (OD₆₀₀) in fresh medium prior to growth assays.

Growth experiments were conducted in Chi.Bio™ bioreactors ^39^ for 50 h under light or dark conditions. For APR strains, 457 nm LEDs were activated after 8 h at a power level of 0.01; for GR strains, 523 nm LEDs were used at a power level of 0.032. After 20 h, a second dose of 5 µg/mL all-trans-retinal was added and grown for 4 h without light. These dark incubations were conducted to avoid light-degradation of retinal and allow cells to incorporate retinal into rhodopsin before light-activation.

Four designed rhodopsins were codon-optimised for Cupriavidus necator and cloned into pLO11 using NEBuilder HiFi DNA Assembly. Plasmids were introduced into C. necator H16 by electroporation (2.5 kV). Cultures were grown at 30 ◦C in LB with tetracycline, then diluted into medium containing 80 mM sodium formate, 5 µg mL−1 all-trans retinal, and 0.2% L-arabinose. Cells were washed into minimal medium and normalised to OD600 = 0.1 before growth assays. Growth assays were run for 50 h in Chi.Bio reactors; APR strains were illuminated with 457 nm LEDs after 8 h, GR controls were illuminated with 523 nm LEDs, and OD600 was monitored continuously. The APR wave-length was chosen from the available Chi.Bio LEDs as the shortest wavelength that supported stable assay operation without measurable chromophore photobleaching over the growth window; GR was illuminated with the available green LED nearest its established operating range.

### ATP/ADP measurements

Cellular energy state was assessed by quantifying ATP and ADP levels using the MAK135 ATP/ADP Assay Kit (Sigma-Aldrich), following the manufacturer’s instructions. For each measurement, 10 µL of culture was taken directly from the Chi.Bio reactors™ after 20 h of growth under the specified light or dark conditions.

Samples were prepared for sequential ATP and ADP measurements using the enzymatic conversion steps provided in the kit. Luminescence was measured using a Tecan microplate reader in white 96-well CELLSTAR® plates (Greiner Bio-One). Results were expressed as ATP fraction (ATP / (ATP + ADP)) and ADP/ATP ratio to provide complementary measures of cellular energy state. All measurements were performed on biological replicates (n = 3 plates), with light and dark conditions paired within each replicate. An internal no cell control using the same growth media was included per plate to measure luminescence drift across plates.

### Statistical analysis

Growth data were analysed using complementary approaches to characterise trajectory differences, time-point divergence, and growth rate kinetics. Full OD₆₀₀ time-course data were analysed using an ordinary least squares (OLS) regression model incorporating time as both linear and quadratic terms, with strain included as a categorical variable and interaction terms with time (OD₆₀₀ ∼ time + time² + strain + strain × time + strain × time²). This model enabled comparison of growth trajectories across strains while accounting for non-linear growth dynamics. Model comparisons were performed using likelihood ratio tests.

To identify time points at which strains diverged, pairwise comparisons between groups were conducted independently at selected time points (20 h, 25 h, 40 h, and 50 h) using Welch’s two-sample t-tests, which do not assume equal variances. P-values were adjusted within each time point using the Benjamini-Hochberg procedure to control the false discovery rate.

Specific growth rates (*μ,* h⁻¹) were calculated for each replicate between consecutive timepoints using μ = ln(OD₂/OD₁) / (t₂ − t₁), excluding intervals where OD values were zero or negative. Time-resolved growth rates were summarised as mean ± SEM across replicates at each interval. To compare growth rates between strains, *μ* values were averaged across the exponential phase (10-40 h) for each replicate, and these replicate means were compared using Welch’s one-way ANOVA, followed by pairwise Welch t-tests with Benjamini-Hochberg correction.

ATP fraction and ADP/ATP ratio were analysed using a two-way linear model with genotype and condition (light vs dark) as fixed factors, including a genotype × condition interaction. Statistical significance of model terms was assessed by Type II ANOVA. Within-strain comparisons between light and dark conditions were additionally performed using Welch t-tests.

All statistical analyses were performed in Python using SciPy and Statsmodels. Statistical significance was defined as an adjusted p-value < 0.05.

### Spectroscopic characterisation of APR rhodopsins

Rhodopsin-expressing *E. coli* cells were resuspended in 7 mL of buffer containing 50 mM Tris-HCl pH 8.0, and 500 mM NaCl, and disrupted by sonication (Branson SFX 250 Digital Sonifier, Branson Ultrasonics) on ice-cold water for 5 min. Crude membranes were collected by ultracentrifugation at 106,800 × g for 30 min at 4 °C (Optima XPN-90 Ultracentrifuge with a SW 32Ti rotor, Beckman Coulter) and solubilised with 1% (w/v) *n*-dodecyl-β-D-maltoside (β-DDM; Dojindo Lab). The solubilised membrane fraction containing rhodopsin protein was obtained by ultracentrifugation at 106,800 × g for 30 min at 4 °C (Optima XPN-90 Ultracentrifuge with a SW 32Ti rotor, Beckman Coulter). Dissolved rhodopsin proteins were bleached with 50 mM hydroxylammonium chloride (Sigma-Aldrich) in the dark. The absorption changes were monitored using a UV-2600 spectrophotometer (Shimadzu) at room temperature.

### Phylogenetic analysis and structural prediction

For the phylogenetic analysis of rhodopsins, amino acid sequences of 48 microbial rhodopsins^40^ were collected from the National Center for Biotechnology Information and aligned using MAFFT version 7.515 with EINSI strategy^41^. The phylogenetic tree was inferred using RAxML version 8.2.12 with the PROTGAMMALGF model using 1,000 rapid bootstrap searches^42^. Model selection was performed with the ProteinModelSelection.pl script in the RAxML package. The phylogenetic trees were visualised with Interactive Tree of Life version 6.8.1^43^. The tertiary structures of rhodopsins were predicted using the AlphaFold3 web server with default parameters^44^. Molecular graphics were generated using PyMOL version 2.5.2, and the structure surfaces were analysed using APBS electrostatics into PyMOL package^45^.

## Supporting information

Computational Supplementary Information

Biological Supplementary Information

## Data Availability Statement

The data generated in this study are provided in the main text and/or Supplementary information, where the source data are listed in the Source Data file. Source data are provided with this paper.

## Acknowledgements

W.E.H. acknowledges financial support from EPSRC (EP/M002403/1 and EP/N009746/1) and BBSRC UK-Japan for engineering biology (ISPF-234, UKRI251). W.E.H gratefully acknowledges funding. EPSRC (EP/M02833X/1) for instrumentation. We thank Prof Kwang-Hwan Jung at Sogang University for providing plasmids of GR. We also thank Oliver Lenz at Technische Universität Berlin, Germany for providing wild-type *Ralstonia eutropha* H16 and cloning plasmids. We also acknowledge the CPU computational support provided by Boba Cloud Inc in the USA.

## Author Contributions Statement

H.S. and W.E.H conceived the original idea; H.S., M.J.L., T.F. and W.E.H. designed research; H.S. and A.Y. performed machine learning and biocomputation; M.J.L., T.F. K. M., S. Y., and K.I. carried out biological experiments; J.H., K.I. A.Y.,T.P., Y.W. contributed new reagents/analytic tools; H.S. M.J.L. T.F., K. M., K.I. and W.E.H. analysed data; H.S., M.J.L., T.F. and W.E.H. drafted the manuscript, and all authors revised the manuscript.

