## Supplementary material for "AI-enabled rhodopsin design for blue-light enhanced bacterial growth": Computational Supplementary Information

AI-enabled rhodopsins designed for enhanced light dependent bacterial growth

**Scope of this document.** This document consolidates the computational supplementary methods and results for the above paper. It covers the sequence-landscape analysis, wavelength predictor, sequence-plausibility filter, genetic algorithm, and GA ablation study. Experimental methods and results (spectroscopic characterization, *C. necator* expression, growth and ATP assays) are reported separately in the main manuscript.

**3 Computational Supplementary Tables** **61**

### 1 Computational Supplementary Methods

The main Methods section summarizes the computational environment and data availability. Detailed methods for each computational analysis are provided below to keep the main narrative focused while preserving the full computational record.

#### 1.1 Phylogenetic Analysis

**Dataset curation and preprocessing.** The primary dataset of microbial rhodopsin sequences and experimentally determined absorption maxima ($\lambda_{\max}$) was derived from Inoue et al. (2021)[1]. That study provides family characterization and functional annotations (Proton Pump, Sodium Pump, Chloride Pump, Sensory Rhodopsin, Channelrhodopsin) based on phylogenetic reconstruction and motif analysis. To address the scarcity of blue-shifted variants, we consolidated three channel subtypes—Anion Channel ($n = 6$), Proton Channel ($n = 10$), and Cation Channel ($n = 21$)—into a single "Channel" superclass ($n = 37$). Quality control excluded sequences containing ambiguous amino-acid codes or with length deviations greater than 20% of the canonical bacteriorhodopsin length ($\tilde{2}50$ residues).

**Topological and phylogenetic analysis.** To map the sequence landscape without imposing an alignment, we used the alignment-free Levenshtein edit distance [2]. For any two sequences $a, b \in \mathcal{S}$, the Levenshtein distance $d_{\text{Lev}}(a, b)$ is the minimum number of single-character insertions ($I$), deletions ($D$) and substitutions ($S$) required to transform $a$ into $b$:

$$d_{\text{Lev}}(a, b) = \min\{\, |I| + |D| + |S| \,\}. \tag{1}$$

**Sequence and structural identity assessment.** To quantify the conservation of the proton pump lineage, we performed pairwise identity assessments in both primary sequence and secondary structure spaces. For a set of $N$ sequences, we computed the $N \times (N-1)/2$ pairwise Levenshtein identities, defined as $I(a, b) = 1 - d_{\text{norm}}(a, b)$, where $d_{\text{norm}}$ is the Levenshtein distance normalized by the maximum length of the two sequences ($\max(|a|, |b|)$). Secondary structure sequences were derived from secondary structure space. This comparative analysis distinguishes between sequence-level diversification and structural-level conservation, providing a physical rationale for using the proton pump architecture as a scaffold for blue-shifted designs.

$$\text{lev}_{a,b}(i, j) = \begin{cases} \max(i, j) & \text{if } \min(i, j) = 0, \\ \text{lev}_{a,b}(i-1, j-1) & \text{if } a_i = b_j, \\ 1 + \min\{\text{lev}_{a,b}(i-1, j), & \\ \quad \text{lev}_{a,b}(i, j-1), & \\ \quad \text{lev}_{a,b}(i-1, j-1)\} & \text{otherwise.} \end{cases} \tag{2}$$

Algorithmically, we implemented Wagner-Fischer (time $O(|a| \cdot |b|)$, space optimised to $O(\min(|a|, |b|))$) to fill the pairwise distance matrix.

The resulting $N \times N$ pairwise distance matrix was used to build a Minimum Spanning Tree (MST) via Kruskal's algorithm [3], selecting the edge set $E_{\text{MST}}$ that minimises the total weight $\sum_{(u,v) \in E_{\text{MST}}} d_{\text{norm}}(u, v)$ while preserving connectivity. We implemented Kruskal's algorithm with a union-find (disjoint set) data structure for efficient cycle detection [4].

Network visualisation used the Kamada-Kawai force-directed layout [5] which minimizes a global energy functional:

$$E(\mathbf{P}) = \sum_{i<j} k_{ij} (\|\mathbf{p}_i - \mathbf{p}_j\| - l_{ij})^2, \tag{3}$$

with $l_{ij} \propto d_{\text{norm}}(i, j)$ and spring constants $k_{ij} \propto d_{\text{norm}}(i, j)^{-2}$ to preserve both local community structure and long-range separations between clades. Layout parameters were chosen to balance local compactness against global separability and to visualise phylogeny-spectrum correspondence clearly.

**Spectral mapping and statistical analysis.** To visualise genotype-phenotype relationships, $\lambda_{\max}$ was discretised into $k = 8$ equidistant spectral bins $B_i$ spanning [420, 620] nm. Network communities were detected by modularity optimisation (Louvain) [6]; community medians

$$\tilde{\lambda}_j = \text{median}\big(\{\lambda_{\max}(n) : n \in C_j\}\big) \tag{4}$$

were used to colour the MST and layout. We estimated class distributions with Gaussian KDE

$$\hat{f}_h(x) = \frac{1}{nh}\sum_{i=1}^{n} K\left(\frac{x - x_i}{h}\right), \quad K(u) = \frac{1}{\sqrt{2\pi}}e^{-u^2/2}, \tag{5}$$

using Scott's rule for bandwidth selection. These density estimates quantify the "spectral censorship" region (e.g., paucity of proton pumps below 500 nm) and inform our extrapolation analyses (see Results).

**Landscape embedding and basin partitioning.** For compact visual summaries we projected high-dimensional sequence features into a 2D discriminative space using Linear Discriminant Analysis (LDA; Fisher criterion). LDA finds the projection matrix $\mathbf{W}$ that maximizes the ratio of between-class to within-class scatter [7]. We then trained a Support Vector Classifier (RBF kernel) on the LDA coordinates to annotate discrete functional basins, partitioning the landscape for downstream interpretation.

**Software and implementation.** Distance computations and MST construction used SciPy and NetworkX; force-directed layouts and community detection used python-igraph and GraphTool. KDE and plotting used Seaborn/Matplotlib. The phylogenetic analysis is integrated into a GUI dashboard that visualizes functional clades, spectral distributions, and landscape embeddings, enabling exploration of genotype-phenotype relationships and identification of spectral gaps.

| Descriptor | Ala (A) | Cys (C) | Glu (E) | Lys (K) | Trp (W) |
|---|---|---|---|---|---|
| Organic Value | 20 | 60 | 60 | 80 | 180 |
| Inorganic Value | 0 | 20 | 150 | 70 | 130 |
| Hydropathy | 1.8 | 2.5 | -3.5 | -3.9 | -0.9 |
| Polarity | 8.1 | 5.5 | 12.3 | 11.3 | 5.4 |
| logP | -2.85 | -2.49 | -3.69 | -3.05 | -1.05 |
| IEP | 6.00 | 5.07 | 3.22 | 9.74 | 5.89 |
| Mol. Weight | 89.09 | 121.16 | 146.12 | 147.20 | 204.23 |
| Volume | 60.4 | 73.4 | 85.9 | 108.5 | 143.9 |
| *Binary Indicators:* | | | | | |
| Acid-Base (Neu/Bas/Aci) | 1/0/0 | 1/0/0 | 0/0/1 | 0/1/0 | 1/0/0 |
| Polarity (Neu/Hydro) | 0/1 | 1/0 | 0/0 | 0/0 | 0/0 |
| Cleft (Ali/Aro/Cys/Neu/PG) | 1/0/0/0/0 | 0/0/1/0/0 | 0/0/0/0/0 | 0/0/0/0/0 | 0/1/0/0/0 |

Table CS1: The 18-dimensional physicochemical feature vector captures biochemical properties of each amino acid. This encoding outperforms categorical schemes for predicting spectral properties.

#### 1.2 Predicting Rhodopsin Absorption Wavelength

**Overview.** We implemented a two-stage LASSO-XGBoost stack that balances sparsity with non-linear correction to predict $\lambda_{\max}$. The pipeline, feature engineering choices, hyperparameter tuning and validation procedures are described in this section.

**Model fitting and selection.** The stacked predictor fits a LASSO baseline and trains XGBoost on residuals. LASSO regularization [8] was optimized via cyclic coordinate descent. XGBoost [9] was trained on residuals using a regularized objective with second-order Taylor approximation. The final hybrid predictor combines the sparse linear baseline with the residual ensemble through additive boosting. Final metrics are reported from shuffled 10-fold cross-validation using the implementation in `Bin.Train.regressor_test`; these folds were random sequence-level folds, not homology-cluster or leave-family-out splits.

**Regression model and encoding ablation grid.** To ensure that the selected regressor was not an artefact of a single encoding or architecture, we evaluated a crossed model-selection grid over six sequence encodings and six estimator families. The encoding grid comprised: (i) physicochemical property descriptors only; (ii) rank-ordinal amino-acid encoding; (iii) the 18-channel Feature Map; (iv) one-hot amino-acid encoding; (v) Feature Map concatenated with one-hot encoding; and (vi) physicochemical properties concatenated with one-hot encoding. The estimator grid comprised LASSO, XGBoost, Random Forest, Extra Trees, radial-basis SVR, and the two-stage LASSO→XGBoost residual stack. For each encoding–estimator pair, the same outer train/test, shuffled 10-fold cross-validation, and Monte Carlo split protocol was applied; models were compared by $R^2$, RMSE, and MAE. The complete grid was written to the per-encoding `models_summary.csv` files, while Supplementary Table CS10 reports the top-performing and most informative baselines and Supplementary Fig. CS6 visualizes architecture and encoding ablations. Family-stratified reliability was then computed for the selected Feature Map LASSO→XGBoost model and is summarized in Supplementary Table CS11.

The LASSO-only models serve as sparse linear baselines, XGBoost and tree ensembles test non-linear sequence-property interactions without residual stacking, SVR tests a kernel baseline, and the stacked model tests whether a sparse linear component plus a non-linear residual learner improves calibration. Encoding ablations test whether performance depends on the Feature Map itself: one-hot models retain residue identity without explicit physicochemical descriptors, ordinal models provide a compact rank-based representation, and concatenated encodings test whether residue identity adds signal beyond biochemical descriptors. Hyperparameter tuning was performed within each outer split when supported by the estimator; all preprocessing steps, including feature standardisation, were fit on training folds only and then applied to held-out folds.

**Validation and extrapolation experiments.** We used shuffled K-fold and Monte Carlo sequence-level splits for model comparison. These validation modes estimate performance on held-out sequences from the same curated dataset and may overestimate generalization to remote homologs. We therefore ran stricter leave-one-family-out and Wildtype-grouped homology-cluster holdout diagnostics using `Working/validation/group_holdout_validation.py`; outputs were written to the grouped-holdout results directory and are interpreted separately from the shuffled-fold headline metrics. We probed out-of-distribution generalization with an extremophile experiment: ordering sequences by $\lambda_{\max}$, training on the central 90%, and evaluating on the held-out top and bottom 5% (spectral tails). We inspected residuals for bias (correlation with observed values), assessed normality (Q-Q plots, Shapiro-Wilk) and examined heteroscedasticity (Breusch-Pagan test). We also tested for residual autocorrelation using the Durbin-Watson statistic

$$d = \frac{\sum_{t=2}^{T}(\epsilon_t - \epsilon_{t-1})^2}{\sum_{t=1}^{T}\epsilon_t^2}, \tag{6}$$

with $d \approx 2$ indicating no first-order autocorrelation. We additionally checked whether extrapolation-set predictions were directionally biased in the blue-shifted region, so candidate triage could account for systematic overestimation of absolute $\lambda_{\max}$.

**Software and implementation.** We used scikit-learn for LASSO, ridge, SVR, and SVC; XGBoost for gradient-boosted trees; numpy and pandas for data processing; scipy for statistics; and matplotlib and seaborn for visualization. SHAP (SHapley Additive exPlanations) values from TreeExplainer identified feature importance for the XGBoost component. The complete pipeline is integrated into a GUI that handles feature engineering, model selection, hyperparameter tuning, and prediction visualization.

##### 1.3 Sequence-Plausibility Filtering

**Objective.** We implemented ensemble anomaly-detection classifiers to score candidate sequences for proton-pump-like sequence plausibility and to act as a gatekeeper during selection.

**Modalities and classifiers.** We used four complementary modalities: (i) *sequence* (primary residues), (ii) *structure* (predicted secondary structure via S4Pred [10]), (iii) *hybrid* (interleaved sequence+structure tokens), and (iv) *stacked* (independent sequence and structure models whose calibrated probabilities are multiplied). Individual classifiers included Chaos Game Representation (CGR) with k-NN scoring, normalized Levenshtein distance baselines, n-gram Markov models with entropy weighting, and alignment-based Needleman-Wunsch scoring (affine gaps) [11].

**Chaos Game Representation (CGR) and distance scoring.** CGR converts a sequence $S = (s_1, \ldots, s_L)$ to a trajectory by assigning each amino acid $a \in \mathcal{A}$ to a vertex $c_a$ and iterating

$$\mathbf{p}_0 = 0, \quad \mathbf{p}_t = \tfrac{1}{2}(\mathbf{p}_{t-1} + \mathbf{c}_{s_t}) \quad (t = 1, \ldots, L), \tag{7}$$

so that $\mathbf{p}_L$ is a exponentially weighted superposition of residue vertices. Final positions were embedded in $d \in \{2, 3\}$ and indexed in a KD-tree for $k$-NN scoring. Distances to nearest neighbors were aggregated with a Minkowski $p$-norm over sorted neighbor distances $D = [d_{(1)}, \ldots, d_{(k)}]$:

$$\text{Score}_{CGR}(S) = \left( \frac{1}{k} \sum_{i=1}^{k} (d_{(i)})^p \right)^{1/p}. \tag{8}$$

We tuned hyperparameters $(k, p, d)$ on validation splits to balance recall and specificity.

**CGR properties.** The iterative map has the property that residue contributions decay exponentially with distance from the sequence end, ensuring local context dominates while preserving global composition. We aggregate distances using the $p$-norm (with $p = 1.5$ by default), where $p \to 1$ gives mean distance and $p \to \infty$ gives maximum distance.

**Geometric placements.** In practice we used both 2D and 3D class point placements. For 2D we placed points on the unit circle at angles $\theta_i = 2\pi i/20$ with coordinates $\mathbf{c}_i = (\cos\theta_i, \sin\theta_i)$. For 3D we used a Fibonacci sphere parametrisation to distribute points approximately uniformly:

$$\begin{gathered} z_i = 1 - \frac{2i+1}{20}, \quad r_i = \sqrt{1 - z_i^2}, \quad \phi_i = i\Phi, \\ \mathbf{c}_i = (r_i \cos\phi_i, r_i \sin\phi_i, z_i), \end{gathered} \tag{9}$$

with golden angle $\Phi = \pi(3 - \sqrt{5})$.

**Levenshtein distance.** We implemented the Wagner-Fischer dynamic programming algorithm to compute edit distances efficiently and produced a normalized $d_{norm}$ for downstream scoring [12]. **Markov models and alignment-based scoring.** For Markov models, we modeled sequence $S = (x_1, \ldots, x_L)$ as an $m$-th order Markov chain:

$$P(S) = P(x_1, \ldots, x_m) \prod_{i=m+1}^{L} P(x_i | x_{i-m}, \ldots, x_{i-1}). \tag{10}$$

Bigram ($m = 1$) and trigram ($m = 2$) conditional probabilities were estimated with smoothing. Laplace (add-$\alpha$) smoothing yields

$$\hat{P}(x_i | h) = \frac{C(h, x_i) + \alpha}{\sum_y (C(h, y) + \alpha)}. \tag{11}$$

For higher order contexts we used absolute-discount Kneser-Ney smoothing. For a bigram this can be written as:

$$P_{KN}(w_i|w_{i-1}) = \frac{\max(C(w_{i-1}w_i) - D, 0)}{\sum_{w'} C(w_{i-1}w')} + \lambda(w_{i-1})P_{KN}(w_i), \quad (12)$$

where $D$ is the discount (chosen by validation) and the backoff weight $\lambda(w_{i-1})$ normalises remaining probability mass and routes it to the lower-order unigram distribution. This formulation preserves mass for rare continuations while using robust lower-order estimates.

Per-sequence log-likelihoods were computed and scored using entropy weighting per history, where the entropy of a context $h$ is $H(h) = -\sum_{y\in\mathcal{A}} P(y|h)\log P(y|h)$. The weighted log-likelihood used in scoring is the length-normalised sum $\sum_i H(h_i)\log P(x_i|h_i)$. Alignment scores were computed with Needleman-Wunsch (global alignment) using affine gap penalties as described in [11, 13]. Per-sequence alignment scores were normalized and converted to calibrated probabilities prior to stacking.

**Calibration and stacking.** We calibrated raw modality scores $s_m(S)$ to probabilities via Platt scaling (logistic fit)

$$P_m(S) = \frac{1}{1 + \exp(A_m s_m(S) + B_m)}, \quad (13)$$

or via temperature scaling for logit outputs $z$: $P = \text{softmax}(z/T)$ with temperature $T$ chosen by validation. Calibrated probabilities were combined multiplicatively for the stacked model:

$$P_{stack}(S) = \prod_m P_m(S)^{w_m}, \quad (14)$$

with weights $w_m$ optimised on a validation split (or set to uniform when no modality dominated). Thresholds (e.g., $\tau_{90}$) were chosen to achieve 90% recall on the proton-pump-labeled positive training set and to quantify acceptance rates and selectivity decay on mutant bins.

Primary classification metrics were computed using random sequence-level train/test and cross-validation splits in the filtering code. These splits benchmark the implemented discriminators on held-out sequences from the curated dataset, but they can overestimate generalization across homologous groups. We therefore ran an additional grouped-holdout diagnostic using `Working/validation/group_holdout_validation.py`. This diagnostic trained a simple positive-class amino-acid bigram Markov model on proton-pump sequences and evaluated family-membership filtering under leave-one-family-out and Wildtype-grouped splits; outputs were written to the grouped-holdout results directory. Because this grouped check is intentionally simpler than the full stacked filter benchmark, it is used to assess homology sensitivity rather than to replace the random-split headline metrics.

**Mutant Library Generation.** We generated a synthetic validation library from the curated Proton Pump parent set $\mathcal{P}$. This library is a chemical proxy benchmark for mutation plausibility, not an experimentally assayed functional dataset. For each parent $P \in \mathcal{P}$ we applied a stochastic mutation operator $\mathcal{M}(P, d, \phi)$ producing a mutant sequence $S$ at mutation distance $d$ (number of substituted residues) and substitution type $\phi \in \{\phi_{cons}, \phi_{rad}\}$. Biophysical amino-acid groups partition the alphabet into disjoint sets $\mathbb{G} = \{\mathcal{G}_{hyd}, \mathcal{G}_{pol}, \mathcal{G}_{pos}, \mathcal{G}_{neg}, \mathcal{G}_{spc}\}$. For clarity we used the following grouping (single-letter codes): $\mathcal{G}_{hyd} = \{A, V, I, L, M, F, W, Y\}$, $\mathcal{G}_{pol} = \{S, T, N, Q, C\}$, $\mathcal{G}_{pos} = \{K, R, H\}$, $\mathcal{G}_{neg} = \{D, E\}$, $\mathcal{G}_{spc} = \{G, P\}$.

Conservative substitutions ($\phi_{cons}$) replace residues only within the same property group: for a site with residue $a$, the replacement $b$ is sampled such that $a, b \in \mathcal{G}_k$. Radical substitutions

($\phi_{rad}$) are forced across group boundaries: if $a \in \mathcal{G}_i$ then $b$ is drawn from $\bigcup_{j \neq i} \mathcal{G}_j$. We refer to these as conservative and radical proxy classes rather than experimentally measured functional classes. To model realistic mutation loads we sampled the mutation count $d$ from a discrete distribution (Poisson or truncated normal) chosen to produce the desired bin means (see Experimental Stratification below). Sites were sampled uniformly without replacement; substitutions were drawn uniformly from the allowed amino acids within the target group.

This process is summarised below (informal pseudocode).

**Algorithm 1** Mutant Library Generation ($\mathcal{M}$)

**Require:** Parent set $\mathcal{P}$, total mutants $N$, mutation type $\phi \in \{\phi_{cons}, \phi_{rad}\}$, bin specification $\mathcal{B}$
**Ensure:** Mutant library $\mathcal{S}$ partitioned by bin

**for** $i = 1$ to $N$ **do**
  sample parent $P \sim \mathcal{P}$
  draw mutation count $d \sim$ sample_distance($\mathcal{B}$) (Poisson or truncated normal)
  sample $d$ sites without replacement from $[1..|P|]$
  $S \leftarrow P$
  **for** each selected site $s$ **do**
    $a \leftarrow P[s]$
    **if** $\phi = \phi_{cons}$ **then**
      sample replacement $b \sim$ Uniform($\mathcal{G}(a)$) (same group)
    **else**
      sample replacement $b \sim$ Uniform($\bigcup_{j \neq k} \mathcal{G}_j$) (different group)
    **end if**
    set $S[s] \leftarrow b$
  **end for**
  assign $S$ to bin $B(d)$ (by Levenshtein distance)
  add $S$ to $\mathcal{S}$
**end for**
**return** $\mathcal{S}$

**Evaluation metrics.** Model scores $f(S)$ were standardised against the proton-pump-labeled positive training distribution, and a decision threshold $\tau_{90}$ was chosen to yield 90% recall on the training positive set. The Acceptance Rate (AR) for mutation bin $B$ and type $T$ is the empirical expectation

$$AR(B, T) = \mathbb{E}_{S \in B,\, \text{type}(S)=T} \left[ \mathbb{I}\big(f(S) > \tau_{90}\big) \right], \tag{15}$$

estimated as the sample mean of the indicator across mutants in the bin. The Selectivity Decay function measures discrimination as a function of mutation distance $d$:

$$\Delta_{sel}(d) = AR(d, \phi_{cons}) - AR(d, \phi_{rad}). \tag{16}$$

Confidence intervals for AR and $\Delta_{sel}$ were computed by nonparametric bootstrap (1000 resamples) over mutants in each bin.

**Threshold selection.** The threshold $\tau_{90}$ was obtained by solving

$$\tau_{90} = \inf\{\tau : \mathbb{P}_{S \sim \text{train}}(f(S) > \tau) \geq 0.90\}. \tag{17}$$

This fixed operating point allows direct comparison of Acceptance Rates across models and bins.

**Model and bin evaluation algorithm.** The evaluation loop used to compute $AR$ and $\Delta_{sel}$ per model and bin is given below (informal pseudocode).

**Experimental categorization.** Mutants were stratified into five bins by Levenshtein distance $d(S, P)$; here we report three representative bins used in Figures:

**Algorithm 2** Evaluation of Acceptance Rates and Selectivity Decay

**Require:** Mutant library $\mathcal{S}$ partitioned by bins $\{B_k\}$, models $\{M_j\}$, threshold $\tau_{90}$
**Ensure:** Matrices $AR$, $CI$ and $\Delta_{sel}$

**for** each model $M_j$ **do**
  **for** each bin $B_k$ **do**
    **for** each type $T \in \{\phi_{cons}, \phi_{rad}\}$ **do**
      $vals \leftarrow [\mathbb{I}(M_j.score(S) > \tau_{90}) : S \in B_k, \text{type}(S) = T]$
      $AR[j,k,T] \leftarrow \text{mean}(vals)$
      $CI[j,k,T] \leftarrow \text{bootstrap\_CI}(vals)$
    **end for**
    $\Delta_{sel}[j,k] \leftarrow AR[j,k,\phi_{cons}] - AR[j,k,\phi_{rad}]$
  **end for**
**end for**
**return** $AR, \Delta_{sel}, CI$

- Bin 1 (Low Distance): $\bar{d} = 5.27 \pm 1.2$ (local neighbourhood).
- Bin 3 (Medium Distance): $\bar{d} = 24.15 \pm 1.8$ (middle exploration zone).
- Bin 5 (High Distance): $\bar{d} = 40.92 \pm 2.4$ (search horizon limit).

The evaluation procedure ensures mutation type, distance and parent distribution are balanced between conservative and radical subsets to avoid sampling bias.

**Mutation-category library.** The synthetic validation library comprised named categories defined by parent functional class and substitution strategy; these map to the acceptance-rate groups shown in Supplementary Fig. CS25 as follows:

- *positive_functional* (**+ve parent, +ve child**): proton-pump parents subjected to within-group (conservative) substitutions; expected to retain proton-pump-like sequence plausibility.
- *positive_disruptive* (**+ve parent, −ve child**): proton-pump parents subjected to radical cross-group substitutions across the full mutation-distance range ($\bar{d} \approx 5$–$44$); shown in Supplementary Fig. CS25.
- *positive_lof_** (targeted loss-of-function; **supplementary only**): proton-pump parents with targeted loss-of-function mutations at short distances ($\bar{d} \approx 2$–$10$). Sub-classes are: *charge_reversal* (Asp/Glu ↔ Lys/Arg); *critical_residue* (substitutions at conserved retinal-binding and proton-gating positions); *proline_helix* (proline insertions into transmembrane helices). These are excluded from the main acceptance-rate panel because the sequence-plausibility filter cannot distinguish targeted single-residue loss-of-function changes from conservative proton-pump mutations, and are shown in Supplementary Fig. CS25.
- *negative_** (**Negative variants**): the same conservative and radical substitution strategies applied to non-proton-pump parents; these are expected to score low regardless of mutation type.
- *non_rhodopsins_baseline* (**Non-rhodopsin baseline**): curated non-rhodopsin ion transporters at zero mutation distance from themselves (see Negative set construction above).

| Classifier | Parameter | Default | Description |
|---|---|---|---|
| CGR | `cgr_steps` | 500 | Iterations of chaos game |
| | `dimension` | 3 | Spatial dimensionality |
| | `metric` | Cosine | Distance metric |
| | $k$ | 3 | Nearest neighbors |
| | $p$ | 1.5 | Minkowski norm |
| Levenshtein | `costs` | 1,1,1 | subst, ins, del costs |
| | `normalize` | True | Length normalization |
| Markov | `order` | 2 | Ngram size |
| | `smoothing` | Laplace | Addone smoothing ($\alpha = 1.0$) |
| | `entropy` | True | Weight by information content |
| Alignment | `algo` | NW | NeedlemanWunsch |
| | `match/miss` | +2 / -1 | Substitution matrix scaling |
| | `gaps` | -10, -1 | Affine (open, extend) |

Table CS2: Selected default parameters for anomaly detection classifiers.

Truncation variants (**_lof_truncation*) were excluded from acceptance-rate reporting as premature stop codons confound Levenshtein distance binning. Scrambled variants (**_lof_scrambled*, mean $d \approx 90$) fall in separate high-distance bins and are included in the Negative-variants group but lie beyond the acceptance-rate plot axis ($x \leq 50$). **Negative set construction (non-rhodopsins).** To create a stringent decoy set we queried bacterial proteomes and selected sequences that satisfied the following filters: length between 250 and 350 amino acids, at least one predicted transmembrane helix (TMH), annotated or putatively involved in ion transport (e.g., sodium, chloride, proton transporters) and without any rhodopsin annotation or conserved retinal-binding motif (e.g., no Lys at the Schiff-base position). Candidate decoys were further curated by manual inspection and by cross-referencing UniProt and specialized transporter databases to minimise label noise. The resulting decoy pool matched the rhodopsin set in length and general transmembrane topology while excluding true rhodopsins.

**Default parameter configuration and software.** Table CS2 summarizes the default parameters (full tables appear in Supplementary Methods):

We implemented classifiers using NumPy, scikit-learn, NetworkX, python-igraph and custom modules for ensemble classification (Markov models, Chaos Game Representation, alignment-based scoring); plotting used Matplotlib/Seaborn. All classifiers integrate into a GUI interface that allows users to configure ensemble parameters, visualize classifier performance, and interactively inspect sequence-plausibility scores.

##### 1.4 Genetic Algorithm for Mutant Generation

We model the directed evolution pipeline as a stochastic optimization process acting over $G$ discrete generations. At generation $t$, a sequence $S^{(t)}$ is drawn from the discrete sequence space $\mathcal{S} = \mathcal{A}^{L_{\text{seq}}}$, where $\mathcal{A}$ denotes the alphabet and $L_{\text{seq}}$ the sequence length. The transition $S^{(t)} \to S^{(t+1)}$ is governed by probabilistic crossover and mutation operators. In the wrapper implementation used for the reported runs, the positional prior distribution $\boldsymbol{\pi} : [1, L_{\text{seq}}] \to \mathbb{R}_{\geq 0}$ modulates mutation-site selection, whereas crossover uses the stochastic segment sampler without positional-prior weighting.

##### Prior distribution

To incorporate prior knowledge about evolutionary hotspots, we define a positional prior $\boldsymbol{\pi} = (\pi_1, \ldots, \pi_{L_{\text{seq}}})$ that weights the likelihood of mutation at each sequence position, with $\sum_{j=1}^{L_{\text{seq}}} \pi_j = 1$. In this study, $\boldsymbol{\pi}$ is obtained via Decoupled Information Theoretic Selection of Key Tuning Residues as described in [14]; we treat $\boldsymbol{\pi}$ as given and use it to modulate mutation-site selection without further modification.

##### Stochastic crossover (recombination)

Recombination between parental sequences is modeled as a stochastic crossover process with a variable number of swap events. For a given offspring, the number of crossover events $K$ is sampled from a Poisson distribution with rate $\lambda_{\text{cross}}$, so that $\mathbb{E}[K] = \lambda_{\text{cross}}$ and $\text{Var}[K] = \lambda_{\text{cross}}$ [15]. For each realized crossover event $k$, we sample a swap segment length $L$ from a truncated normal distribution restricted to $[1, L_{\text{seq}}]$ [16].

**Load - expectations.** Conditional on mutation being triggered, the number of mutation events $M \sim \text{Pois}(\lambda_{\text{mut}})$ has expectation and variance

$$\mathbb{E}[M] = \lambda_{\text{mut}} = \eta_{\text{mut}} N_{\text{valid}}, \qquad \text{Var}[M] = \lambda_{\text{mut}}.$$

This makes the mean mutation load directly proportional to the number of eligible positions and the tunable intensity $\eta_{\text{mut}}$.

Given a segment length $L$, we then choose a valid starting index from the set

$$\mathcal{I}_L = \{i \in [1, L_{\text{seq}}] : i + L - 1 \leq L_{\text{seq}}\},$$

using a uniform distribution over valid segment starts. The probability of choosing position $j$ is

$$P(i = j) = \frac{\mathbb{I}(j \in \mathcal{I}_L)}{|\mathcal{I}_L|},$$

where $\mathbb{I}(\cdot)$ is the indicator function.

Given a primary parent $P_1$ and a secondary parent $P_2 \sim \text{Uniform}(\mathcal{P}^{(t)}_{\text{parents}})$, the recombination operation constructs a new offspring sequence by splicing a segment from $P_2$ into $P_1$:

$$S_{\text{new}} = P_1[1{:}i] \oplus P_2[i{:}i + L] \oplus P_1[i + L{:}\text{end}],$$

where $\oplus$ denotes concatenation. To preserve alignment validity in gapped sequences, we enforce a gap validation constraint

$$\mathcal{G}(P_1, P_2, i, L) = \mathbb{I}(\texttt{-} \notin P_1[i{:}i + L] \cup P_2[i{:}i + L]),$$

such that crossover events are only accepted if both parental segments are free of gaps over the recombined region.

##### Hotspot-guided mutagenesis

Mutation is implemented as a compound stochastic process that is also modulated by the positional prior $\boldsymbol{\pi}$. The procedure consists of three stages.

**Trigger.** For each sequence, mutation is applied with probability $\rho_{\text{mut}}$ via a Bernoulli trial

$$I_{\text{mut}} \sim \text{Bernoulli}(\rho_{\text{mut}}).$$

Sequences with $I_{\text{mut}} = 0$ are carried forward unmutated.

**Load.** Conditional on $I_{\text{mut}} = 1$, the number of mutation events $M$ is drawn from a Poisson distribution whose rate scales with the number of valid (mutable) positions:

$$M \sim \text{Pois}(\lambda_{\text{mut}}), \qquad \lambda_{\text{mut}} = \eta_{\text{mut}} \cdot N_{\text{valid}},$$

where $\eta_{\text{mut}}$ is a tunable mutation load parameter and $N_{\text{valid}}$ denotes the number of positions eligible for mutation.

**Site selection and substitution.** Mutation sites are sampled without replacement from the set of valid indices $\mathcal{J}$, again using the prior to bias toward evolutionarily permissive positions:

$$P(\text{site} = j \mid \boldsymbol{\pi}) = \frac{\pi_j \cdot \mathbb{I}(j \in \mathcal{J})}{\sum_{k \in \mathcal{J}} \pi_k}.$$

This scheme concentrates mutational exploration on hotspots inferred by the prior [14]. For each selected site, the replacement amino acid is drawn uniformly from the canonical alphabet $\mathcal{A}$, ensuring that the mutation process remains agnostic to the downstream fitness model.

#### Parent selection and generational transition

At each generation, selection operates at the population level. Let $\mathcal{P}^{(t)} = \{S_1^{(t)}, \ldots, S_{N_{\text{pop}}}^{(t)}\}$ denote the population at generation $t$, and let

$$\mathbf{F}^{(t)} = \left[F(S_1^{(t)}), \ldots, F(S_{N_{\text{pop}}}^{(t)})\right]^\top$$

denote the corresponding vector of fitness values. We select the top $n_{\text{parents}}$ sequences by fitness ranking:

$$\mathcal{P}_{\text{parents}}^{(t)} = \left\{ S_i^{(t)} \in \mathcal{P}^{(t)} : i \in \text{argsort}_{1:n_{\text{parents}}} \left( \mathbf{F}^{(t)} \right) \right\}.$$

The next generation is then generated by iteratively applying crossover followed by mutation to sampled parents. For each child index $c \in [1, n_{\text{children}}]$, we sample a primary parent $P_1^{(c)} \in \mathcal{P}_{\text{parents}}^{(t)}$, perform recombination with a secondary parent, and subsequently apply hotspot-guided mutagenesis:

$$\mathcal{P}^{(t+1)} = \left\{ \Omega_{\text{mut}} \left( \Phi_{\text{cross}}(P_1^{(c)}, \mathcal{P}_{\text{parents}}^{(t)}) \right) : c \in [1, n_{\text{children}}] \right\},$$

where $\Phi_{\text{cross}}$ denotes the crossover operator and $\Omega_{\text{mut}}$ the mutation operator defined above.

#### Constraint-based hierarchical selection

To prevent the accumulation of sequences that are high in predicted fitness but far from the intended proton-pump-like sequence distribution, we augment fitness-based selection with an explicit constraint filter, implementing a "generate-and-filter" scheme. Offspring are evaluated both by predictive fitness $F(S)$ (here, the squared distance to the target) and by an auxiliary constraint function $\mathcal{C} : \mathcal{S} \to \mathbb{R}$ that returns a scalar proton-pump sequence-plausibility score. In the implementation used here, $\mathcal{C}$ is the Markov sequence likelihood under the proton-pump family model, so higher values indicate stronger membership in the learned proton-pump-like sequence distribution rather than direct experimental stability or folding propensity.

We first define a reference distribution for the constraint score using positive proton-pump training examples $\{S_1, \ldots, S_{N_{\text{pos}}}\}$. For a configured filter strength $q \in [0,1]$, the sequence-plausibility threshold is the empirical $q$-quantile of the positive-score distribution:

$$\tau_{\mathcal{C}} = Q_q \left( \{\mathcal{C}(S_n)\}_{n=1}^{N_{\text{pos}}} \right).$$

A candidate sequence $S$ is deemed acceptable only if its constraint score meets or exceeds this threshold:

$$\mathbb{I}_{\text{accept}}(S) = \mathbb{I}\big(\mathcal{C}(S) \geq \tau_{\mathcal{C}}\big).$$

To integrate this constraint into selection, we define an effective fitness

$$F_{\text{eff}}(S) = \begin{cases} F(S), & \text{if } \mathbb{I}_{\text{accept}}(S) = 1, \\ \infty, & \text{otherwise,} \end{cases}$$

such that low-scoring sequences are excluded from the parent pool. The main directed-evolution run used filter strength $q = 0.2$, while the schema-ablation grid varied $q$ from 0.05 to 0.4.

#### Experimental hyperparameters

All simulations were initialized from wild-type proton-pump sequences after applying the wrapper's proton-pump filter. Unless otherwise specified, we used the following hyperparameters.

**Simulation.** We evolved populations for up to $G = 100$ generations with a fixed population size $N_{\text{pop}} = 50$, $n_{\text{parents}} = 15$ selected parents per generation, and 5 elite sequences retained between generations.

**Crossover.** The Poisson rate for the number of crossover events was set to $\lambda_{\text{cross}} = 1.0$, and the truncated normal distribution for segment lengths had mean $\mu_{\text{cross}} = 6$ and standard deviation $\sigma_{\text{cross}} = 6$.

**Mutation.** The per-sequence mutation trigger probability was $\rho_{\text{mut}} = 1.0$, and the mutation load parameter was $\eta_{\text{mut}} = 0.01$.

#### Functional landscape mapping (LDA & SVC)

To visualize the evolutionary trajectory within the context of the natural protein family, we projected the high-dimensional sequence space ($d = 9990$) onto a 2D latent manifold using Linear Discriminant Analysis (LDA) [7]. We trained a Support Vector Classifier (SVC) with a Radial Basis Function (RBF) kernel on the LDA coordinates to define discrete functional basins, partitioning the landscape into the five observed regions.

#### Evolutionary dynamics metrics

Genetic diversity during the trajectory was quantified using Shannon's Positional Entropy ($H$). For each position $i$ in the multiple sequence alignment of the population at generation $g$:

$$H_i = -\sum_{a \in \mathcal{A}} p_a \log_2 p_a,$$

where $\mathcal{A}$ is the set of 20 amino acids and $p_a$ is the frequency of residue $a$ at position $i$. To capture active search dynamics rather than background conservation, entropy statistics (min, mean, median) were computed exclusively for the top 10% most variable positions (highest $H_i$ for a given generation). Phases of "Exploration" were defined by rising mean entropy (population diversification), while "Exploitation" phases were defined by collapsing entropy (fixation of beneficial alleles).

Convergence was defined via two complementary criteria. First, a fitness-stagnation criterion monitors the improvement of the best fitness: let $F_{\min}^{(t)}$ denote the minimum fitness at generation $t$, and define the improvement

$$\Delta^{(t)} = F_{\min}^{(t-1)} - F_{\min}^{(t)}.$$

The simulation is terminated early if the improvement in best fitness remains below a small threshold $\epsilon = 0.1$ for 10 consecutive generations:

$$\sum_{\tau=T-9}^{T} \mathbb{I}\big(\Delta^{(\tau)} < \epsilon\big) \geq 10.$$

Second, we use a direct proximity-to-target convergence definition based on the predicted wavelength: a run is considered to have "converged" at generation $t^*$ if the best-predicted wavelength satisfies

$$\left|\hat{\lambda}_{\max}^{(t^*)} - \lambda_{\text{target}}\right| \leq 0.01\,\lambda_{\text{target}},$$

(i.e., within 1**Acceptance rate and selectivity decay.** To quantify filter behaviour on synthetic mutants we use the Acceptance Rate (AR) for mutation bin $B$ and type $T$ (conservative or radical) [17], defined as $AR(B,T) = \mathbb{E}_{S\in B,\text{type}(S)=T}\,[\mathbb{I}(f(S) > \tau_{90})]$, where $\tau_{90}$ is the decision threshold yielding 90% recall on the positive proton-pump training set. The Selectivity Decay $\Delta_{sel}(d)$ measures the difference in acceptance between conservative and radical proxy mutations as a function of evolutionary distance $d$: $\Delta_{sel}(d) = AR(d, \phi_{cons}) - AR(d, \phi_{rad})$. Higher $\Delta_{sel}$ indicates stronger discrimination between the two synthetic proxy classes.

#### Main evolution algorithm

The constrained directed evolution procedure used in our simulations is summarized in Algorithm 3 (pseudocode). Selection is truncation based using effective fitness; non-viable sequences (constraint score below threshold) are excluded from parent selection.

#### Derivation and calibration of the positional prior

The positional prior $\boldsymbol{\pi}$ was derived from LASSO coefficients and XGBoost feature importances (SHAP values) learned during regressor training on the full rhodopsin dataset (n=884 sequences across functional classes). For each of the $L_{\text{seq}} = 250$ positions, we computed:

$$\pi_j^{\text{LASSO}} = |\beta_j^{\text{LASSO}}|, \tag{18}$$

$$\pi_j^{\text{SHAP}} = \frac{1}{n}\sum_{i=1}^{n} |\text{SHAP}_j(x_i)|, \tag{19}$$

where $\beta_j^{\text{LASSO}}$ are the fitted LASSO coefficients and $\text{SHAP}_j(x_i)$ denotes the SHAP value for position $j$ in sequence $i$ as returned by the XGBoost model. The two importance vectors were normalized to probability distributions (summing to 1) and combined via a weighted average:

$$\pi_j = w_{\text{LASSO}} \cdot \frac{\pi_j^{\text{LASSO}}}{\sum_k \pi_k^{\text{LASSO}}} + (1 - w_{\text{LASSO}}) \cdot \frac{\pi_j^{\text{SHAP}}}{\sum_k \pi_k^{\text{SHAP}}}, \tag{20}$$

with weight $w_{\text{LASSO}} = 0.5$ to balance linear and non-linear feature importance. This prior concentrates evolutionary search on the $\sim 20$ most important positions (accounting for $> 80\%$ of cumulative importance), primarily located in the retinal-binding pocket and Schiff base linker region, while allowing rare mutations at lower-importance positions. The prior

**Algorithm 3** Constrained Directed Evolution

**Require:** Initial population $\mathcal{P}^{(0)}$, fitness $F$, constraint $\mathcal{C}$, target $\lambda^*$, max generations $G$
**Ensure:** Best sequence $S^*$ and trajectory history

compute threshold $\tau_{\mathcal{C}} \leftarrow Q_q(\mathcal{C}(S_{\text{pos}}))$
$S^* \leftarrow \arg\min_{S \in \mathcal{P}^{(0)}} F(S)$; $F^* \leftarrow F(S^*)$
*patience* $\leftarrow 0$
**for** $t = 1$ to $G$ **do**
    $\mathcal{P}^{(t)}_{\text{parents}} \leftarrow \text{TopK}(\mathcal{P}^{(t-1)}, n_{\text{parents}}, F)$
    $\mathcal{P}^{(t)} \leftarrow \text{TopK}(\mathcal{P}^{(t-1)}, n_{\text{elite}}, F)$
    **for** $c = 1$ to $n_{\text{children}} - n_{\text{elite}}$ **do**
        **repeat**
            $P_1 \leftarrow \text{Sample}(\mathcal{P}^{(t)}_{\text{parents}})$
            $S_c \leftarrow \Phi_{\text{cross}}(P_1, \mathcal{P}^{(t)}_{\text{parents}})$ ▷ Crossover
            $S_c \leftarrow \Omega_{\text{mut}}(S_c)$ ▷ Mutation
        **until** $S_c \notin \mathcal{P}^{(t)}_{\text{parents}}$
        **if** $\mathcal{C}(S_c) \geq \tau_{\mathcal{C}}$ **then**
            add $S_c$ to $\mathcal{P}^{(t)}$
        **end if**
    **end for**
    $S^{(t)}_{\text{best}} \leftarrow \arg\min_{S \in \mathcal{P}^{(t)}} F(S)$
    **if** $F(S^{(t)}_{\text{best}}) < F^* - 0.1$ **then**
        $S^* \leftarrow S^{(t)}_{\text{best}}$; $F^* \leftarrow F(S^*)$; *patience* $\leftarrow 0$
    **else**
        *patience* $\leftarrow$ *patience* $+ 1$
    **end if**
    **if** *patience* $\geq 10$ **then**
        **break** ▷ Early stopping
    **end if**
**end for**
**return** $S^*$, trajectory

is computed from the regressor's learned feature importances and held fixed throughout all GA runs to ensure repeatability and to isolate the effect of the evolutionary algorithm. The complete directed evolution pipeline is accessible through a GUI that manages prior computation, filter configuration, and iterative GA execution.

#### Computational complexity and runtimes

Fitness evaluation dominates runtime (feature extraction + inference). The reported main runs used up to $G = 100$ generations and $N = 50$ sequences per generation; the schema-ablation grid used $G = 40$ and $N = 50$. Per-generation complexity scales approximately as $O(N \cdot L)$ for variation plus $O(N \cdot L \cdot d)$ for feature extraction where $d$ is the feature dimensionality.

#### Software and reproducibility

All evolutionary code is implemented in Python and provided in the repository; simulation configuration files and random seeds are saved per replicate to ensure full reproducibility.

For structural visualization of the accepted APR designs, each AlphaFold 3 model sequence was globally aligned to its closest wild-type proton-pump homolog. Per-residue substitution dissimilarity was computed from BLOSUM62 as

$$d_i = \max\{0, \mathrm{BLOSUM62}(a_i, a_i) - \mathrm{BLOSUM62}(a_i, h_i)\},$$

where $a_i$ is the APR residue and $h_i$ is the aligned homolog residue; positions aligned to gaps used the BLOSUM62 gap fallback score of $-4$. The resulting per-residue distances were min–max normalized to 0–100 and written into the CIF B-factor field for the Fig. 5f colouring. This structural colouring is therefore a BLOSUM62 substitution-distance map, whereas GA candidate-distance panels report Levenshtein distance from the closest wild-type homolog.

#### 1.5 GA Ablation Study

##### Hyperparameter Sensitivity Analysis

We conducted a Monte Carlo hyperparameter sweep over six evolutionary parameters using the random-search implementation in `Working/10_evolution_hyperparam_search/`. The sweep sampled 200 configurations with one random seed per configuration. Population size was sampled uniformly from 30–80, parent count was sampled as 10–40% of the population, mutation factor was sampled log-uniformly from 0.0005–0.02, crossover rate was sampled uniformly from 0.5–1.0, crossover length from 3–15, and crossover variance from 2–12. One cached model fit was reused across runs where possible, and each run saved its sampled parameters and seed in the corresponding result directory.

This sweep was designed as a broad sensitivity screen rather than the final biological design experiment. The wavelength regressor, sequence-plausibility filter family, target objective, and sequence-processing pipeline were held fixed while the evolutionary operator settings were varied. The six ablated hyperparameters correspond to the main degrees of freedom in the wrapper GA: population breadth, truncation-selection pressure, per-child mutation load, probability of recombination, typical recombination segment length, and recombination-length variance. The resulting distributions are reported in the hyperparameter main-effect and interaction panels (Supplementary Figs. CS27 and CS28); the schema-grid ANOVA reported in Supplementary Table CS15 was analysed separately from this random hyperparameter screen.

Performance was assessed using two metrics:

- *Convergence generation*: The first generation $t^*$ where the best-predicted wavelength satisfies $|F(S_{\text{best}}^{(t^*)}) - \lambda_{\text{target}}| \leq 0.01 \cdot \lambda_{\text{target}}$ (i.e., within 1% of target).
- *Final predicted wavelength*: The wavelength predicted for the best-performing sequence at the final recorded generation (or at convergence if the run stopped early).

We grouped convergence generation values into discrete classes (bins: 1-3 gen, 4-6 gen, 7-9 gen, 10-12 gen, $\geq 13$ gen) to stabilize estimation and facilitate comparison. Empirical cumulative distribution functions (ECDFs) were computed for each parameter-class combination and visualized. Main effects were quantified using one-way ANOVA with Levene's homogeneity test; effect sizes (eta-squared $\eta^2$) were computed as

$$\eta^2 = \frac{\text{SS}_{\text{between}}}{\text{SS}_{\text{total}}}, \tag{21}$$

where SS denotes the sum of squares. Two-way interactions were studied using heatmap visualization (mean convergence time colored by parameter pair) and interpreted for synergistic patterns.

##### Ablation of Sequence-Plausibility Filter, Target, and Positional Prior

We assessed the sensitivity of GA outcomes to Markov filter strength and positional mutation prior. The schema-ablation grid used seven prior types (regressor, entropy, inverse-regressor, inverse-entropy, uniform, MI, inverse-MI), eight filter strengths (0.05–0.4), and three target wavelengths (490, 450, 410 nm). Each filter-prior-target condition was run with three independent seeds (42, 123, 456), yielding $7 \times 8 \times 3 \times 3 = 504$ individual runs. The grid varied filter stringency by changing the quantile threshold derived from positive proton-pump training scores. Uniform prior runs serve as the no-hotspot baseline for the mutation prior. A true no-filter condition was implemented separately by setting `filter_model_name=none`, which disables Markov scoring so all generated candidates pass; this is distinct from increasing the quantile threshold, because higher quantiles are more restrictive in the wrapper. The no-filter condition was run for all seven priors, three targets, and three seeds ($7 \times 3 \times 3 = 63$ runs). The final no-filter candidates were then rescored post hoc with a normally trained Markov sequence filter, and pass/fail status was computed at positive-training-score quantile thresholds of 0.05–0.50. The baseline evolution parameters were population size 50, parent count 15, mutation rate 1.0, mutation factor 0.01, crossover rate 1.0, crossover length 6, crossover variance 6, and elitism 5. Implementation used the wrapper-based GA in `Bin.wrapper`, maintaining identical sequence sampling and selection logic across grid conditions.

The seven positional priors were chosen to test whether mutation-site bias improves directed search or merely changes mutation distance. The *uniform* prior samples all eligible positions equally. The *entropy* prior upweights variable alignment positions, whereas *inverse-entropy* upweights conserved positions. The *MI* prior upweights positions with high mutual information with $\lambda_{\max}$, whereas *inverse-MI* deliberately biases away from those positions. The *regressor* prior uses the normalised combination of LASSO coefficient magnitude and XGBoost SHAP importance from the wavelength model, whereas *inverse-regressor* biases away from regressor-important positions. All non-uniform priors were normalised to sum to one and blended with a uniform floor before sampling mutation sites, so every position retained non-zero probability.

The target-wavelength axis tests increasing extrapolation difficulty. The 490 nm target is inside the blue target window used for candidate selection, 450 nm is more strongly blue-shifted but still near observed channel-like wavelengths, and 410 nm is a deliberately hard blue target used to expose failure modes under stringent filter thresholds. The filter-strength axis tests the trade-off between sequence plausibility and optimization freedom: low quantiles are permissive, while higher quantiles require generated candidates to remain closer to the positive proton-pump Markov-score distribution. The no-filter arm isolates the scalar wavelength objective from the sequence-plausibility constraint and is therefore used to test whether apparent convergence produces proton-pump-like candidates or only regressor-favourable sequences. Success-rate summaries are reported in Supplementary Table CS16, component-wise effects in Supplementary Table CS15, LDA trajectory panels in Supplementary Figs. CS29–CS31, and aggregate trajectory/success diagnostics in Supplementary Figs. CS32–CS35.

**Metrics and statistical procedures.**

- *Convergence time*: Generations to reach 1% wavelength tolerance; reported as median ± 95% bootstrap CI (n=1000 resamples).
- *Filter acceptance rate*: Fraction of proposed candidates passing the Markov threshold, where the threshold is the configured quantile of positive proton-pump training scores.
- *Low-score fraction*: Proportion of proposed candidates with Markov scores below the run-specific threshold.
- *Mutation distance*: Levenshtein distance from wild-type reference at convergence; stratified into discrete classes (12-18, 19-25, 26-35 mutations) to analyse robustness.
- *Success rate*: Fraction of runs achieving convergence by the final generation; reported per condition and per target wavelength.

Statistical summaries were computed in Python (SciPy, NumPy, Pandas) with scripts in Working/10_evolution_hyperparam_search/ and Working/11_evolution_schema_test/. Main effects were summarized using ANOVA over sampled configurations and class-binned hyperparameter groups; because the random-search sweep used one seed per sampled configuration, these statistics should be interpreted as sensitivity screens rather than replicated causal estimates for each exact hyperparameter setting.

For the schema-ablation grid, statistical tests were computed on replicate-level run summaries after grouping by target wavelength, filter strength, and prior type. One-way and multi-factor ANOVA were used for convergence generation and final predicted wavelength, and effect sizes were reported as $\eta^2$. Interaction heatmaps were generated for filter-strength by prior, filter-strength by target, and prior by target contrasts. Success was defined before aggregation as reaching the target within 1% tolerance by the final generation; per-condition success rates were then averaged over the three random seeds and, where stated, over prior types. LDA trajectory plots used the same trained LDA/SVC landscape as the main GA visualization and therefore compare accepted and rejected candidates on a common functional-class projection.

**Software and implementation.** All hyperparameter sweeps, filter-strength grids, prior grids, and summary statistics were implemented in Python using NumPy, SciPy, Pandas, and Matplotlib. The schema grid maintains identical baseline evolutionary parameters while varying filter strength, target wavelength, prior type, and seed. A GUI-based dashboard displays comparisons of convergence distributions, success rates, and interactions among these settings.

### 2 Computational Supplementary Results

#### 2.1 Phylogenetic Analysis

Secondary-structure MSTs and pairwise distance distributions complement the primary-sequence analysis in the main text. The primary-sequence identity assessment also reports BLAST-like nearest-neighbour support: after duplicate removal, 296 of 309 proton-pump sequences (95.8%) have at least one other proton-pump sequence with minimum $E < 0.01$ (Table CS9). This supports the use of sequence-neighbourhood diagnostics while showing that pairwise identity across the full proton-pump cluster remains broad.

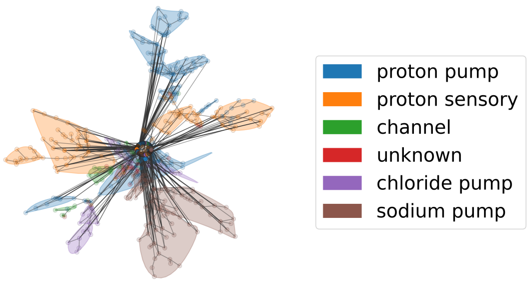

(a) Secondary-structure MST by functional class

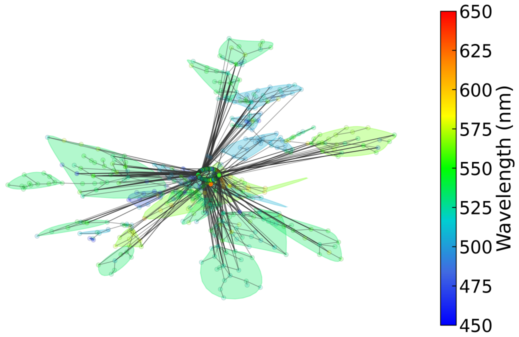

(b) Secondary-structure MST by $\lambda_{\max}$

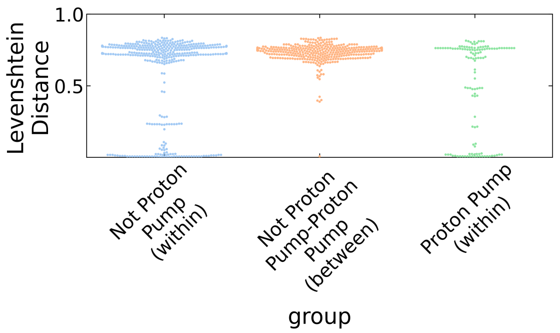

(c) Primary-sequence pairwise Levenshtein distance distributions (log scale)

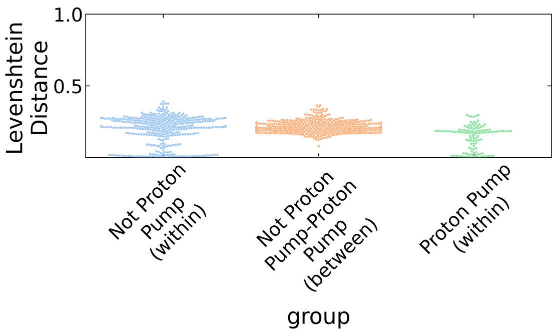

(d) Secondary-structure pairwise distance distributions (log scale)

Figure CS1: **Secondary-structure MSTs and pairwise distance distributions.** (a) Secondary-structure MST colored by functional class. (b) Same MST colored by $\lambda_{\max}$. (c) Primary-sequence Levenshtein distance distributions (log scale) for proton pumps versus other classes; proton pumps are internally conserved. (d) Secondary-structure distance distributions, showing higher structural conservation (mean pairwise identity 74.03%) than primary-sequence conservation (mean 29.43%). Close-homolog support for the proton-pump set is summarized in Supplementary Table CS9.

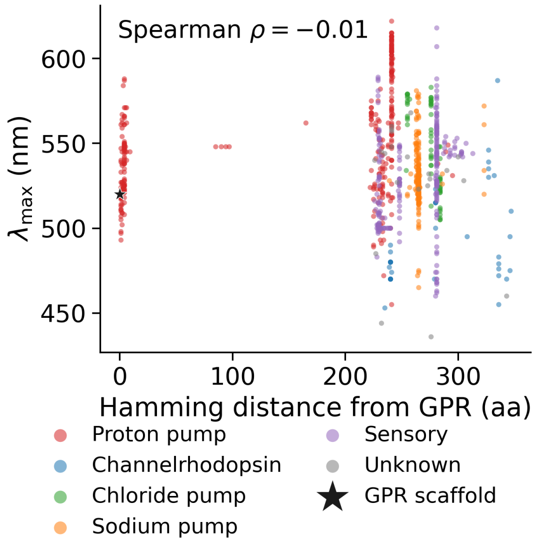

(a) $\lambda_{\max}$ vs GPR distance

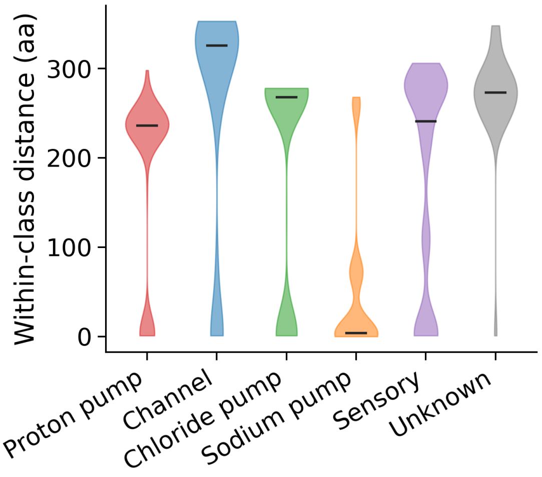

(b) Within-class Levenshtein distances

Figure CS2: **Supplementary sequence-distance panels.** (a) $\lambda_{\max}$ versus Levenshtein distance from the GPR scaffold. (b) Within-class pairwise Levenshtein distances; proton pumps are internally conserved relative to channelrhodopsins.

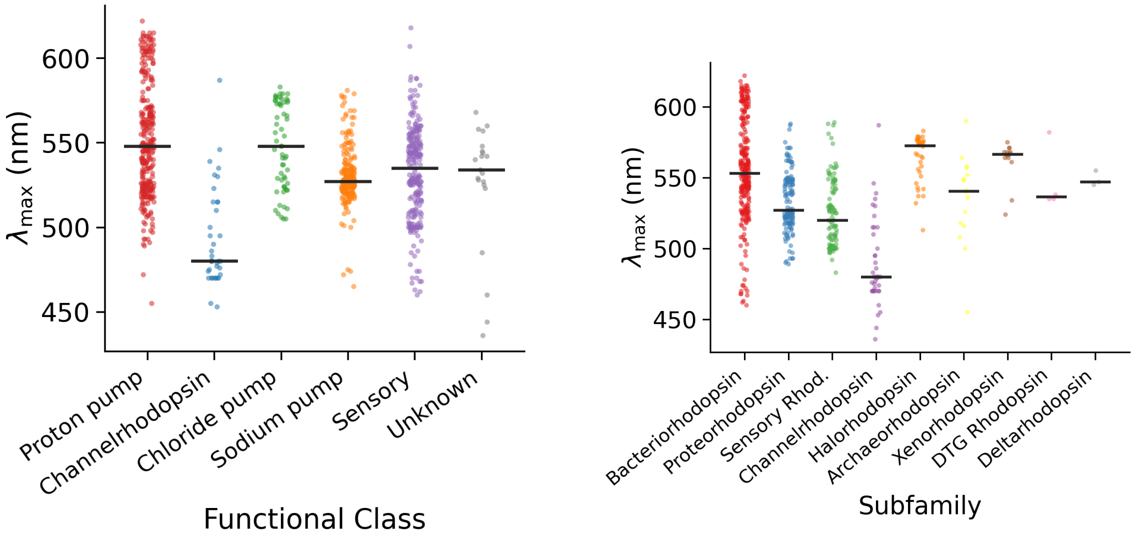

(a) $\lambda_{\max}$ by functional class

(b) $\lambda_{\max}$ by subfamily

Figure CS3: **Absorption wavelength distributions by functional class and subfamily.** (a) Swarm plot of $\lambda_{\max}$ stratified by functional superclass; the paucity of proton pumps below 500 nm motivates the blue-shifted design targets. (b) Swarm plot by phylogenetic subfamily, showing the spectral range accessible within each clade.

```
Alignment of five family representatives (43-519 of 555 columns).
Classes follow Inoue et al. (2021). lambda_max values in nm.

                      43                                                     102
BR      Proton pump    560nm  ------------------------------------------------------------
KR2     Sodium pump    525nm  ------------------------------------------------------MTQELG
NpHR    Chloride pump  577nm  -------------------------M--TETLPPVTESA---VALQAEVTQR-ELFEFVL
NpSRII  Sensory rhod.  498nm  ------------------------------------------------------------
CrChR2  Channel (cat.) 470nm  MDYGGALS-------AVGREL----L--FVTNPVVV-NG---SVL-VPED---QCY-CAG

                      103                                                    162
BR      Proton pump    560nm  -----------------Q--AQITGRPEWIWLALGTALMGLGTLYFLVKGMGVSDPDAKK
KR2     Sodium pump    525nm  NANF---ENFIGATEGFS--EIAYQFTSHILT-LGYAVMLAGLLYFILTIKNVDKKFQMS
NpHR    Chloride pump  577nm  N---------------------DPLLASSLYI--NIALAGLSILLFVFMTRGLDDPRAKL
NpSRII  Sensory rhod.  498nm  ----------------------MVGLTTLFWL--GAIGMLVGTLAFAWAGRDAGSGE-RR
CrChR2  Channel (cat.) 470nm  ------------WIESRG--TNGAQTASNVLQ-W-LAAGFSILLLMFYAYQTWKSTCGWE

                      163                                                    222
BR      Proton pump    560nm  FYAITT-LVPAIAFTMYLSMLLGYGLTMVPF-----G-------------GEQNP---IY
KR2     Sodium pump    525nm  NILSAV--VMVSAFLLLYAQAQNWTSSFTFN-EEV-GRYF------LD--PSGDL--FNN
NpHR    Chloride pump  577nm  IAVSTI-LVPVVSIASYTGLASGLTISVLEM-PA--GHFAEGSSVMLGG--EEVDGVVTM
NpSRII  Sensory rhod.  498nm  YYVTLV-GISGIAAVAYAVMALG--VG--WV-P-------------VA---ERT----VF
CrChR2  Channel (cat.) 470nm  EIYVCA--IEMVKVILEFFFEFKNPSM-LYL-AT--G----------------HR---VQ

                      223                                                    282
BR      Proton pump    560nm  WARYADWLFTTPLLLLDLALL------V------------D-----ADQGTILA-LVGA-
KR2     Sodium pump    525nm  GYRYLNWLIDVPMLLFQILFV------VSLTT--------S-----KFSSVRNQ-FWFS-
NpHR    Chloride pump  577nm  WGRYLTWALSTPMILLALGLL------A------------G-----SNATKLFT-AITF-
NpSRII  Sensory rhod.  498nm  VPRYIDWILTTPLIVYFLGLL------A------------G-----LDSREFGI-VITL-
CrChR2  Channel (cat.) 470nm  WLRYAEWLLTCPVILIHLSNL------TGL-S--------N-----DYSRRTMG-LLVS-

                      283                                                    342
BR      Proton pump    560nm  ---DGIMIGTGLVGAL--TKV---------YSYRFVWWAIST-AAMLY-------ILY-V
KR2     Sodium pump    525nm  ---GAMMIITGYIGQF--YEV-------SNLTAFLVWGAISS-AFFFH-------ILW-V
NpHR    Chloride pump  577nm  ---DIAMCVTGLAAAL--TTS-------SHLM-RWFWYAISC-ACFLV-------VLY-I
NpSRII  Sensory rhod.  498nm  ---NTVVMLAGFAGAM--VPG---------IE-RYALFGMGA-VAFIG-------LVY-Y
CrChR2  Channel (cat.) 470nm  ---DIGTIVWGATSAM--A-T-------G--YVKVIFFCLGL-CYGAN-------TFF-H

                      343                                                    402
BR      Proton pump    560nm  LFFGF-T------------------S---------KA--ESMRPE---------------
KR2     Sodium pump    525nm  MKKVI-N------------------E---------GKE-GI-SPAG--------------
NpHR    Chloride pump  577nm  LLVEW-A------------------Q---------DA--KAAGTA---------------
NpSRII  Sensory rhod.  498nm  LVGPM-T------------------E---------SA--SQRSSG---------------
CrChR2  Channel (cat.) 470nm  AAKAY-I------------------E---------GY--HTVPKGR--------------

                      403                                                    462
BR      Proton pump    560nm  -----VASTFKVLRNVTVVLWSAYPVVWLIGSE-G-A-GI---------V--PLNIETLL
KR2     Sodium pump    525nm  ------QKILSNIWILFLISWTLYPGAYLMPYLTGVD-GFLYS------ED-GVMARQLV
NpHR    Chloride pump  577nm  -------DMFNTLKLLTVVMWLGYPIVWALGVE-G-I-AV---------L--PVGVTSWG
NpSRII  Sensory rhod.  498nm  -----IKSLYVRLRNLTVVLWAIYPFIWLLGPP-G-V-AL---------L--TPTVDVAL
CrChR2  Channel (cat.) 470nm  -----CRQVVTGMAWLFFVSWGMFPILFILGPE-G-F-GV---------L--SVYGSTVG

                      463                                                 519
BR      Proton pump    560nm  FMVLDVSAKVGFGLI-LLR---SRAI-FGEAEA-----PEPSAG----DGAAATSD-
KR2     Sodium pump    525nm  YTIADVSSKVIYGVL-LGN-LAITLS--KNKEL-----VEANS--------------
NpHR    Chloride pump  577nm  YSFLDIVAKYIFAFL-LLN---YLTS-N-ESVV-----SGSILDV--PSASGTPADD
NpSRII  Sensory rhod.  498nm  IVYLDLVTKVGFGFI-ALD---AAAT-L-RAEH-----GESLAGV--DTDTPAVAD-
CrChR2  Channel (cat.) 470nm  HTIIDLMSK------------------------------------------------
```

Figure CS4: **Multiple sequence alignment of five family representatives.** One wildtype sequence per functional class, shown over alignment columns 43–519 (gap-flanked columns omitted; dashes represent insertions relative to other sequences). Functional-class annotations follow [1]. **BR**: *Halobacterium salinarum* bacteriorhodopsin (proton pump, $\lambda_{\max} = 560\,\text{nm}$); **KR2**: *Krokinobacter eikastus* rhodopsin 2 (sodium pump, 525 nm); **NpHR**: *Natronomonas pharaonis* halorhodopsin (chloride pump, 577 nm); **NpSRII**: *N. pharaonis* sensory rhodopsin II (498 nm); **CrChR2**: *Chlamydomonas reinhardtii* channelrhodopsin 2 (cation channel, 470 nm). Column numbers refer to the MAFFT multiple sequence alignment used throughout this study (555 total columns).

#### 2.2 Spectral Predictor Diagnostics

The main text reports core wavelength-prediction performance. Residual-distribution and normality diagnostics are shown below, together with architecture-selection, encoding-ablation, cross-validation residual, spectral-tail, and model-interpretability panels (Figs. CS5–CS9; Table CS11).

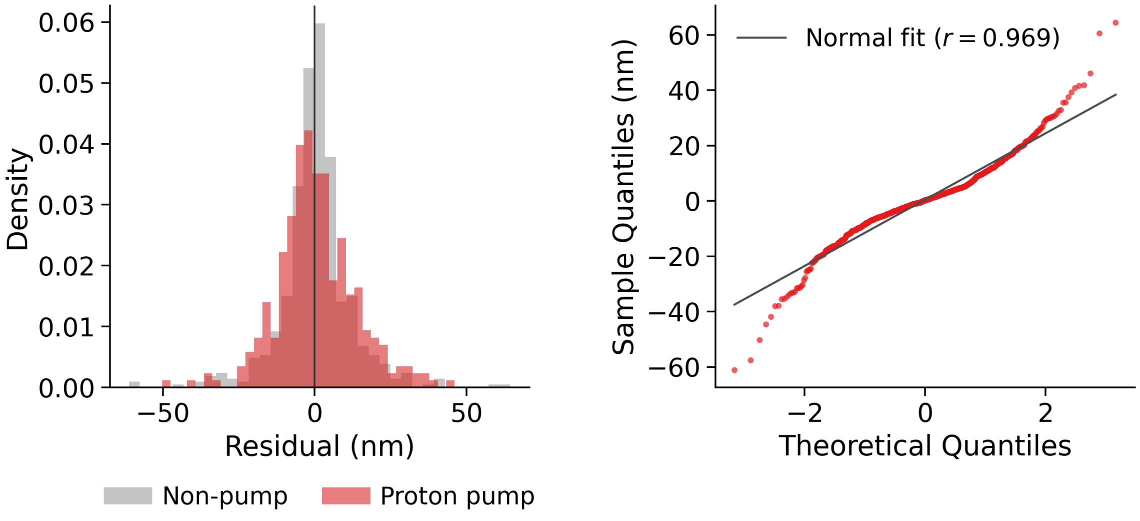

(a) Residual histogram (b) Q-Q plot

Figure CS5: **Residual diagnostics for the stacked $\lambda_{\max}$ predictor.** (a) Residual density; distributions are approximately zero-centred across functional classes. (b) Q-Q plot; slight heavy tails indicate mild non-normality at the extremes ($r = 0.969$).

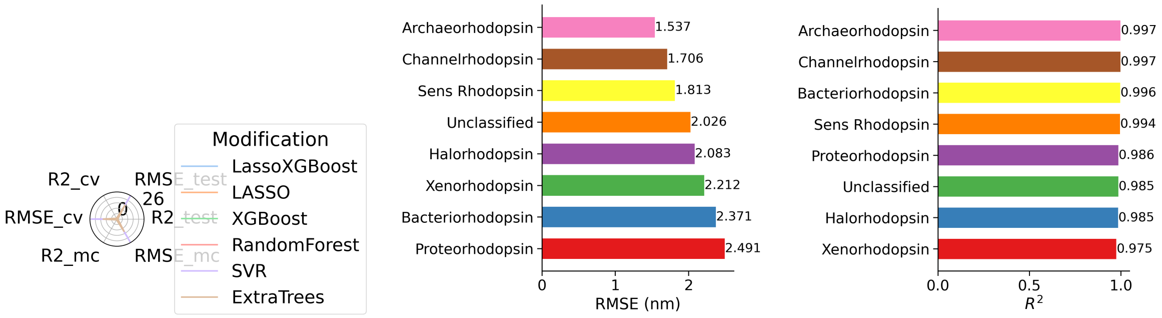

(a) Architecture comparison (b) Stacked model RMSE (c) Stacked model $R^2$

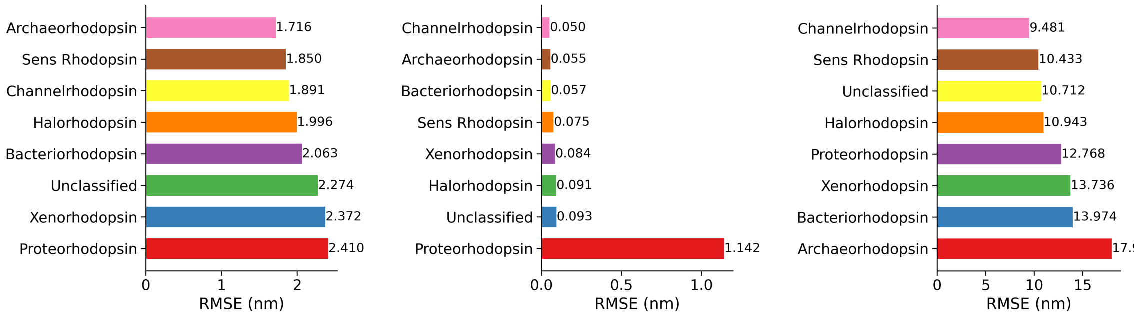

(d) One-hot encoding (e) XGBoost baseline (f) LASSO baseline

Figure CS6: **Architecture-selection and encoding-ablation diagnostics for the wavelength predictor.** (a) Multi-metric architecture comparison for physicochemical feature-map models. (b–c) Family-stratified RMSE and $R^2$ for the LASSO→XGBoost stacked model. (d) Family-stratified RMSE for the one-hot stacked model. (e–f) Family-stratified RMSE for XGBoost and LASSO baselines.

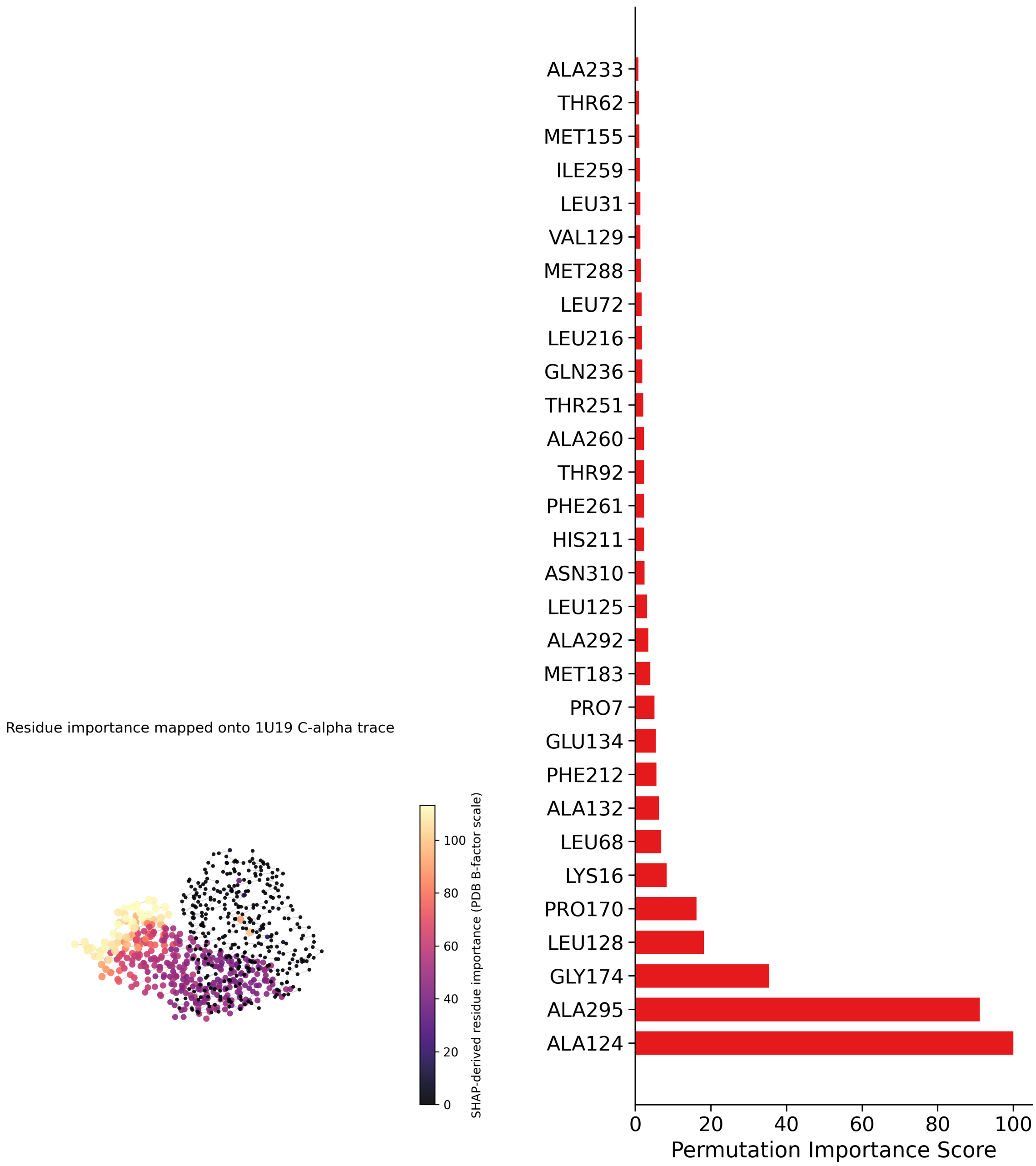

(a) Structural mapping

(b) Residue importance

Figure CS7: **Structural interpretability analysis for the wavelength predictor.** (a) C-alpha trace of 1U19 colored and scaled by SHAP-derived residue-importance values. (b) Bar chart of the highest-importance aligned positions. These panels support qualitative interpretation of the wavelength predictor rather than a mechanistic assignment of causal residues.

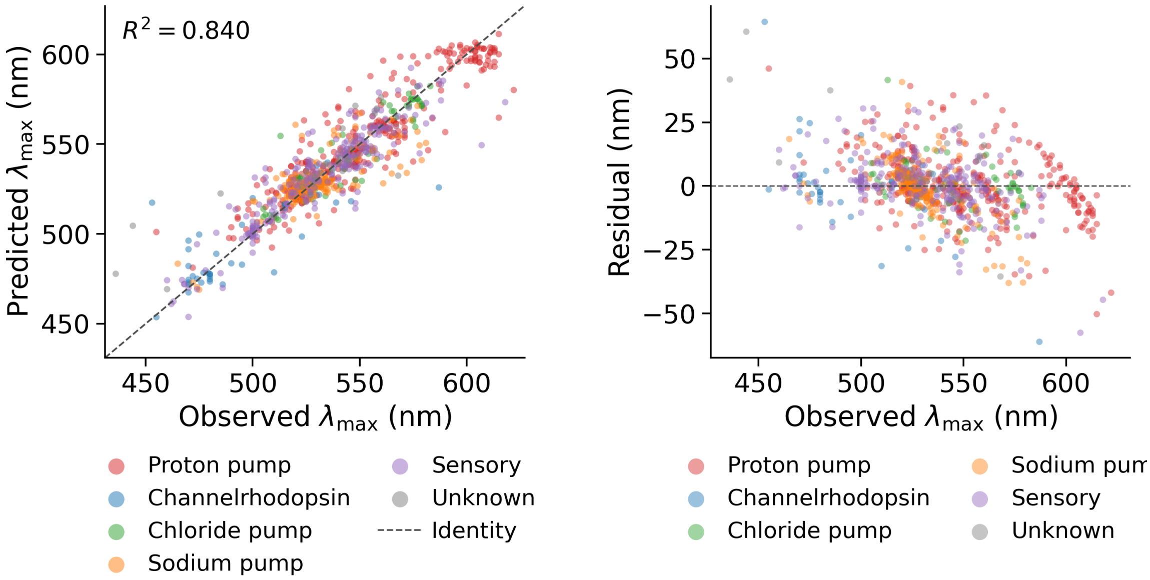

(a) Cross-validated predictions vs true $\lambda_{max}$ (b) Residuals vs true $\lambda_{max}$

Figure CS8: **Cross-validation predictions and residuals for the stacked $\lambda_{max}$ predictor.** (a) Predicted versus true $\lambda_{max}$ across all 10-fold cross-validation folds, coloured by functional class. (b) Residuals versus true $\lambda_{max}$; variance increases at the spectral extremes, consistent with the mild non-normality in the Q-Q plot.

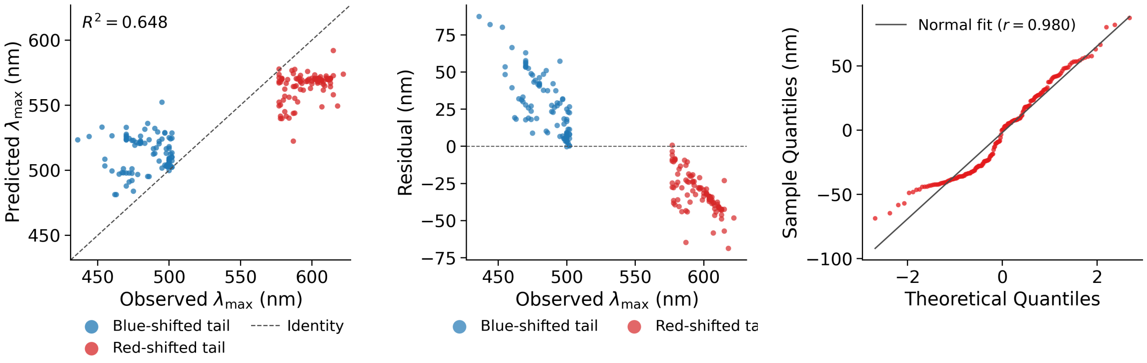

(a) Spectral-tail predictions (b) Spectral-tail residuals (c) Spectral-tail Q-Q plot

Figure CS9: **Out-of-distribution generalization: spectral-tail holdout experiment.** The model was trained on the central 90% of sequences ordered by $\lambda_{max}$ and evaluated on the held-out top and bottom 5% spectral tails. (a) Predicted versus true $\lambda_{max}$ for held-out spectral-tail sequences; compression toward the training mean is visible at extremes. (b) Residuals versus true $\lambda_{max}$; systematic bias at blue and red tails confirms out-of-distribution performance limits. (c) Q-Q plot for spectral-tail holdout; heavier tails than in-distribution residuals confirm reduced extrapolation accuracy.

##### 2.3 Full Sequence-Plausibility Filter Analysis

The main text gives the design-relevant summary of the Markov sequence-plausibility filter. The full benchmark, calibration, distributional, and mutation-plausibility analyses are provided below. Coarse rhodopsin-versus-decoy diagnostics include F1, MCC, AUROC, PR-AUC, mode-wise performance, and stacked score distributions (Supplementary Figs. CS10, CS11, CS12, CS13, CS14, and CS15). In the metric-sweep figures, panels (a–f) respectively show mean heatmaps, fold-variance heatmaps, hyperparameter line plots, top-architecture summaries, architecture–mode interactions, and mean–variance trade-offs; the mode-performance and score-distribution figures show sequence, structure, hybrid, and stacked panels in order (Supplementary Figs. CS10–CS15). Proton-pump subfamily diagnostics include the corresponding F1, MCC, AUROC, PR-AUC, mode-wise, and stacked-distribution panels, with the same panel ordering used to show the collapse of structure-only discrimination and the recovery of sequence-aware/stacked discrimination (Supplementary Figs. CS16, CS17, CS18, CS19, CS20, and CS21). Threshold calibration, wavelength neutrality, and mutation-distance acceptance are shown in Supplementary Table CS5, Supplementary Table CS6, and Supplementary Figs. CS22–CS25; these panels compare Markov, Levenshtein, CGR, and modality-specific acceptance curves for conservative versus radical proxy mutations.
Discriminating rhodopsin-like candidates from sequence decoys and off-target families is required for evolutionary engineering. We implemented a Markov-based filter that operates at four increasing levels of specificity:

- **Level 1 (Coarse):** Distinguish rhodopsins from non-rhodopsins (e.g., bacterial ion transporters). Target: AUROC $\geq 0.95$. Goal: Keep candidates in the rhodopsin-like sequence regime.
- **Level 2 (Fine):** Discriminate proton pumps from phylogenetically related subfamilies (chloride pumps, sodium pumps, sensory rhodopsins). Target: AUROC $\geq 0.90$. Goal: Keep the evolutionary search close to the target proton-pump lineage.
- **Level 3 (Calibration):** Verify wavelength neutrality - ensure the filter does not systematically reject blue-shifted candidates. Target: No correlation between filter score and $\lambda_{\text{max}}$. Goal: Enable unbiased filtering during evolution toward blue-shifted targets.
- **Level 4 (Mutation):** Distinguish synthetically generated conservative variants from radical-substitution mutants based on substitution chemistry and distance. Target: High selectivity ($\Delta_{sel} > 0$). Goal: Accept chemically conservative changes while rejecting mutation patterns expected to be less sequence-plausible.

We evaluated a Markov model across all four levels by varying threshold stringency. Operationally, the Markov score is a sequence likelihood under the proton-pump family model, so the filter is best interpreted as a family-membership or sequence-plausibility filter rather than a direct physical stability assay. This design is supported by the sequence-landscape stratification observed in Supplementary Figs. CS1 and CS2. The Minimum Spanning Tree analysis showed that functional classes occupy distinct regions of sequence space, with proton pumps segregated from related lineages. This separation motivates the four-level evaluation. Coarse filtering exploits the divergence between rhodopsins and non-rhodopsins. Fine filtering uses subfamily motifs that distinguish proton pumps from chloride pumps and sensory rhodopsins. Wavelength calibration tests whether filter score is coupled to absorption properties. Mutation-distance filtering tests whether the score discriminates chemically conservative from radical synthetic substitutions across mutation distances. We tested this filtering pipeline on progressively challenging discrimination tasks, beginning with the coarse-grained problem:

We benchmarked four architectures: Markov Models, Chaos Game Representation (CGR), Alignment-based scoring, and Levenshtein distance. Normalized score distributions (Fig. CS15a–CS15d) showed clear separation across multiple feature modalities. Levenshtein sequence models achieved high median scores (F1=0.989, AUROC=1.000), confirming that global sequence homology serves as an effective discriminator when decoys are phylogenetically distant. The Markov model's structure-based modality also performed well (median AUROC 0.980), reflecting the distinct statistical signatures of rhodopsin transmembrane bundles compared to other transporters.

To capture specific design limits, Markov models were constructed by defining transition probabilities $P(X_i|X_{i-1})$ from a multiple sequence alignment. Probabilities were smoothed using a Dirichlet prior ($\alpha = 0.6$) to allow for chemically similar but unobserved transitions. This generative approach scores the grammatical correctness of a sequence based on the underlying rhodopsin syntax. Quantitative analysis (Table CS3) shows that the Stacked Markov Model achieves high performance under the random sequence-level split (median F1=$0.980 \pm 0.022$), occupying the high-precision quadrant in benchmark analysis (Fig. CS10).

We then added a second validation step–subfamily detection–to test whether the evolutionary search could be constrained near the target lineage. This analysis tested subfamily specificity: can we distinguish the target proton pump family from phylogenetically related but functionally distinct subfamilies (chloride pumps, sodium pumps, and sensory rhodopsins)?

Score distributions (Fig. CS21) showed distinct peaks for proton pumps, with the Stacked Markov model achieving clear segregation (proton pumps $> 0.8$, off-targets $< 0.1\%$). Quantitative benchmarking (Table CS4) showed strong median performance for Stacked Markov (AUROC=1.000, F1=0.984) and Levenshtein Structure (AUROC=1.000, F1=0.944) under the random sequence-level split protocol. A critical divergence emerged for the structural modality in Markov models: unlike the coarse filter, the Markov Structure model collapses on this task (median AUROC 0.489; Table CS4). Gross secondary structural topology was highly similar between proton pumps and chloride pumps, rendering structure-only models insensitive to the sequence features used for this lineage distinction (Fig. CS20). The Stacked Markov model captured sequence-transition motifs associated with ion-specific lineages, reducing the chance that the evolutionary search drifts into chloride-pump-like regions. The bar charts (Fig. CS20) show this contrast: structure-only models (b) fail while Stacked models (d) maintain high median AUROC, AUPRC, and F1 scores.

We calibrated the trade-off between positive-class retention (Recall) and decoy rejection (Specificity) across varying filter strengths to identify the filter's operational window (Table CS5). The results revealed a critical operational regime: coarse-grained filtration maintained high specificity at low strengths (TNR $> 94\%$ at strength 0.01), but subfamily discrimination required tighter thresholds (0.10-0.25) to achieve meaningful specificity (TNR $> 85\%$) without discarding too many proton-pump-labeled sequences.

Sequence-aware ensemble models achieved high specificity when comparing proton-pump-labeled sequences to off-target lineages under random sequence-level splits (Table CS4), supporting their use as lineage filters during GA search. Before proceeding to mutation-level filtering, we verified that the filter does not introduce systematic bias against specific wavelengths.

#### Wavelength calibration

To ensure the sequence-plausibility filter does not inadvertently reject blue-shifted candidates (our design target), we analyzed filter acceptance rates stratified by absorption wavelength ($\lambda_{\max}$). We plotted the filter's anomaly probability against $\lambda_{\max}$ for all proton pumps in the validation set, coloring proton pumps (red) separately from other rhodopsin families (gray) (Fig. CS22).

The scatter plots show no systematic correlation between filter score and wavelength for proton pumps. Blue-shifted proton pumps ($\lambda_{\max} < 500$ nm) and red-shifted ones ($\lambda_{\max} > 550$

Table CS3: **Performance of Coarse-Grained Rhodopsin Filters.** Metrics are median ± fold SD from random sequence-level 10-fold cross-validation for discriminating rhodopsins ($n = 884$) from non-rhodopsin ion transporters ($n = 10,000$). Levenshtein Sequence achieves median AUROC = 1.000, PR-AUC = 1.000, and F1 = 0.989 under this split protocol.

| Architecture | Modality | AUROC | PR-AUC | F1-Score |
|---|---|---|---|---|
| **Levenshtein** | **Sequence** | **1.000 ± 0.001** | **1.000 ± 0.001** | **0.989 ± 0.011** |
| | Structure | 1.000 ± 0.002 | 1.000 ± 0.002 | 0.989 ± 0.019 |
| **Markov Model** | **Stacked** | **0.999 ± 0.007** | **0.999 ± 0.007** | **0.980 ± 0.022** |
| | Structure | 0.980 ± 0.024 | 0.966 ± 0.055 | 0.960 ± 0.035 |
| Alignment | Hybrid | 1.000 ± 0.007 | 1.000 ± 0.005 | 0.989 ± 0.032 |
| | Sequence | 0.917 ± 0.094 | 0.960 ± 0.045 | 0.679 ± 0.094 |
| CGR | Hybrid | 0.896 ± 0.122 | 0.924 ± 0.095 | 0.799 ± 0.151 |

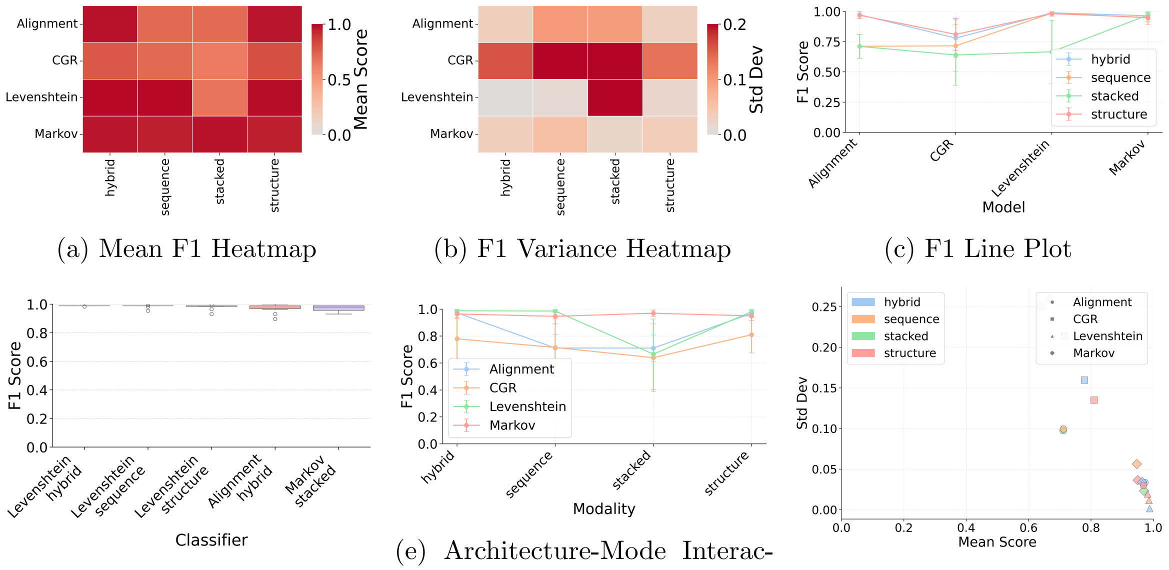

(a) Mean F1 Heatmap (b) F1 Variance Heatmap (c) F1 Line Plot

(d) Top Architectures (F1) (e) Architecture-Mode Interaction (f) Mean vs Variance (F1)

Figure CS10: **Hyperparameter stability analysis for F1 score.** (a) Heatmap of mean F1 scores across architecture-modality combinations. (b) Heatmap of F1 variance across cross-validation folds. (c) Line plot of F1 vs hyperparameter setting for each architecture. (d) Bar chart of top-performing architectures by median F1. (e) Scatter plot of architecture-mode interaction effects. (f) Mean vs variance scatter: best models cluster in upper-right (high F1, low variance).

Table CS4: **Subfamily Specificity Breakdown.** Discrimination of Proton Pumps ($n = 310$) from phylogenetically related families (Chloride Pumps, Sensory Rhodopsins; $n = 296$) under random sequence-level cross-validation. Markov Stacked achieves median AUROC = 1.000 and F1 = 0.984 under this split protocol. Markov Structure fails (median AUROC = 0.489).

| Architecture | Modality | AUROC | PR-AUC | F1-Score |
|---|---|---|---|---|
| **Markov Model** | **Stacked** | **1.000 ± 0.157** | **1.000 ± 0.119** | **0.984 ± 0.165** |
| Markov Model | Structure | 0.489 ± 0.293 | 0.490 ± 0.241 | 0.618 ± 0.249 |
| Levenshtein | Structure | 1.000 ± 0.035 | 1.000 ± 0.024 | 0.944 ± 0.056 |
| Levenshtein | Sequence | 0.986 ± 0.058 | 0.991 ± 0.052 | 0.967 ± 0.091 |

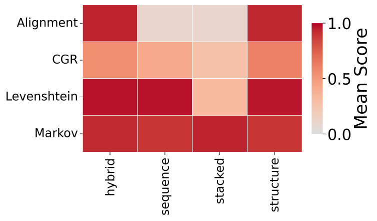

(a) Mean MCC Heatmap

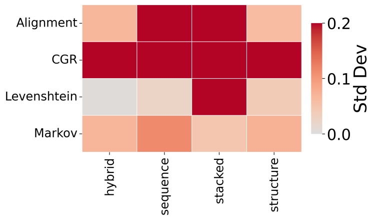

(b) MCC Variance Heatmap

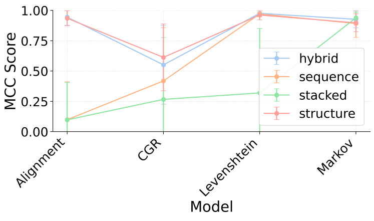

(c) MCC Line Plot

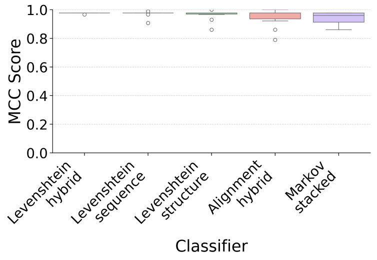

(d) Top Architectures (MCC)

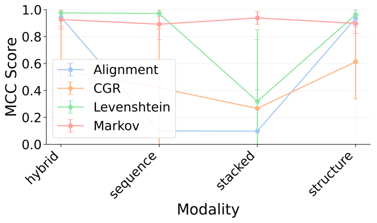

(e) Architecture-Mode Interaction

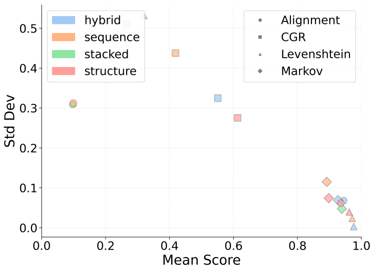

(f) Mean vs Variance (MCC)

Figure CS11: **Evaluation of balanced discrimination via Matthews Correlation Coefficient (MCC).** (a) Heatmap of mean MCC across architecture-modality combinations. (b) MCC variance heatmap. (c) Line plot of MCC vs hyperparameter. (d) Bar chart of top architectures by median MCC. (e) Architecture-mode interaction scatter. (f) Mean vs variance scatter: sequence-based models achieve MCC ¿ 0.95, indicating balanced classification beyond majority baseline.

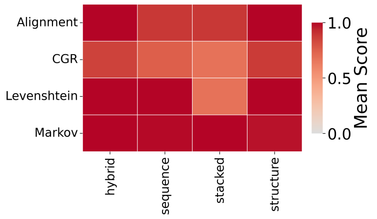

(a) Mean ROC AUC Heatmap

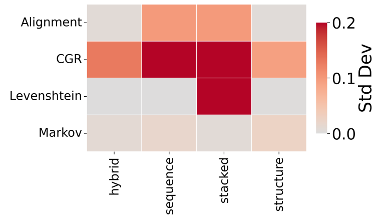

(b) ROC AUC Variance Heatmap

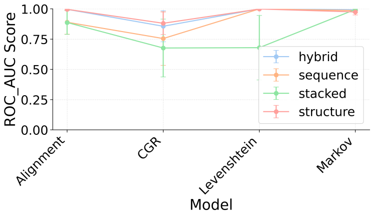

(c) ROC AUC Line Plot

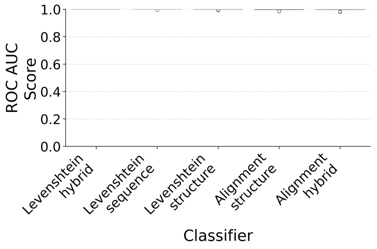

(d) Top Architectures (ROC AUC)

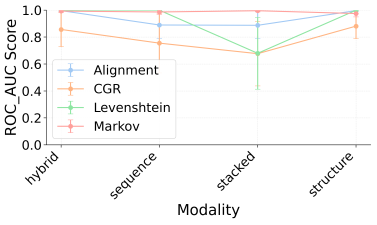

(e) Architecture-Mode Interaction

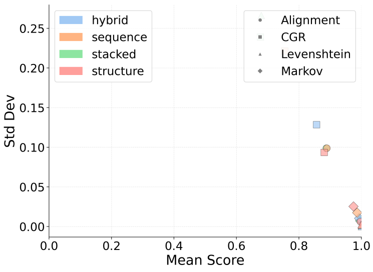

(f) Mean vs Variance (ROC AUC)

Figure CS12: **Global discrimination performance (ROC AUC).** (a) Heatmap of mean AUROC across architectures and modalities. (b) AUROC variance heatmap. (c) Line plot of AUROC vs hyperparameter. (d) Bar chart of top architectures by median AUROC. (e) Architecture-mode interaction effects. (f) Mean vs variance scatter: sequence-based models achieve AUROC $\approx$ 1.0, indicating near-perfect separation of rhodopsins from decoys.

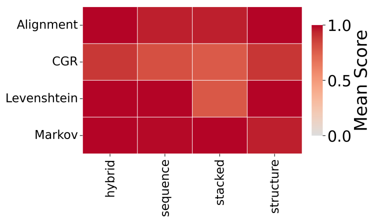

(a) Mean PR AUC Heatmap

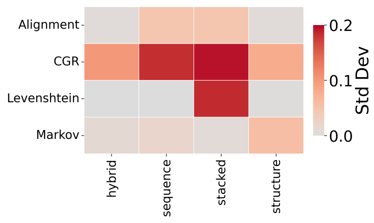

(b) PR AUC Variance Heatmap

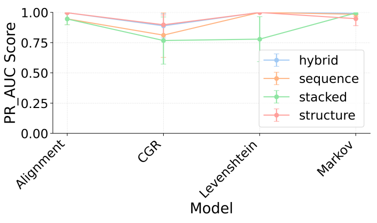

(c) PR AUC Line Plot

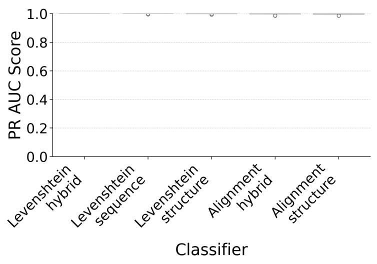

(d) Top Architectures (PR AUC)

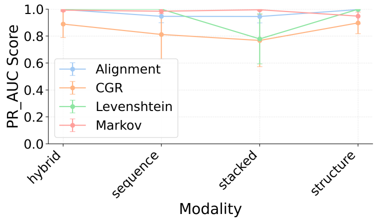

(e) Architecture-Mode Interaction

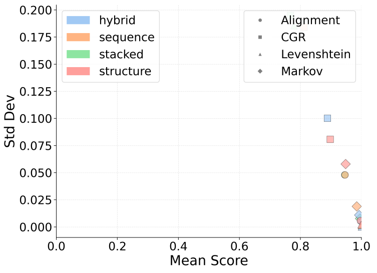

(f) Mean vs Variance (PR AUC)

Figure CS13: **Precision-Recall Analysis (PR AUC).** (a) Heatmap of mean PR AUC. (b) PR AUC variance heatmap. (c) Line plot of PR AUC vs hyperparameter. (d) Bar chart of top architectures by median PR AUC. (e) Architecture-mode interaction effects. (f) Mean vs variance scatter: best models cluster at high PR AUC with low variance, maintaining low FPR at high recall.

Table CS5: **Filter Strength Calibration.** FPR, FNR, TPR, TNR, and F1 scores across filter strength values (0.01–0.30), where strength sweeps the FNR. At strength 0.01, coarse filter achieves TNR = 0.942, FNR = 0.010. Subfamily filter requires higher strength (0.10–0.25) to achieve TNR ¿ 0.86.

| **Task** | **Strength** | **FPR** | **FNR** | **TPR** | **TNR** | **F1** |
|---|---|---|---|---|---|---|
| **Coarse (Rho vs Non)** | 0.01 | 0.0576 | 0.0102 | 0.9898 | 0.9424 | 0.9685 |
| | 0.05 | 0.0216 | 0.0509 | 0.9491 | 0.9784 | 0.9638 |
| | 0.10 | 0.0120 | 0.1007 | 0.8993 | 0.9880 | 0.9414 |
| | 0.15 | 0.0108 | 0.1505 | 0.8495 | 0.9892 | 0.9136 |
| | 0.20 | 0.0072 | 0.2002 | 0.7998 | 0.9928 | 0.8854 |
| | 0.25 | 0.0060 | 0.2500 | 0.7500 | 0.9940 | 0.8544 |
| | 0.30 | 0.0036 | 0.2998 | 0.7002 | 0.9964 | 0.8220 |
| **Subfamily (Proton vs Other)** | 0.01 | 0.7697 | 0.0100 | 0.9900 | 0.2303 | 0.7070 |
| | 0.05 | 0.7697 | 0.0498 | 0.9502 | 0.2303 | 0.6883 |
| | 0.10 | 0.1388 | 0.0997 | 0.9003 | 0.8612 | 0.8799 |
| | 0.15 | 0.0473 | 0.1495 | 0.8505 | 0.9527 | 0.8951 |
| | 0.20 | 0.0284 | 0.1993 | 0.8007 | 0.9716 | 0.8748 |
| | 0.25 | 0.0095 | 0.2492 | 0.7508 | 0.9905 | 0.8528 |
| | 0.30 | 0.0000 | 0.2990 | 0.7010 | 1.0000 | 0.8242 |

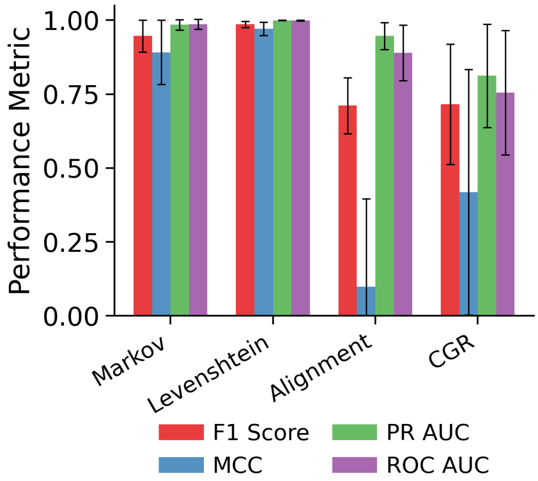

(a) Sequence Modality

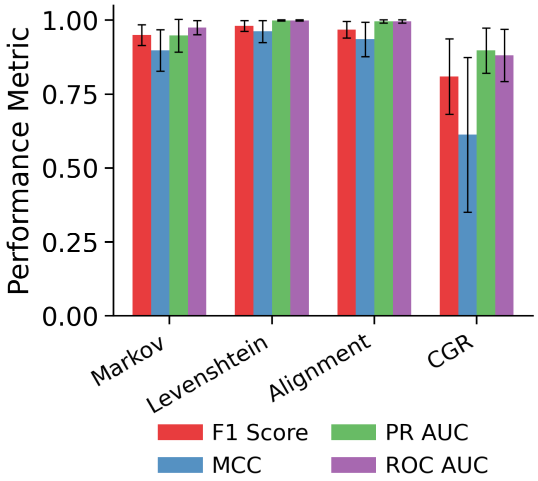

(b) Structure Modality

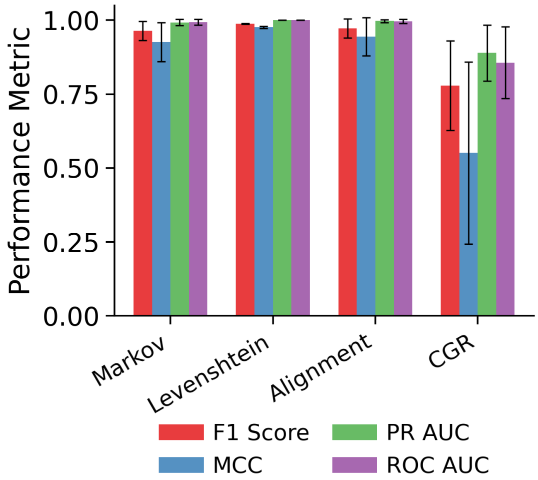

(c) Hybrid Modality

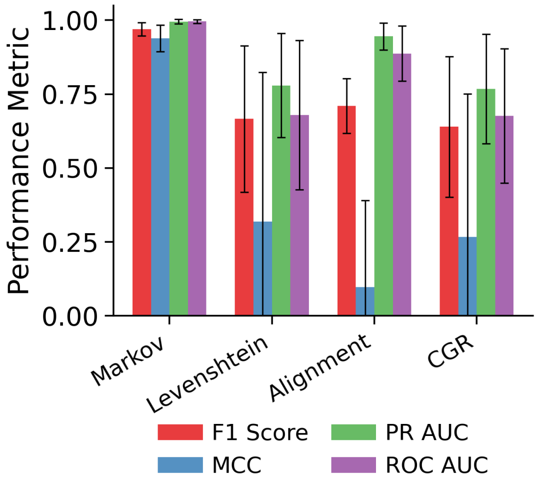

(d) Stacked Modality

Figure CS14: **Model performance across input modalities.** (a-d) Bar charts compare three cross-validated metrics (AUROC, AUPRC, F1) for Sequence, Structure, Hybrid, and Stacked input modalities. Sequence-only inputs use raw amino acid features; Structure inputs use predicted secondary structure; Hybrid combines both feature sets; Stacked ensembles the three models. All metrics are from 10-fold cross-validation.

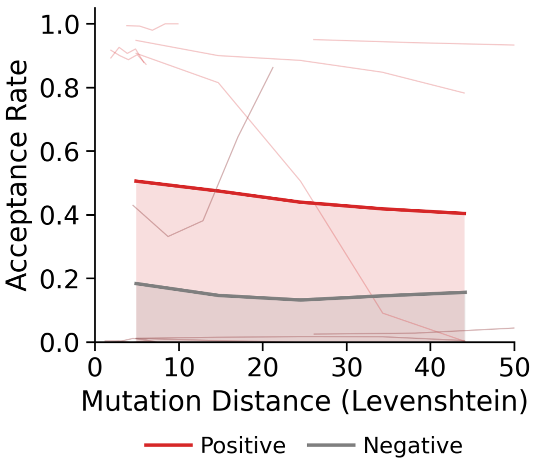

(a) Sequence Distributions

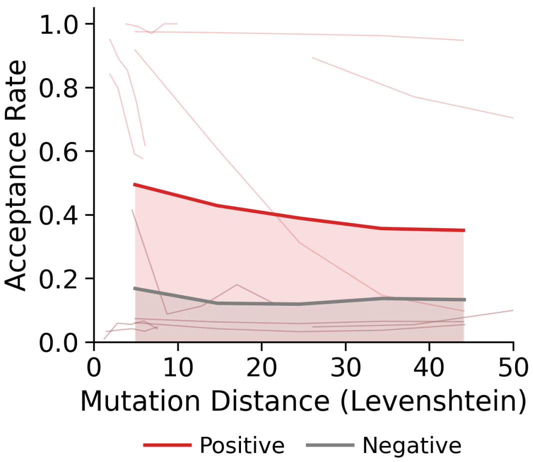

(b) Structure Distributions

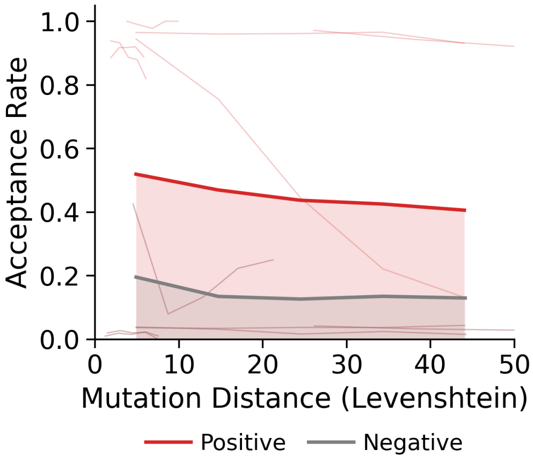

(c) Hybrid Distributions

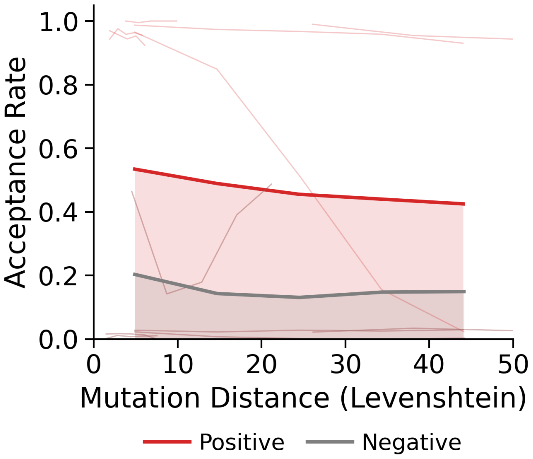

(d) Stacked Distributions

Figure CS15: **Verification of decision boundaries for coarse filtering.** (a) Sequence modality: rhodopsin-positive (green) and decoy (red) distributions. (b) Structure modality distributions. (c) Hybrid modality distributions. (d) Stacked modality: minimal overlap between positive and decoy classes, providing clearest separation.

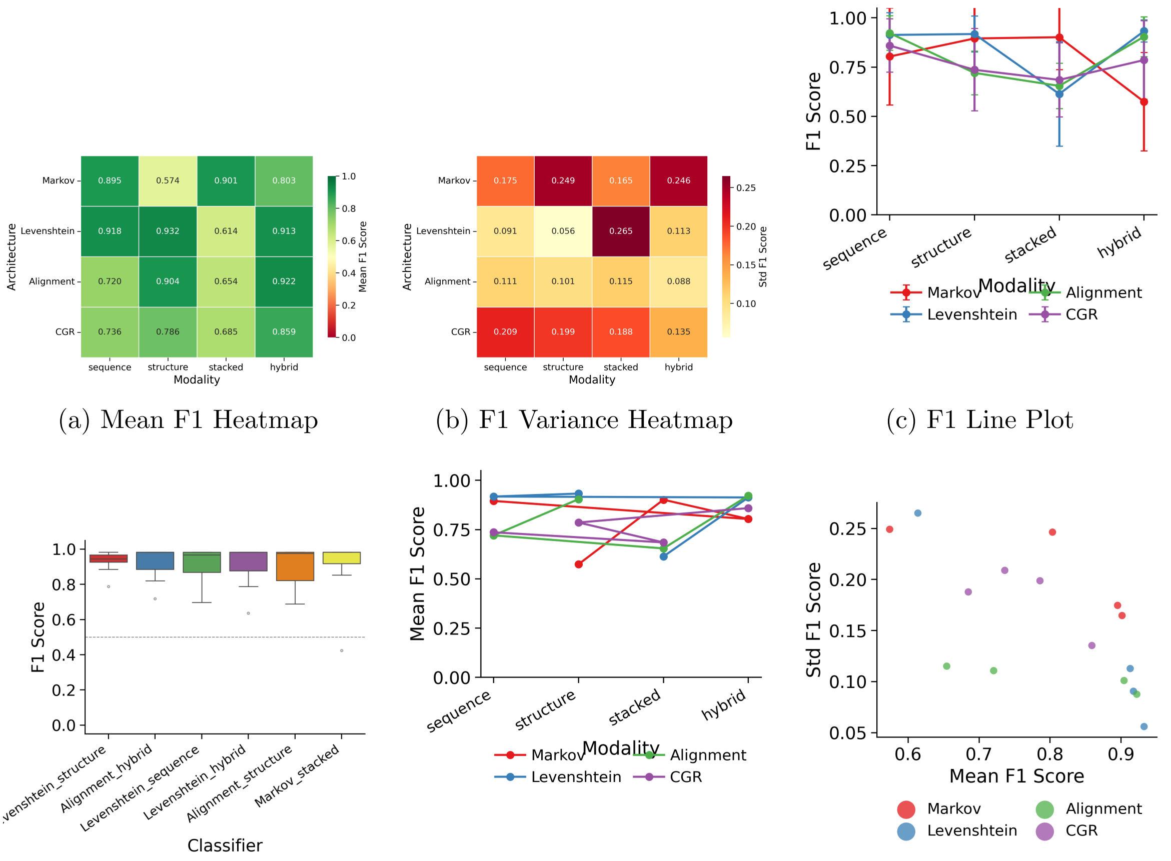

(a) Mean F1 Heatmap

(b) F1 Variance Heatmap

(c) F1 Line Plot

(d) Top Architectures (F1)

(e) Architecture-Mode Interaction

(f) Mean vs Variance (F1)

Figure CS16: **Subfamily Specificity: Hyperparameter analysis for F1 score.** (a) Mean F1 heatmap for subfamily discrimination (proton pumps vs related families). (b) F1 variance heatmap. (c) Line plot of F1 vs hyperparameter. (d) Bar chart of top architectures. (e) Architecture-mode interaction. (f) Mean vs variance: Stacked and Levenshtein models achieve high F1 despite phylogenetic proximity of decoys.

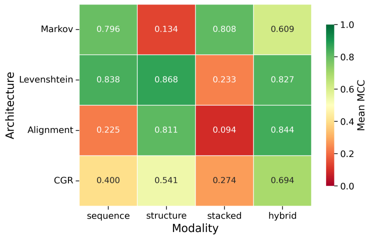

(a) Mean MCC Heatmap

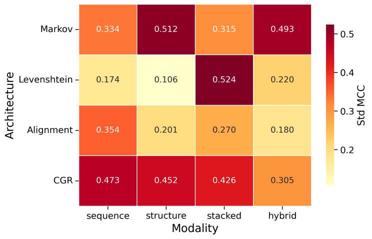

(b) MCC Variance Heatmap

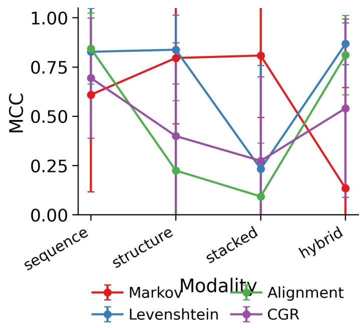

(c) MCC Line Plot

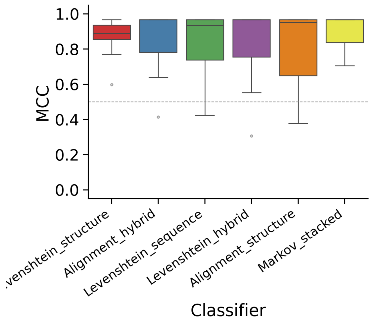

(d) Top Architectures (MCC)

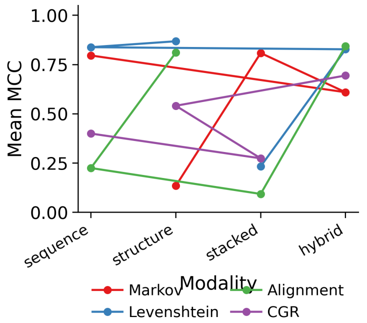

(e) Architecture-Mode Interaction

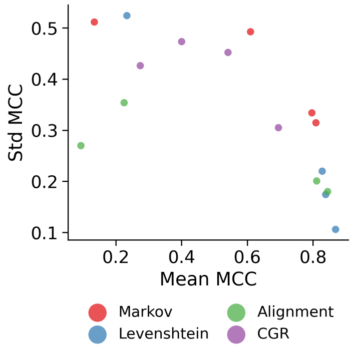

(f) Mean vs Variance (MCC)

Figure CS17: **Subfamily Specificity: Matthews Correlation Coefficient analysis.** (a) Mean MCC heatmap for subfamily discrimination. (b) MCC variance heatmap. (c) Line plot vs hyperparameter. (d) Bar chart of top architectures. (e) Architecture-mode interaction. (f) Mean vs variance scatter: sequence-based models show consistent high MCC while structure-only models collapse.

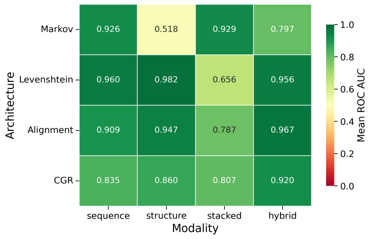

(a) Mean ROC AUC Heatmap

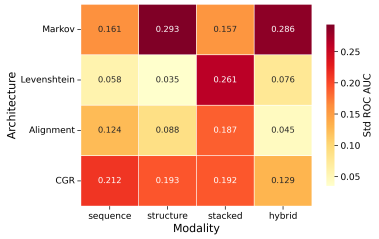

(b) ROC AUC Variance Heatmap

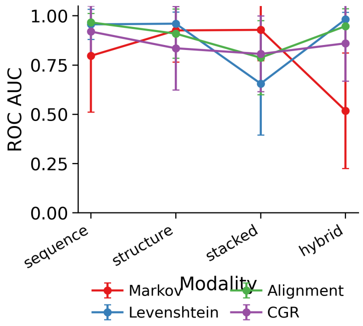

(c) ROC AUC Line Plot

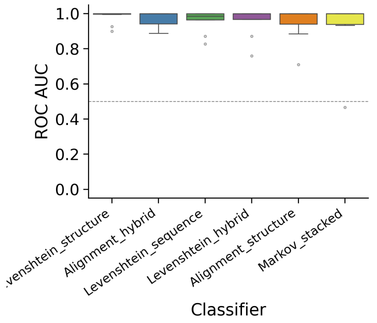

(d) Top Architectures (ROC AUC)

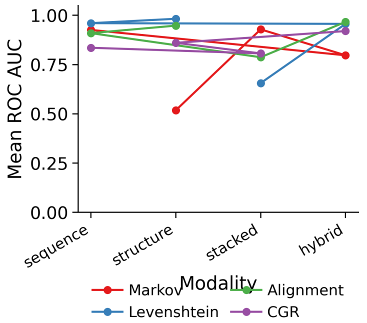

(e) Architecture-Mode Interaction

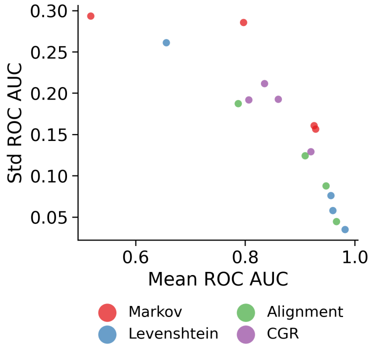

(f) Mean vs Variance (ROC AUC)

Figure CS18: **Subfamily Specificity: ROC AUC analysis.** (a) Mean AUROC heatmap. (b) AUROC variance heatmap. (c) Line plot vs hyperparameter. (d) Bar chart of top architectures. (e) Architecture-mode interaction. (f) Mean vs variance scatter: sequence-aware models reach high median AUROC, detecting sequence signatures that distinguish proton pumps from related families.

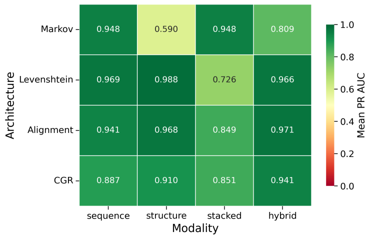

(a) Mean PR AUC Heatmap

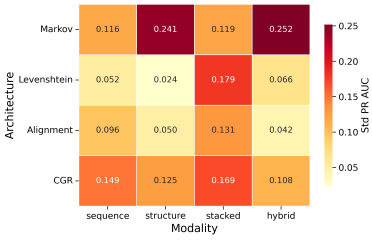

(b) PR AUC Variance Heatmap

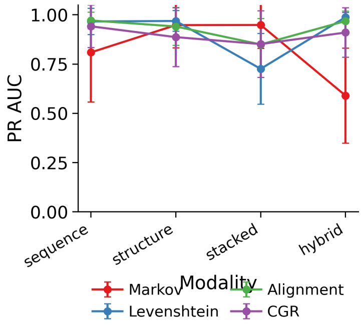

(c) PR AUC Line Plot

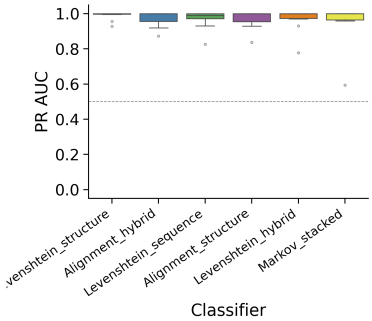

(d) Top Architectures (PR AUC)

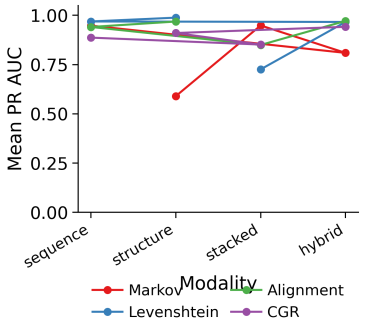

(e) Architecture-Mode Interaction

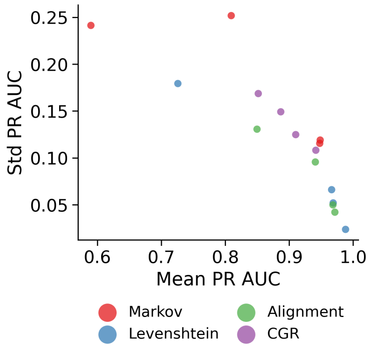

(f) Mean vs Variance (PR AUC)

Figure CS19: **Subfamily Specificity: PR AUC analysis.** (a) Mean PR AUC heatmap. (b) PR AUC variance heatmap. (c) Line plot vs hyperparameter. (d) Bar chart of top architectures. (e) Architecture-mode interaction. (f) Mean vs variance scatter: Stacked and Levenshtein models maintain low false positive rates with minimal chloride-pump misclassification.

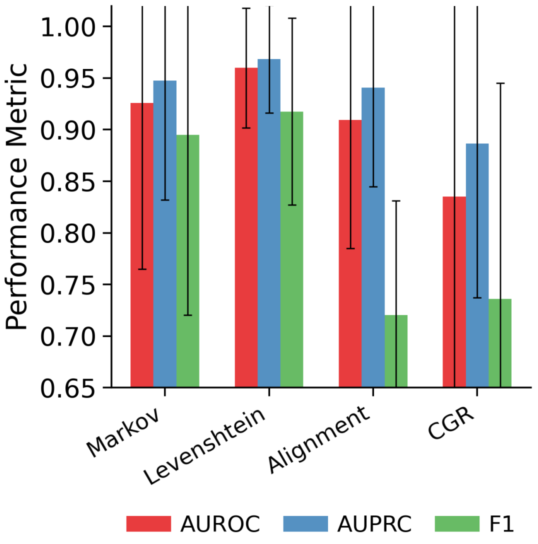

(a) Sequence Modality

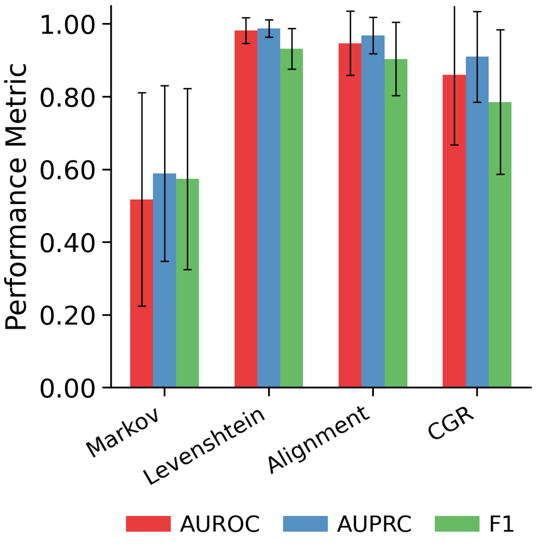

(b) Structure Modality

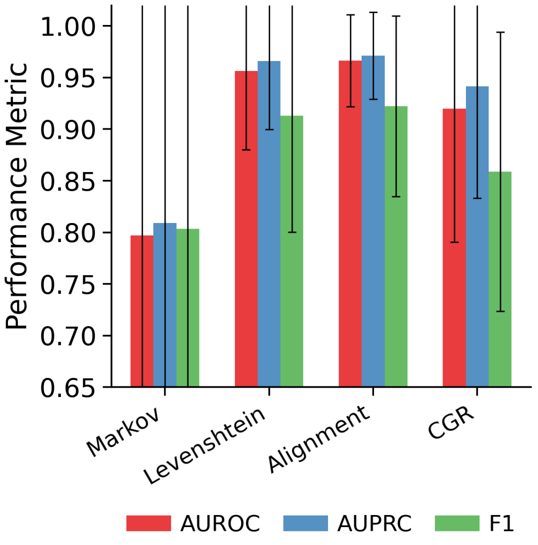

(c) Hybrid Modality

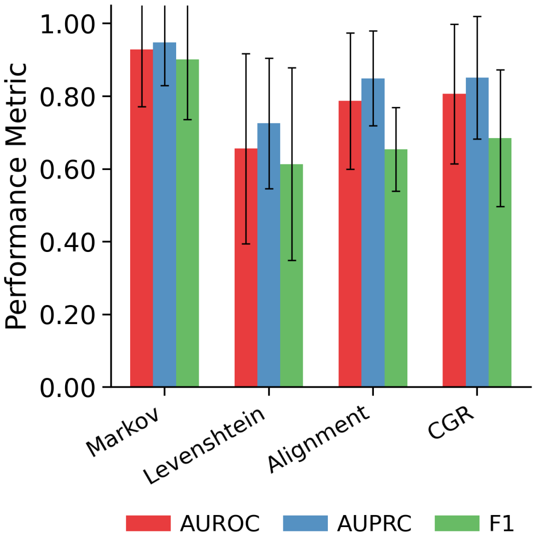

(d) Stacked Modality

Figure CS20: **Subfamily Specificity: Performance by modality.** (a-d) Bar charts comparing AUROC, AUPRC, and F1 scores for Sequence, Structure, Hybrid, and Stacked input modalities when discriminating proton pumps from related families. Structure-only models (b) fail at this task, while Stacked models (d) maintain high performance.

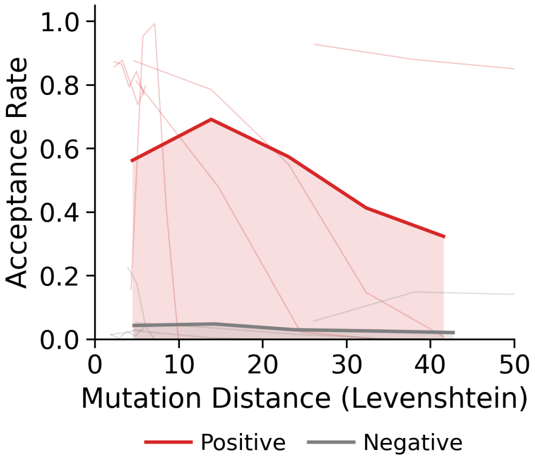

(a) Sequence Distributions

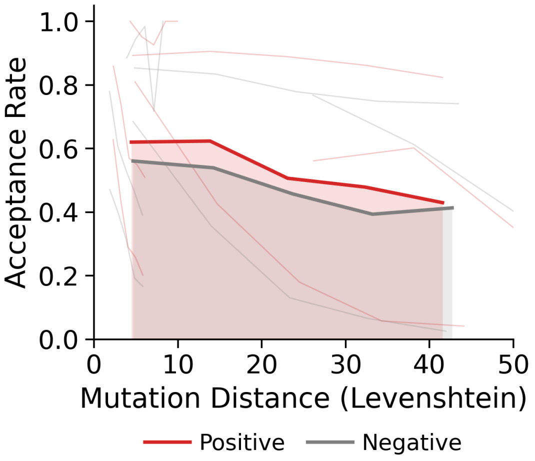

(b) Structure Distributions

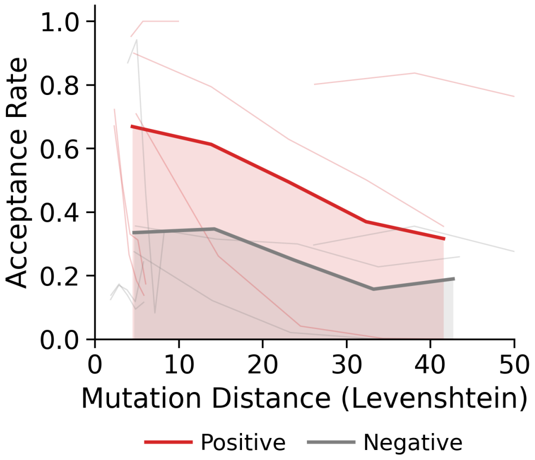

(c) Hybrid Distributions

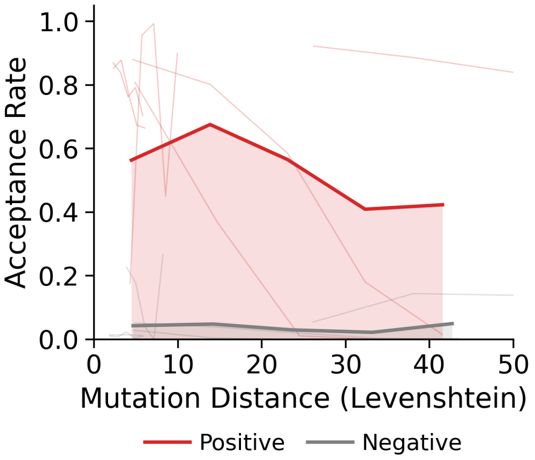

(d) Stacked Distributions

Figure CS21: **Subfamily Specificity: Decision boundary analysis.** (a) Sequence modality: proton pumps (green) vs related families (red). (b) Structure modality: high overlap between proton pumps and chloride pumps. (c) Hybrid modality distributions. (d) Stacked modality: clear separation between proton pumps and other families, achieving median AUROC 1.000.

nm) both span the full range of filter scores. This indicates that the Markov model score is approximately wavelength-neutral within the curated proton-pump set. During GA-guided evolution toward blue targets, rejection is therefore not directly driven by the wavelength objective. Non-proton-pump families (gray points) cluster at lower acceptance probabilities, consistent with the subfamily discrimination results above.

We quantified this wavelength neutrality using linear regression of filter score against $\lambda_{\max}$. For all architectures tested, the $R^2$ values are near zero (Table CS6), indicating no systematic relationship between filter performance and spectral properties. The Markov Stacked model achieves $R^2 = 0.002$ for proton pumps (focus set), supporting approximately wavelength-neutral filtering within the curated proton-pump set.

Having observed approximate wavelength neutrality within the curated proton-pump set, we proceed to the final filtering level: testing whether the filter distinguishes conservative synthetic mutants from radical-substitution mutants.

#### Mutation-distance stratification and selectivity

After excluding non-rhodopsin decoys, we evaluated the models' ability to distinguish conservative synthetic variants from radical-substitution mutants. Directed evolution requires filtering that remains permissive to chemically conservative changes while rejecting mutation patterns expected to be less plausible under the proxy definition. We tested this using a synthetic library of 10,000 rhodopsin mutants, divided by mutation distance and substitution type. The positive synthetic class contained conservative substitutions within amino-acid property groups, while the negative synthetic class contained non-conservative substitutions across property groups. These labels are chemical proxies for plausibility rather than direct experimental measurements of function.

We defined the ideal performance across three evolutionary stages:

- **Low Mutation Distance:** High acceptance of conservative variants for local optimization, but reduced acceptance of radical variants.
- **Medium Mutation Distance:** Non-zero acceptance of conservative variants to allow path traversal, with near-zero acceptance of radical variants.
- **High Mutation Distance:** Rejection of all variants to bound the search space.

At Low Mutation Distance (Bin 1, $\bar{d} \approx 5$), the architectures diverged sharply (Fig. CS24). The Markov model family showed consistent performance across modalities (Sequence, Structure, Stacked). Every Markov variant accepted 40-60% of conservative synthetic variants while suppressing radical variants. The Stacked Markov Model led this group with a positive margin ($\Delta_{sel} = +17.7\%$), indicating that probabilistic transition statistics capture local sequence constraints. By modeling the probability of residue $i$ given its context $(i-1, i-2, \dots)$, the Markov approach detected transitions that are rare under the learned local sequence model, even when global sequence identity remains high.

In contrast, all other architectures failed this initial proxy test. The Levenshtein models exhibited an inverse behavior, accepting fewer conservative variants than radical variants ($\Delta_{sel} = -14.5\%$). The CGR models were indiscriminate, failing to separate the classes meaningfully ($\Delta_{sel} \approx +2\%$). The Alignment models, while selective, were overly restrictive, rejecting $> 85\%$ of conservative variants and limiting local exploration. This behavior reflects a limitation of distance-based metrics: they treat all substitutions equally. A conservative mutation (e.g., Leucine to Isoleucine) has the same "distance cost" as a radical one (e.g., Leucine to Proline), despite very different expected structural implications. The Markov model, trained on natural transition frequencies, captures this distinction by favoring common transitions such as L→I and penalizing rare transitions such as L→P in helical contexts.

At Medium Mutation Distance (Bin 3, $\bar{d} \approx 24$), distance-based metrics failed. Levenshtein, Alignment, and CGR models all converged to near-zero acceptance ($< 0.2\%$) for conservative

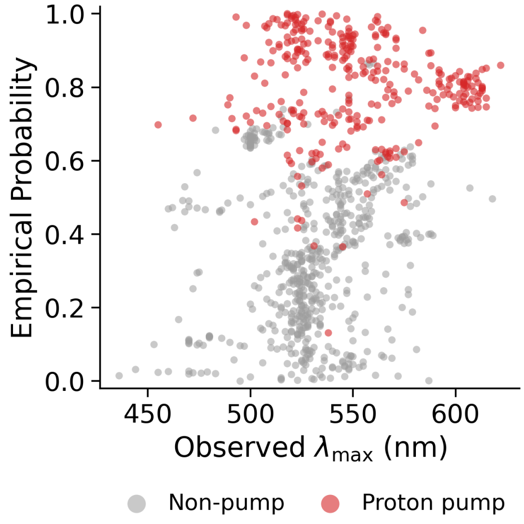

(a) Markov Stacked

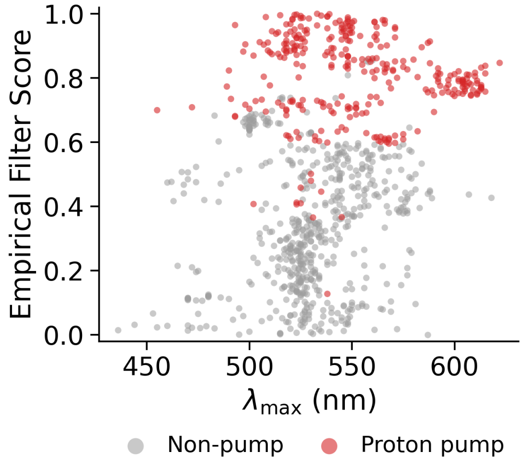

(b) Levenshtein Stacked

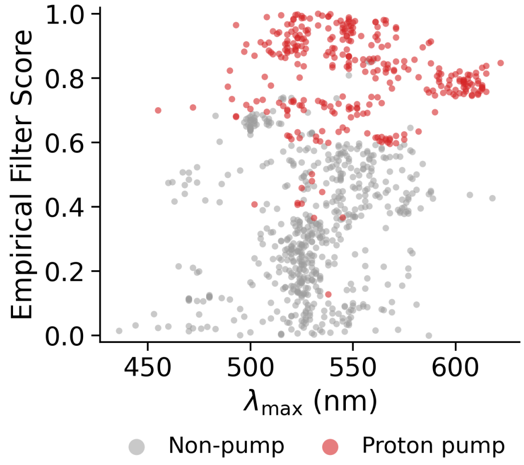

(c) Markov Sequence

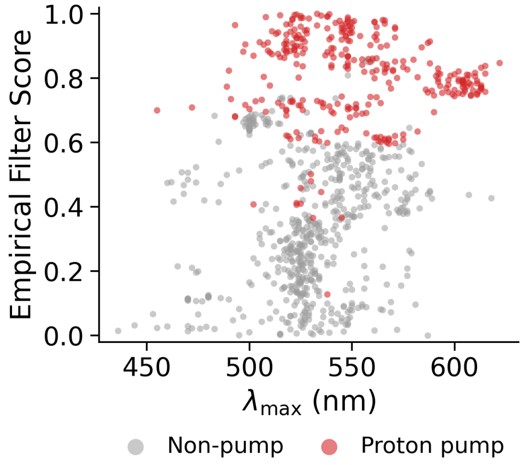

(d) Levenshtein Sequence

Figure CS22: **Wavelength calibration of the sequence-plausibility filter.** (a-d) Filter anomaly probability vs $\lambda_{max}$ for proton pumps only, showing approximately wavelength-neutral filtering across architectures. Points span the full probability range at all wavelengths, with $R^2 < 0.05$ for all models. This indicates that the filter score is not strongly coupled to spectral properties within the curated set.

variants. This premature rejection limited evolutionary progress. Uniquely, the Markov architecture maintained a sequence-plausible path. It retained a 9.3% conservative-variant acceptance rate while filtering radical mutants (0.09%). This capacity to discriminate at intermediate distances allows the algorithm to traverse the search landscape.

At High Mutation Distance (Bin 5, $\bar{d} \approx 41$), all models succeeded in the final criterion: bounding the search space. Every architecture converged to a hard rejection regime ($> 99\%$ rejection), preventing drift into low-scoring sequence regions outside the learned proton-pump-like distribution (Fig. CS23, CS24).

Quantitative benchmarking across all models (Fig. CS23 and CS24) supported use of the Stacked Markov Model as the sequence-plausibility filter. This model maximized the Area Under the Acceptance Curve (AUC) for conservative variants (≈21.3) while suppressing distinct families (≈8.9) and radical mutants (≈11.9). In comparison, structure-based models often failed to reject off-target families (AUC ≈19.3), while alignment-based approaches were overly restrictive for all variants. Because the Markov architecture was the only framework to succeed across all three proxy regimes–avoiding the inversion of Levenshtein, the looseness of CGR, and the restrictiveness of Alignment–we selected the Stacked Markov Model for sequence-plausibility filtering. Applied in practice, this filter reduces the candidate pool by an order of magnitude compared with an unconstrained GA, focusing experimental effort on a compact set of candidates that remain closer to the proton-pump-like sequence distribution.

We plotted acceptance rates across the full mutation trajectory for representative models to visualize these divergent behaviors (Fig. CS25). The Stacked Markov Model (panel a) and Levenshtein Structure (panel b) showed the target gatekeeper profile: high acceptance of conservative variants at low distance, decaying as distance increases, with consistently low acceptance of radical variants. In contrast, models like Levenshtein Stacked (panel d) and CGR Stacked (panel e) exhibited indistinguishable or inverted curves, confirming their inability to separate the proxy classes.

Table CS6: **Wavelength Calibration: $R^2$ of Filter Score vs $\lambda_{\max}$.** Regression coefficients measure correlation between filter anomaly probability and absorption wavelength. Near-zero $R^2$ values ($< 0.05$) indicate wavelength-neutral filtering. Lower values are better.

| Architecture | Mode | Focus Set ($R^2$) | Other Fam. ($R^2$) | All Data ($R^2$) |
|---|---|---|---|---|
| **Markov Stacked** | **Stacked** | **0.0023** | **0.0156** | **0.0627** |
| Markov Sequence | Sequence | 0.0023 | 0.0146 | 0.0605 |
| Levenshtein Stacked | Stacked | 0.0050 | 0.0001 | 0.0025 |
| Levenshtein Sequence | Sequence | 0.0436 | 0.0001 | 0.1103 |
| Markov Hybrid | Hybrid | 0.0039 | 0.0536 | 0.0797 |

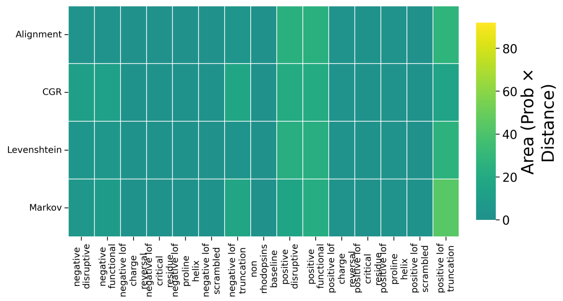

(a) Sequence Modality

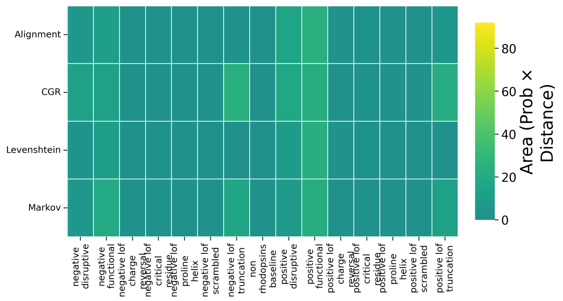

(b) Structure Modality

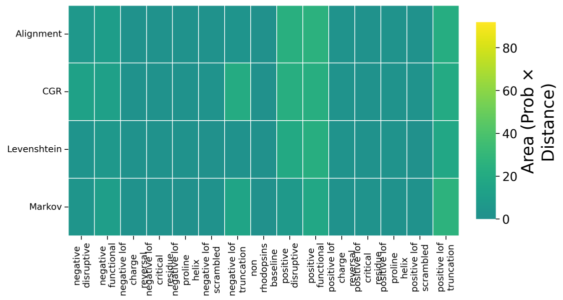

(c) Hybrid Modality

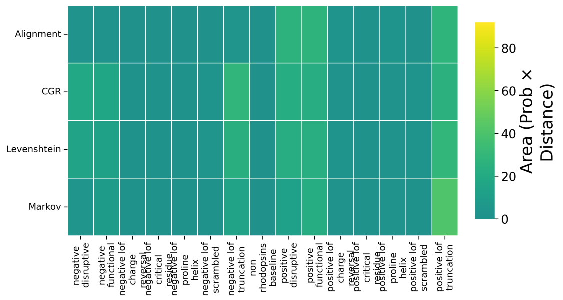

(d) Stacked Modality

Figure CS23: **Selectivity analysis via Area Under Acceptance Curve (AUC).** (a) Sequence modality: conservative (solid) vs radical-substitution (dashed) synthetic variant acceptance. (b) Structure modality. (c) Hybrid modality. (d) Stacked modality: AUC $\approx 21.3$ for conservative variants, $\approx 11.9$ for radical variants (high selectivity). Levenshtein models show inverted selectivity.

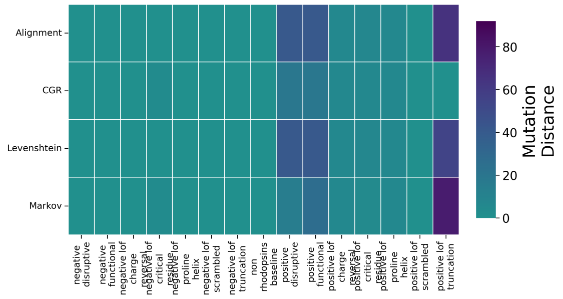

(a) Sequence Modality

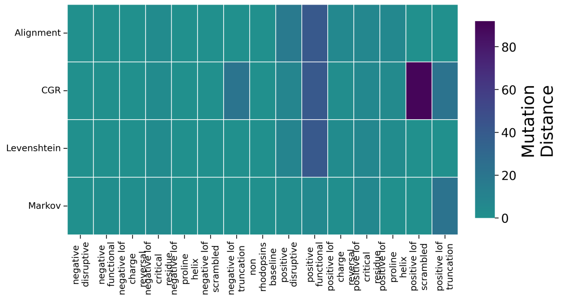

(b) Structure Modality

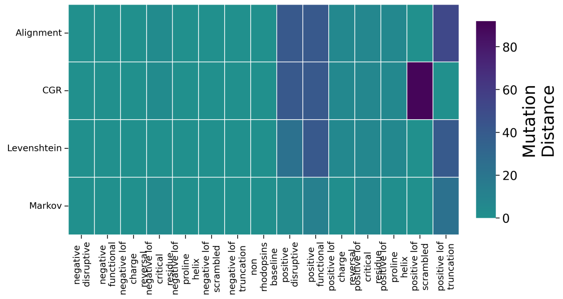

(c) Hybrid Modality

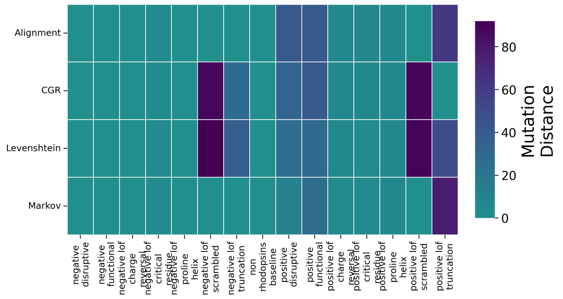

(d) Stacked Modality

Figure CS24: **Proxy acceptance persistence via IC50 (Mutation Distance at 50% Acceptance).** IC50 = mutation distance where acceptance drops to 50%. (a) Sequence modality IC50 heatmap. (b) Structure modality. (c) Hybrid modality. (d) Stacked modality: separation between conservative and radical-substitution proxy curves enables exploration while maintaining sequence-plausibility boundaries.

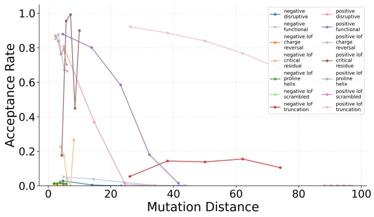

(a) Markov Stacked

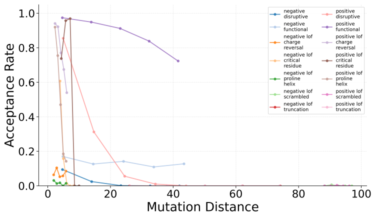

(b) Levenshtein Structure

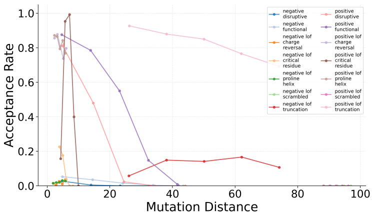

(c) Markov Sequence

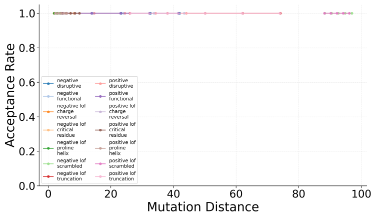

(d) Levenshtein Stacked

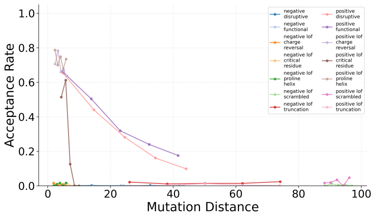

(e) CGR Stacked

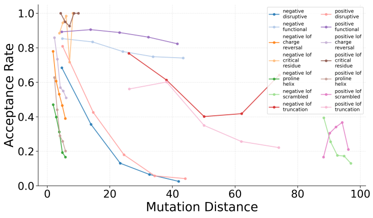

(f) Markov Structure

Figure CS25: **Acceptance Rate Trajectories across Mutation Distance.** Conservative (solid) vs radical-substitution (dashed) synthetic variant acceptance curves. (a) Markov Stacked: clear conservative-radical separation across mutation distances. (b) Levenshtein Structure: selective acceptance of conservative variants at low distance. (c) Markov Sequence: higher overall acceptance across both classes. (d) Levenshtein Stacked: inverted selectivity (lower conservative than radical acceptance). (e) CGR Stacked: limited class separation. (f) Markov Structure: reduced class separation relative to stacked models.

#### 2.4 Full GA Ablation and Sensitivity Analysis

The main text summarises the key prior and filter-strength effects. The full component-wise ANOVA, convergence distributions, success-rate breakdown, and trajectory analyses are provided below (Supplementary Figs. CS27, CS28, CS29, CS30, CS31, CS32, CS33, CS34, and CS35).

We quantified the contribution of each pipeline component to convergence speed, predicted wavelength, and sequence plausibility (Supplementary Figs. CS27–CS35). In these ablations, success means reaching the scalar wavelength objective within tolerance while satisfying the Markov plausibility constraint.

One-way ANOVA over 200 uniformly sampled configurations identified population size as the dominant driver of convergence time ($\eta^2 = 0.587$, $p < 0.0001$), followed by parent count ($\eta^2 = 0.271$, $p = 0.0017$). Mutation factor dominated final predicted wavelength ($\eta^2 = 0.98$, $p < 0.0001$) but had minimal effect on convergence time ($\eta^2 < 0.01$).

We then varied Markov filter strength (0.05–0.40), mutation prior (uniform, entropy, MI, inverse-MI, inverse-entropy, regressor, inverse-regressor), and target wavelength (490, 450, and 410 nm) with twelve independent seeds per condition, giving 84 runs per prior–target cell and 84 runs per filter–target cell (Supplementary Table CS16). At permissive filter strengths (0.05–0.10), success rates exceeded 94% across all targets (490 nm: 100%; 450 nm: 100%; 410 nm: 94–96%). Stricter thresholds progressively reduced success, with the hardest target most affected: at filter strength 0.30, success fell to 16.7% for 410 nm while remaining 48.8% for 490 nm; at 0.40, only 6.0% (5/84) of 410 nm runs succeeded versus 50.0% (42/84) for 490 nm.

Without filtering, 249/252 (98.8%) of no-filter runs converged to the wavelength target, but post hoc Markov rescoring revealed that the vast majority of these designs fall below the 20th-percentile plausibility threshold (Supplementary Table CS16). The Markov model therefore acts during optimization, not only after it: it constrains the selectable sequence set, and the wavelength objective alone does not keep designs proton-pump-like.

All priors are blended with a 10% uniform floor over positions to guarantee non-zero mutation probability everywhere (Methods); without this, sparse priors such as Lasso-Regressor lock most of the alignment at probability 0 and the GA cannot reach any target. With the floor in place, success rates separate by how well each prior concentrates mutations on positions that move $\lambda_{\text{max}}$. All priors achieved 100% success at 490 and 450 nm; only Inverse-MI fell below 100% at 410 nm (75%, 9/12 runs per prior–seed group at filter 0.05), confirming it as the weakest prior for difficult blue-shifted targets.

At 490 nm with filter strength 0.05, MI-weighted and Inverse-Regressor converged at a median of 7 generations, Uniform and Inverse-Entropy at 8, Inverse-MI at 15, Entropy at 24, and Regressor at 26. At 410 nm the ordering held but absolute generations grew (Inverse-Entropy 18, Inverse-Regressor 18, MI 21, Uniform 22, Entropy 60, Regressor 68; Inverse-MI converged in only 9/12 runs, median 41 among those). Stricter filtering kept trajectories closer to the training distribution (Figs. CS29–CS31; Fig. CS26).

We conducted ablation studies to determine how the GA's performance depends on key parameters and architectural components (filter and prior).

##### Ablation of Evolutionary Hyperparameters

We performed a Monte Carlo hyperparameter sweep over evolutionary parameters across 200 sampled configurations, with one random seed per sampled configuration. We measured performance using convergence generation and final predicted wavelength. Statistical analysis using one-way ANOVA identified significant main effects of population size ($p < 0.0001$, $\eta^2 = 0.587$) and parents count ($p = 0.0017$, $\eta^2 = 0.271$) on convergence generation. Mutation factor was significant for final predicted value ($p < 0.0001$, $\eta^2 = 0.980$ in the unbinned sampled values; $\eta^2 = 0.179$ after class binning), but was not significant for convergence generation in the binned analysis.

The sampled hyperparameters affected convergence unevenly across their parameter classes (Fig. CS27). Population size (Fig. CS27a) showed the strongest convergence-time effect, consistent with larger populations maintaining more search diversity. Parents count also affected convergence, whereas crossover parameters showed weaker marginal effects. Mutation-factor classes did not show a significant convergence-generation effect in the binned ANOVA, but mutation factor strongly affected final predicted wavelength. Two-way heatmaps (Fig. CS28) should therefore be interpreted as exploratory interaction screens over the sampled random-search space rather than fully replicated factorial estimates.

##### Ablation of Filter Strength and Prior

To quantify sensitivity to filter stringency and hotspot prior choice, we systematically varied (i) the Markov filter strength (values: 0.05, 0.1, 0.15, 0.2, 0.25, 0.3, 0.35, 0.4), (ii) the mutation prior type (uniform, entropy-weighted, MI-weighted, inverse-MI, inverse-entropy, inverse-regressor, regressor), and measured performance across target wavelengths (490, 450, and 410 nm). The GA was re-run with three independent seeds (42, 123, 456) per filter-prior-target condition, yielding 504 runs. Performance was quantified by convergence generation (first generation where predicted wavelength is within 1% of target value), final predicted wavelength, and success rate (convergence achieved by the end of the run).

**Filter Strength Effect:** At the minimal filter setting (strength 0.05) with uniform mutation prior, median convergence was 8 generations at target 490 nm, with 100% success. In this context, *filter strength* is the quantile used to set the Markov threshold from positive proton-pump training scores; strength 0.05 sets a permissive boundary near the lower tail of known positive examples. As filter strength increased from 0.05 to 0.25, target-dependent behavior emerged: at 490 nm, success remained high (90.5–100%); at 450 nm, success declined from 100% to 66.7–85.7%; at 410 nm, success dropped to 23.8–28.6% once strength reached 0.15. At strengths $\geq 0.3$, success collapsed to 0% for 410 and 450 nm, while 490 nm retained 23.8% success at strength 0.4. For the most challenging 410 nm target, strengths 0.05–0.1 maintained 90.5% success across all priors, and the uniform-prior median convergence was 16–17 generations. These results indicate that blue targets require looser filtering (strength $\leq 0.1$), whereas red targets tolerate stronger filtering (up to 0.25).

**No-Filter Post Hoc Check:** We also ran a true no-filter baseline by disabling Markov scoring during evolution. This condition tests whether the wavelength objective alone keeps the search inside the proton-pump-like sequence distribution. It did not. Across 63 no-filter runs (seven priors, three targets, three seeds), all runs could optimize the wavelength objective, but post hoc rescoring of the final candidates with the standard Markov proton-pump sequence-plausibility model showed that only 10/63 candidates (15.9%) passed a moderate 0.20 quantile threshold, and only 7/63 (11.1%) passed thresholds of 0.30–0.50. The failure was target-dependent: at the 0.20 threshold, 9/21 candidates passed for the 490 nm target, 1/21 for 450 nm, and 0/21 for 410 nm. Thus, without the filter, the GA often satisfies the spectral objective by moving into sequence regions that would fail the intended proton-pump plausibility constraint.

Representative LDA projections illustrate how increasing filter strength constrains the search trajectory for each target wavelength (Fig. CS29–CS31).

**Prior Type Effect:** At minimal filter setting (strength 0.05) and target 490 nm, all prior types achieved 100% success rate but with substantial variance in convergence speed. MI-weighted prior showed the fastest convergence with median 6 generations ($n = 3$ seeds). Entropy, inverse-entropy, inverse-regressor, regressor, and uniform priors showed median 8 generations, a 2-generation slowdown versus MI. Inverse-MI prior performed worst among all priors: median convergence 18 generations (~3-fold slower than MI), yet still achieved 100% success. This pattern indicates that priors derived from information-theoretic features can align with the adaptive landscape at this target, while inverse-priors are counterproductive but not catastrophic. At target 410 nm (blue, extrapolated), convergence was slower and more variable: at strength 0.05, MI converged in a median of 19 generations, entropy in 16 generations, and inverse-MI in 35 generations with only one successful seed.

**Filter × Prior Interaction:** Increasing filter stringency had severe consequences at target 410 nm: under a uniform prior, success dropped from 100% at strength 0.05 to 33.3% at strength 0.15, and median convergence among successful runs slowed from 16 to 38 generations (Fig. CS32). The relative effect of prior type variation became secondary compared to the filter phase transition. At target 490 nm, the same filter-strength sweep showed gentler degradation: under a uniform prior, convergence at strength 0.05 was 8 generations, increasing to 12 generations at strength 0.2, with success remaining 100% up to strength 0.25. Thus, the filter's impact is target-dependent: stronger at blue targets than at red targets. Changing from MI to uniform prior at fixed filter 0.05 increased convergence from 6 to 8 generations at target 490 nm, whereas at target 410 nm the uniform and entropy-like priors converged faster than MI in this small three-seed sample.

**Target Wavelength Sensitivity:** At filter strength 0.05 with MI prior, convergence times at different targets showed clear spectral dependence: target 410 nm (blue, extrapolated) required median 19 generations, target 450 nm required 11 generations, and target 490 nm required 6 generations. Across all priors at filter strength 0.05, the corresponding medians among successful runs were 16, 10, and 8 generations, respectively. The slower convergence for 410 nm reflects the extrapolation challenge identified in the Phylogenetic Analysis section. With the exception of inverse-MI at 410 nm, all strength-0.05 prior-target combinations achieved 100% success across the three seeds.

**Summary of Ablation Patterns:** The ablation reveals four key findings: (1) Filter strength exhibits a steep phase transition: restrictive filters ($\geq 0.3$) collapse to 0% success at 410 and 450 nm, while 490 nm retains 23.8% success at strength 0.4. (2) Prior type modulation produces 2-3 generation differences between MI-derived and inverse priors, with 100% success maintained across all prior types at minimal filter strength. (3) Target wavelength and filter strength interact strongly: blue targets are slower and require looser filtering; red targets tolerate tighter filtering and converge faster. (4) A no-filter search can optimize predicted wavelength, but post hoc Markov rescoring shows that many final sequences, especially at blue targets, fail the proton-pump sequence-plausibility constraint. Quantification of success rates (Table CS7 and Fig. CS35) confirms these patterns.

At the permissive 0.05 filter setting, success remained high across targets, but convergence speed depended strongly on target and prior. For target 490 nm, MI reached the target in a median of 6 generations compared with 18 generations for inverse-MI. For target 410 nm, successful runs were slower and inverse-MI was least reliable (33.3% success at strength 0.05). The no-filter baseline converged rapidly in predicted wavelength space (mean convergence generation 3.24 across all 63 runs), but post hoc Markov rescoring showed that most of these final candidates would not pass the normal proton-pump plausibility criterion at moderate thresholds. These results indicate that (i) filter strength must be tuned relative to target spectral location, (ii) prior choice impacts convergence speed, (iii) blue extrapolation targets require more permissive filtering, and (iv) the filter enforces a real sequence-plausibility constraint that the optimizer would otherwise violate. We compared these settings through computational re-runs of the GA; experimental assays were performed only on final designs from the intact pipeline.

##### Summary of Ablation Results

Together, these hyperparameter and grid sweeps show that GA behavior depends on algorithmic settings, statistical filtering, and guided mutation. The ML regressor provides a fast objective but shows spectral-tail extrapolation bias; the Markov filter provides sequence-plausibility screening but can reject too many candidates at strict thresholds; and the hotspot prior changes convergence speed by biasing mutation placement. The no-filter baseline sharpens this interpretation: removing the filter allowed rapid wavelength optimization, but most final sequences failed post hoc Markov proton-pump plausibility scoring. The hyperparameter sensitivity analysis (200 sampled configurations) shows that population size and parent count affect convergence in the sampled range, while mutation factor strongly affects final predicted wavelength. The schema grid shows that permissive filtering and suitable priors are especially important for blue extrapolation targets. These findings support using the pipeline as a tunable in-silico design workflow, with experimental calibration required before transferring the same settings to other objectives.

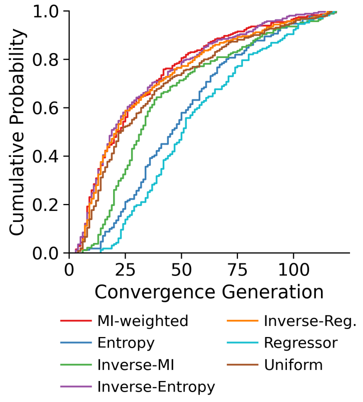

(a) Convergence by prior

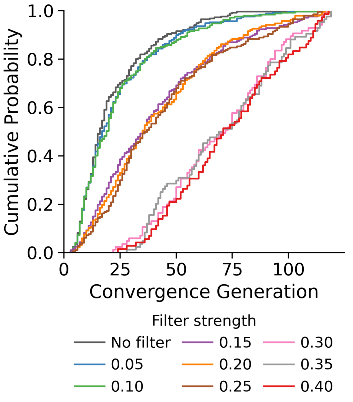

(b) Convergence by filter

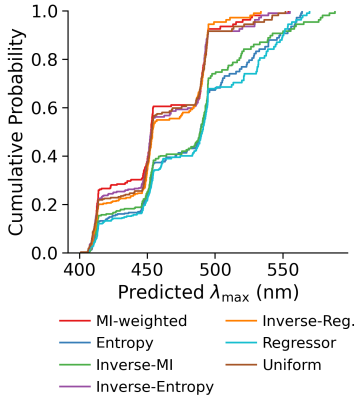

(c) Predicted $\lambda_{\max}$ by prior

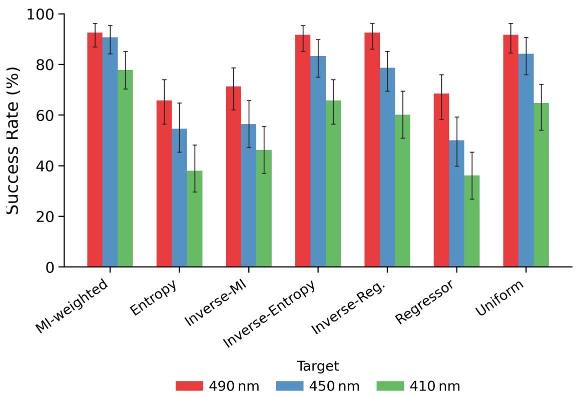

(d) Success rate by prior and target wavelength

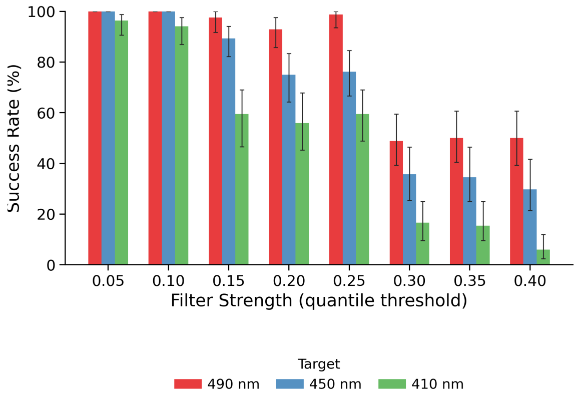

(e) Success rate by filter strength and target wavelength

Figure CS26: **GA ablation: effect of mutation prior and filter strength on convergence and success.** (a) ECDF of convergence generation by mutation prior (all targets pooled); Inverse-MI converges slowest and MI-weighted and Uniform are fastest. (b) ECDF of convergence generation by filter strength; stricter thresholds slow or prevent convergence; the unfiltered condition (gray) converges fastest but without sequence plausibility constraints. (c) ECDF of final predicted $\lambda_{\max}$ by prior, showing Inverse-MI produces bluer-shifted sequences while MI-weighted concentrates near the target. (d) Success rate (convergence before generation limit) by prior and target wavelength (490, 450, 410 nm); MI-weighted is highest at every target; Inverse-MI underperforms; the remaining priors track each other within ∼10 percentage points. (e) Success rate by filter strength; permissive thresholds (0.05–0.10) achieve near-100% success across all targets; strict thresholds (≥0.30) prevent convergence at 450 and 410 nm. Error bars are 95% bootstrap CIs over grouped runs (panel d: 84 runs per prior–target group; panel e: 84 runs per filter–target group). Full hyperparameter interactions are in Supplementary Figs. CS27–CS35.

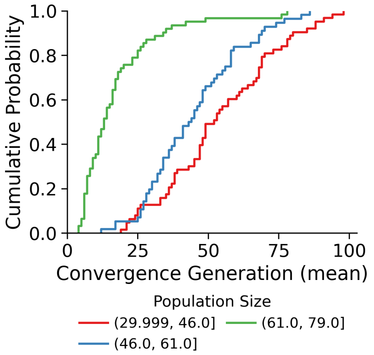

(a) Population Size

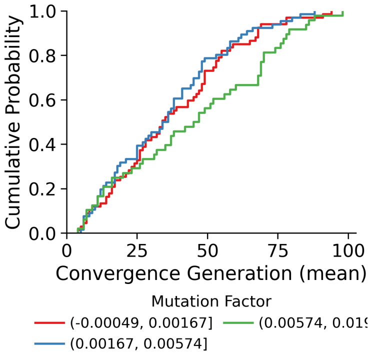

(b) Mutation Factor

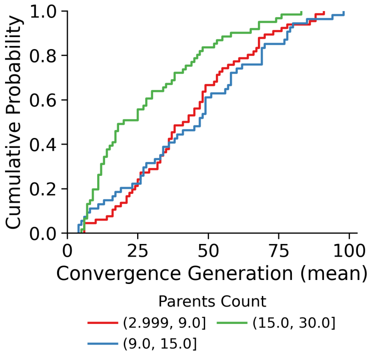

(c) Parents Count

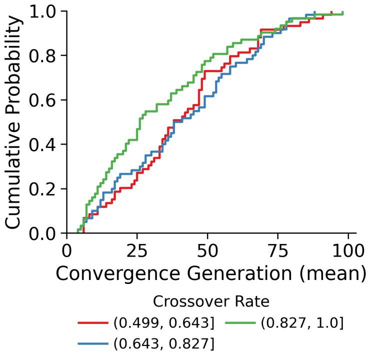

(d) Crossover Rate

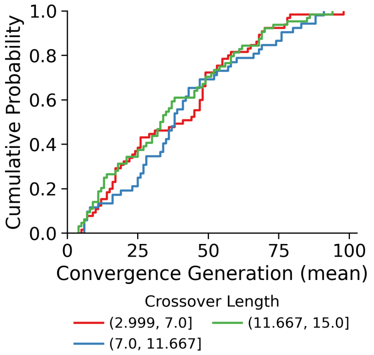

(e) Crossover Length

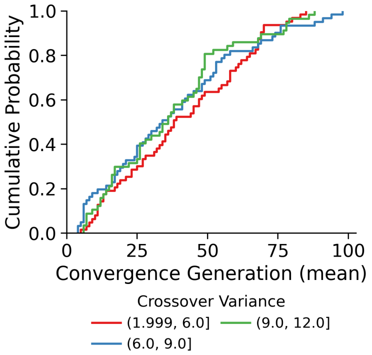

(f) Crossover Variance

Figure CS27: **Main effects of evolutionary hyperparameters on convergence time.** (a-f) Class-stratified ECDFs show the marginal effect of each hyperparameter on convergence generation. Population size (a) and parents count (c) show the clearest convergence effects in the sampled sweep, mutation factor (b) affects final predicted wavelength more strongly than convergence time, and crossover parameters (d-f) show weaker marginal effects.

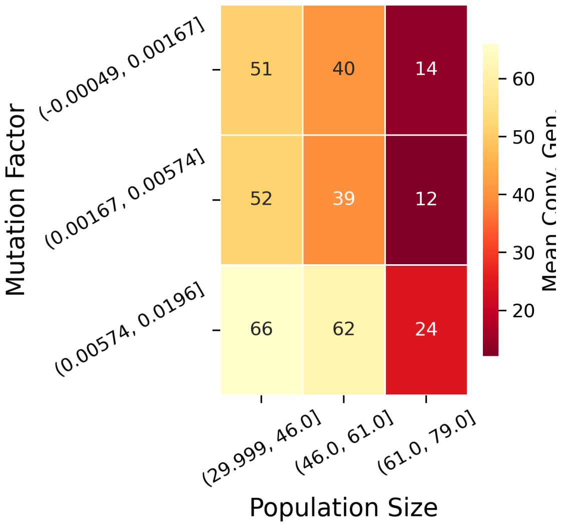

(a) Mut Factor × Pop Size

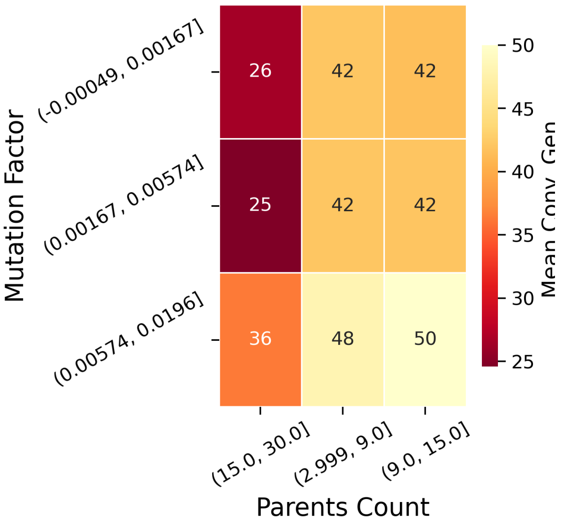

(b) Mut Factor × Parents

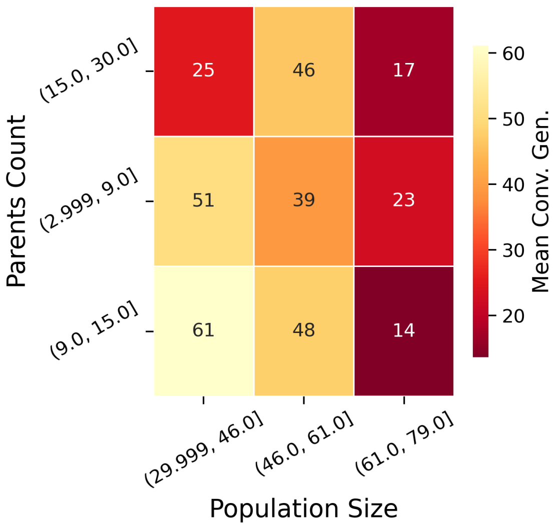

(c) Parents × Pop Size

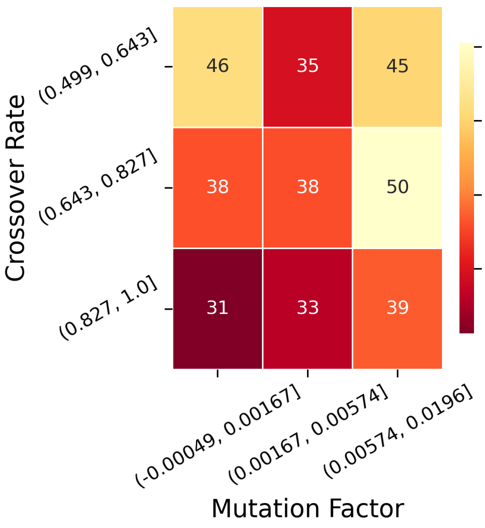

(d) Crossover × Mut Factor

Figure CS28: **Exploratory interactions between sampled hyperparameters.** Heatmaps summarize mean convergence generation across pairs of class-binned hyperparameters in the random-search sweep. (a) Population size and mutation factor. (b) Mutation factor and parents count. (c) Population size and parents count. (d) Crossover rate and mutation factor. These plots are exploratory because exact hyperparameter combinations were not fully factorially replicated. Lower convergence generation indicates faster in-silico target attainment.

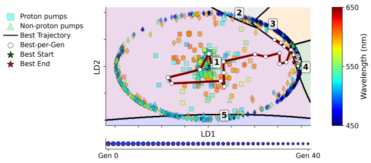

(a) 0.05, seed 42

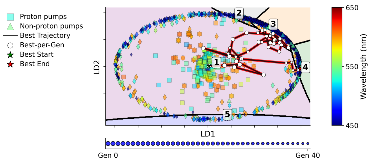

(b) 0.05, seed 123

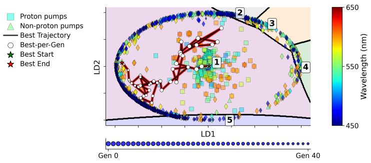

(c) 0.05, seed 456

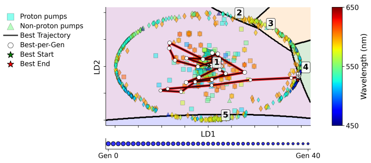

(d) 0.2, seed 42

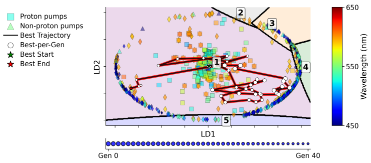

(e) 0.2, seed 123

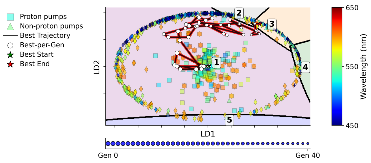

(f) 0.2, seed 456

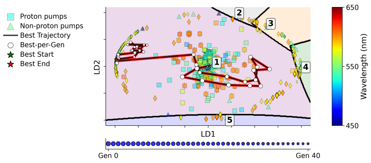

(g) 0.35, seed 42

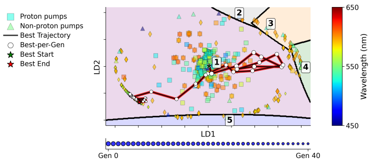

(h) 0.35, seed 123

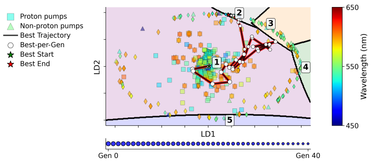

(i) 0.35, seed 456

Figure CS29: **Representative LDA trajectory projections across filter strengths for target 410 nm.** Uniform mutation prior runs are shown for three filter strengths (0.05, 0.2, 0.35; rows) and three seeds (42, 123, 456; columns).

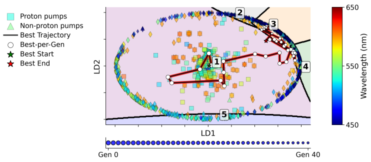

(a) 0.05, seed 42

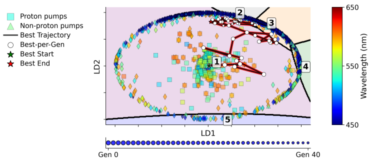

(b) 0.05, seed 123

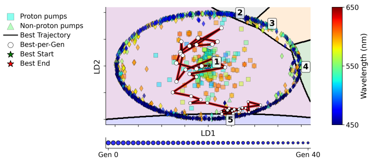

(c) 0.05, seed 456

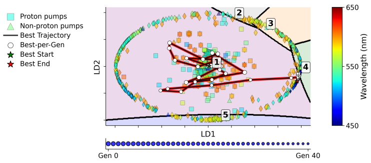

(d) 0.2, seed 42

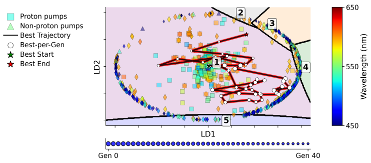

(e) 0.2, seed 123

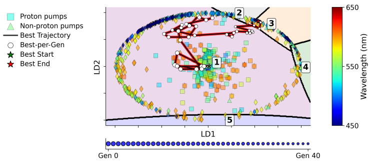

(f) 0.2, seed 456

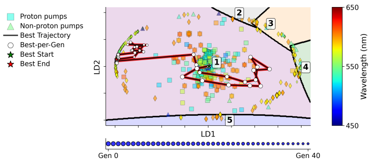

(g) 0.35, seed 42

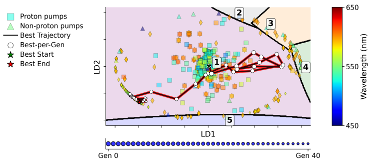

(h) 0.35, seed 123

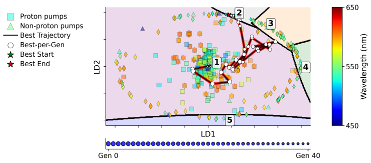

(i) 0.35, seed 456

Figure CS30: **Representative LDA trajectory projections across filter strengths for target 450 nm.** Uniform mutation prior runs are shown for three filter strengths (0.05, 0.2, 0.35; rows) and three seeds (42, 123, 456; columns).

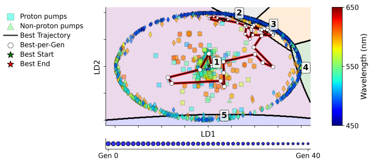

(a) 0.05, seed 42

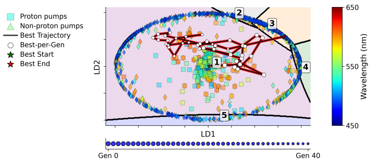

(b) 0.05, seed 123

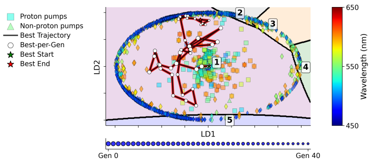

(c) 0.05, seed 456

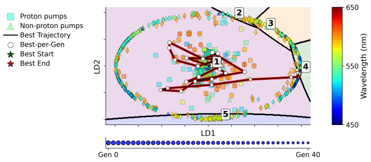

(d) 0.2, seed 42

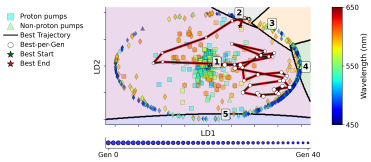

(e) 0.2, seed 123

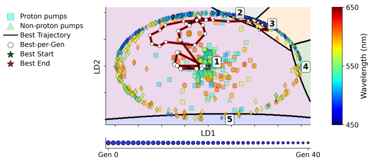

(f) 0.2, seed 456

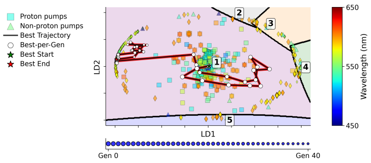

(g) 0.35, seed 42

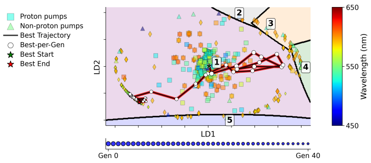

(h) 0.35, seed 123

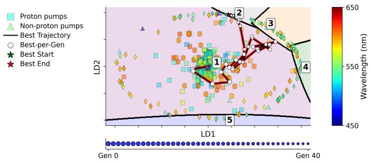

(i) 0.35, seed 456

Figure CS31: **Representative LDA trajectory projections across filter strengths for target 490 nm.** Uniform mutation prior runs are shown for three filter strengths (0.05, 0.2, 0.35; rows) and three seeds (42, 123, 456; columns).

Table CS7: **Evolutionary Success Rates.** Percentage of independent runs achieving the target phenotype (within 1% tolerance), aggregated across seven mutation priors and three seeds per filter-target pair ($n = 21$ runs). A sharp phase transition is observed: restrictive filters ($\geq 0.3$) cause total search collapse, particularly for blue-shifted targets (410 nm).

| **Filter Strength** | **410** nm | **450** nm | **490** nm |
|---|---|---|---|
| 0.05 | 90.5% | 100.0% | 100.0% |
| 0.1 | 90.5% | 100.0% | 100.0% |
| 0.15 | 23.8% | 66.7% | 100.0% |
| 0.2 | 28.6% | 81.0% | 90.5% |
| 0.25 | 28.6% | 85.7% | 95.2% |
| 0.3 | 0.0% | 0.0% | 0.0% |
| 0.35 | 0.0% | 0.0% | 0.0% |
| 0.4 | 0.0% | 0.0% | 23.8% |

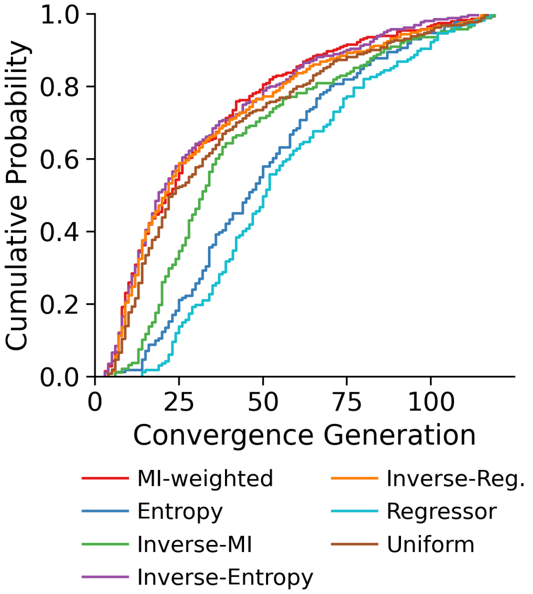

(a) Conv. Time (Prior)

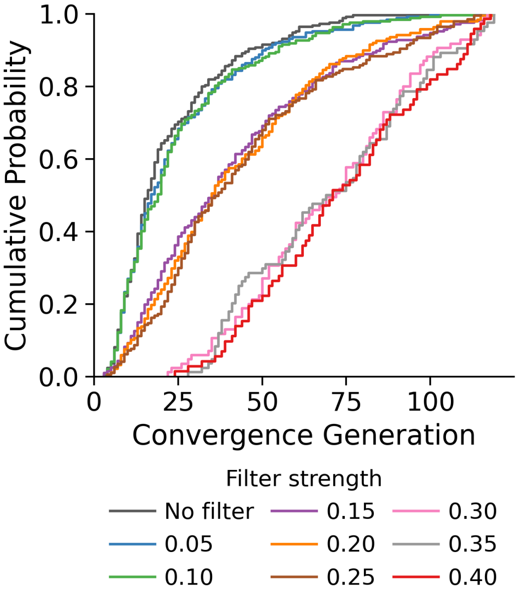

(b) Conv. Time (Filter)

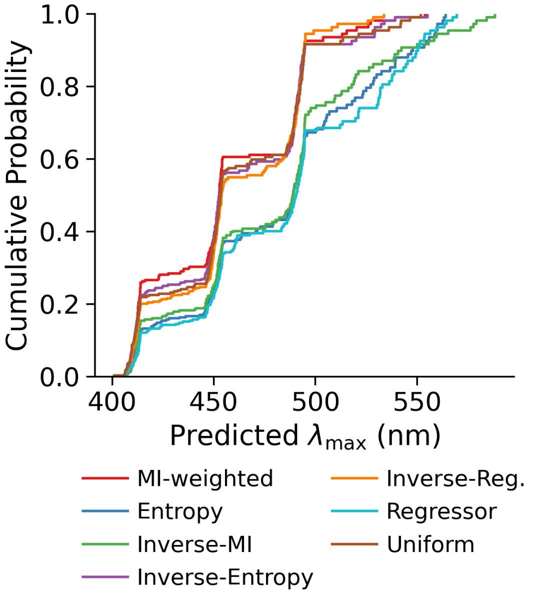

(c) Pred. Wave. (Prior)

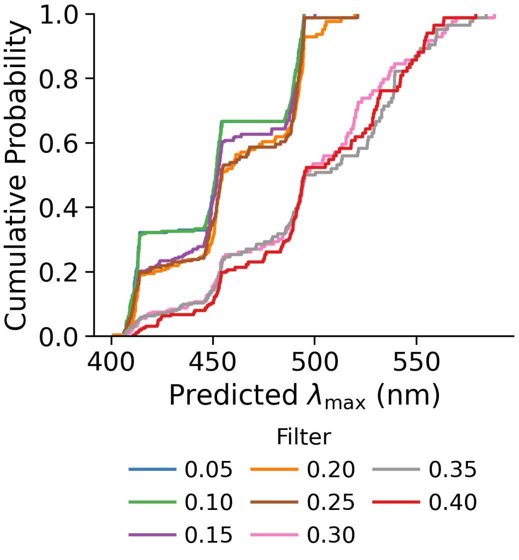

(d) Pred. Wave. (Filter)

Figure CS32: **Main effects of filter strength and positional prior on convergence and wavelength.** (a-b) ECDF plots show convergence generation across mutation-prior classes and Markov filter-strength settings. Information-theoretic priors alter convergence speed, while stricter filter thresholds reduce feasible search space. (c-d) Final predicted wavelength distributions show how prior type and filter strength shift the computational search outcome.

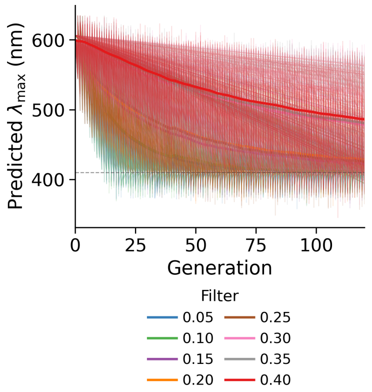

(a) Target 410nm - Filter

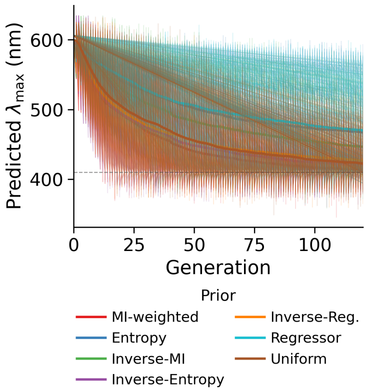

(b) Target 410nm - Prior

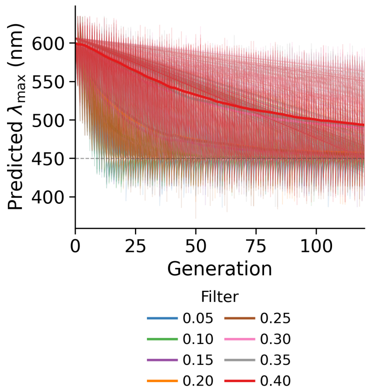

(c) Target 450nm - Filter

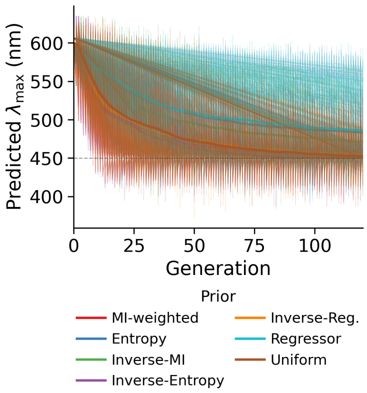

(d) Target 450nm - Prior

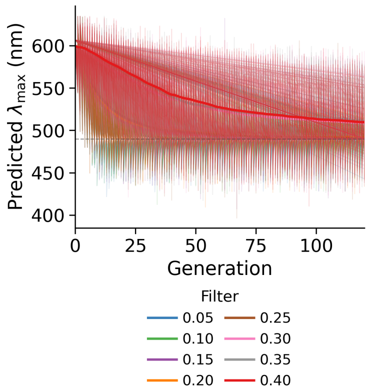

(e) Target 490nm - Filter

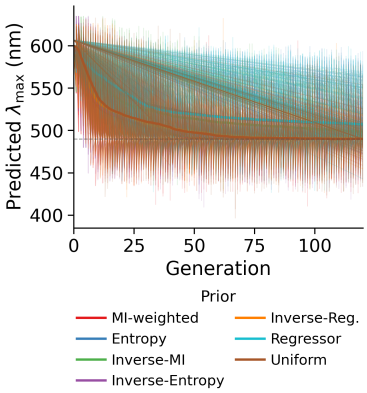

(f) Target 490nm - Prior

Figure CS33: **Evolutionary trajectories across generations stratified by target wavelength and ablation condition.** Each panel shows mean trajectories (bold) and individual runs (faded) for predicted wavelength across generations. (Left Column) Filter ablation reveals its role in constraining the search, especially for blue-shifted targets. (Right Column) Positional prior comparison shows the acceleration provided by information-theoretic weighting (MI, entropy) versus uniform search.

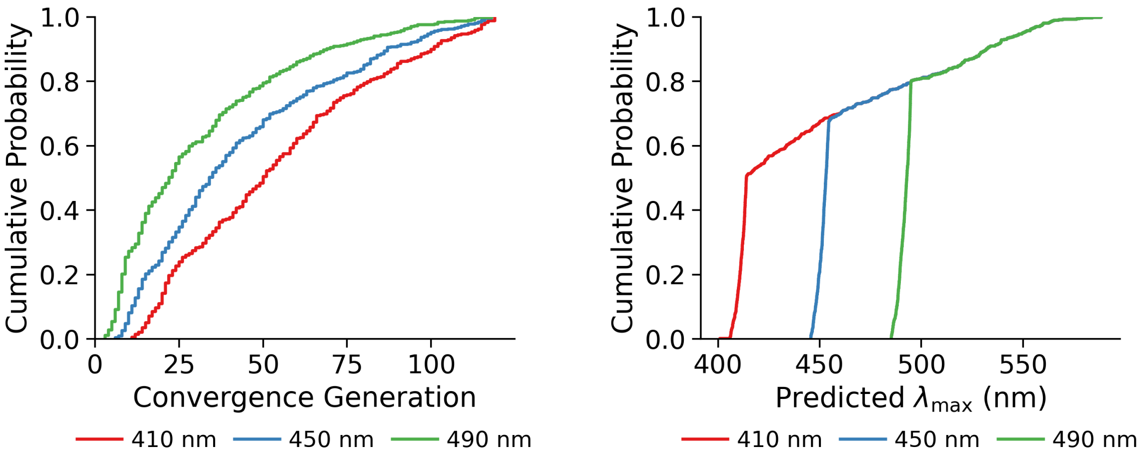

(a) Conv. Time (Target)

(b) Pred. Wave. (Target)

Figure CS34: **Target wavelength dependency of convergence dynamics.** (a) Convergence generation stratified by target wavelength (410, 450, 490 nm) shows the extrapolation challenge: at filter strength 0.05 with MI prior, blue targets (410 nm) require median 19 generations, while 450 nm and 490 nm targets converge in 11 and 6 generations respectively. (b) Final wavelength prediction accuracy across all targets shows that successful runs reach the target wavelength despite spectral distance from training data.

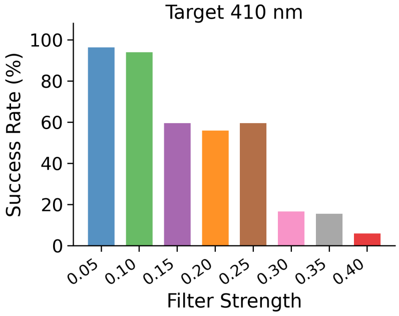

(a) Target 410nm - Filter

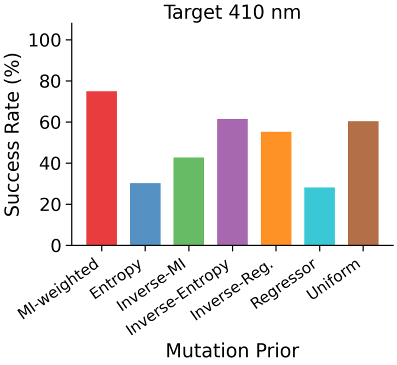

(b) Target 410nm - Prior

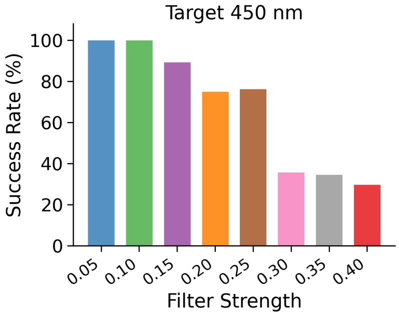

(c) Target 450nm - Filter

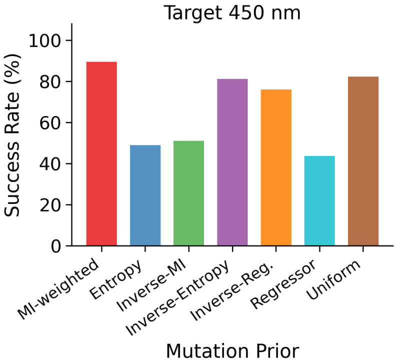

(d) Target 450nm - Prior

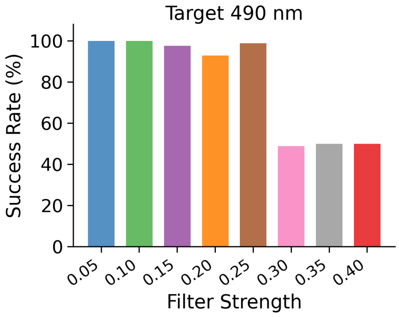

(e) Target 490nm - Filter

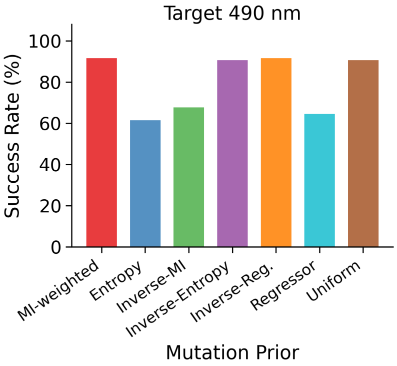

(f) Target 490nm - Prior

Figure CS35: **Success rate analysis across target wavelengths, filter strengths, and priors.** Success rates (final predicted wavelength within $\pm 1\%$ of target) for different targets (rows) and grid settings (columns). (Left Column) Success rate versus filter strength shows a phase transition across all targets, where restrictive filters ($\geq 0.3$) cause search failure. (Right Column) Mutation prior comparison shows that information-theoretic priors (MI, entropy) maintain high success rates, while inverse-priors reduce performance but remain stable at permissive filter settings.

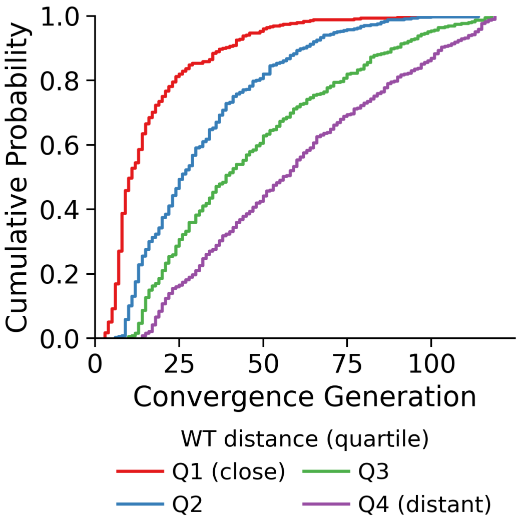

(a) Convergence generation by distance from wildtype

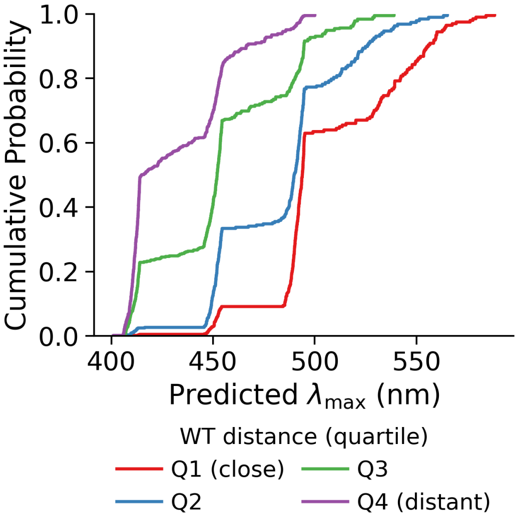

(b) Predicted $\lambda_{max}$ by distance from wildtype

Figure CS36: **GA outcomes stratified by Levenshtein distance from the wildtype reference.** (a) ECDF of convergence generation binned by the final design's Levenshtein distance from the nearest wildtype proton pump; designs requiring more mutations to reach the target converge later. (b) Final predicted $\lambda_{max}$ stratified by wildtype distance; blue-shifted targets necessarily accumulate more mutations.

### 3 Computational Supplementary Tables

Table CS8: **Supplementary Table CS1: Quantitative breakdown of the rhodopsin sequence database.** Summary of functional classes, sequence counts ($N = 884$), and absorption wavelength ($\lambda_{max}$) statistics. Proton pumps dominate the dataset ($n = 310$, 35.1%) but are excluded from the blue extreme (< 500 nm), which is occupied by channelrhodopsins ($n = 37$, median 480 nm).

| **Functional Class** | $n$ | **%** | **Med. $\lambda_{max}$ (nm)** | **Mean $\lambda_{max}$ (nm)** | **Range (nm)** |
|---|---|---|---|---|---|
| Proton Pump | 310 | 35.1 | 548.0 | 551.4 | 455–622 |
| Proton Sensory | 237 | 26.8 | 535.0 | 531.8 | 460–618 |
| Sodium Pump | 220 | 24.9 | 527.0 | 530.3 | 465–581 |
| Chloride Pump | 59 | 6.7 | 548.0 | 547.5 | 505–583 |
| Channel | 37 | 4.2 | 480.0 | 493.1 | 453–587 |
| Unknown | 21 | 2.4 | 534.0 | 525.1 | 436–568 |
| **Total / Combined** | **884** | **100.0** | **537.0** | **539.9** | **436–622** |

Table CS9: **Supplementary Table CS2: Pairwise Sequence, Structural Identity, and Close-Homolog Statistics.** Comparison of Levenshtein identity metrics in primary sequence versus secondary structure space for unique proton pump sequences ($n = 309$ for primary sequence; $n = 138$ with secondary-structure annotations). Primary sequence shows high divergence (mean identity 29.43%, median 8.84%) while secondary structure is highly conserved (mean 74.03%, median 70.77%). BLAST-like $E$-value summaries show that 95.8% of primary sequences have at least one close proton-pump homolog ($E < 0.01$) in the dataset.

| **Metric** | **Primary Sequence** | **Sec. Structure** | **Ratio (S/P)** |
|---|---|---|---|
| Mean Identity | 29.43% | 74.03% | 2.52 |
| Median Identity | 8.84% | 70.77% | 8.01 |
| Standard Deviation | 37.89% | 13.43% | 0.35 |
| Min / Max Identity | 2.87% / 99.63% | 47.70% / 99.62% | 16.62 / 1.00 |
| Pairwise $E$-Value < 0.01 | 23.9% | 100.0% | – |
| Sequences with min $E$-Value < 0.01 | 95.8% | 100.0% | – |

Table CS10: **Supplementary Table CS3: Regressor Performance Comparison.** Ten-fold cross-validation on the full dataset ($N = 884$) compares LassoXGBoost stack against RandomForest, XGBoost, and LASSO baselines. Physicochemical embedding (Feature Map) achieves lowest RMSE (12.64 nm) and highest $R^2_{CV}$ (0.840).

| **Modality** | **Architecture** | $R^2_{test}$ | $R^2_{CV}$ | $RMSE_{CV}$ | $R^2_{MC}$ | $RMSE_{MC}$ |
|---|---|---|---|---|---|---|
| **Feature Map** | **LassoXGBoost** | **0.843** | **0.840** | **12.64** | **0.835** | **12.42** |
| | RandomForest | 0.771 | 0.782 | 14.41 | 0.785 | 14.22 |
| | XGBoost | 0.825 | 0.800 | 13.83 | 0.812 | 13.55 |
| | LASSO | 0.783 | 0.783 | 14.39 | 0.780 | 14.51 |
| **One-Hot** | LassoXGBoost | 0.805 | 0.812 | 13.38 | 0.807 | 13.35 |
| | ExtraTrees | 0.749 | 0.799 | 13.86 | 0.799 | 14.27 |
| | SVR (Radial) | 0.353 | 0.353 | 24.86 | 0.359 | 24.99 |
| **Ordinal** | LassoXGBoost | 0.773 | 0.750 | 15.46 | 0.741 | 15.66 |

Table CS11: **Supplementary Table CS4: Family-Specific Predictive Reliability.** Performance of LassoXGBoost (Feature Map) stratified by rhodopsin subfamily. Well-represented classes (e.g., Bacteriorhodopsin, $n = 262$) show higher $R^2$ than sparse classes (e.g., Xenorhodopsin, $n = 14$).

| **Subfamily** | $n$ | $R^2$ | **RMSE (nm)** | **MAE (nm)** |
|---|---|---|---|---|
| Bacteriorhodopsin | 262 | 0.879 | 12.46 | 8.84 |
| All Proton Pumps | 310 | 0.843 | 13.11 | 9.73 |
| Sensory Rhodopsin | 107 | 0.784 | 11.21 | 7.72 |
| Archaeorhodopsin | 16 | 0.732 | 15.67 | 10.28 |
| Proteorhodopsin | 139 | 0.717 | 11.13 | 8.59 |
| Halorhodopsin | 36 | 0.577 | 10.91 | 7.39 |
| Channelrhodopsin | 40 | 0.476 | 22.01 | 14.13 |
| Xenorhodopsin | 14 | -0.302 | 15.94 | 11.47 |

Subfamily classification differs from the functional superclass taxonomy in Table CS8. 'Channelrhodopsin' ($n = 40$) encompasses all channel-type sequences; 'Sensory Rhodopsin' ($n = 107$) is a sub-lineage of the broader 'Proton Sensory' superclass ($n = 237$).

Table CS12: **Supplementary Table CS5: Performance of Coarse-Grained Rhodopsin Filters.** Metrics are median ± fold SD from random sequence-level 10-fold cross-validation for discriminating rhodopsins ($n = 884$) from non-rhodopsin ion transporters ($n = 10,000$).

| **Architecture** | **Modality** | **AUROC** | **PR-AUC** | **F1-Score** |
|---|---|---|---|---|
| **Levenshtein** | **Sequence** | **1.000 ± 0.001** | **1.000 ± 0.001** | **0.989 ± 0.011** |
| | Structure | 1.000 ± 0.002 | 1.000 ± 0.002 | 0.989 ± 0.019 |
| **Markov Model** | **Stacked** | **0.999 ± 0.007** | **0.999 ± 0.007** | **0.980 ± 0.022** |
| | Structure | 0.980 ± 0.024 | 0.966 ± 0.055 | 0.960 ± 0.035 |
| Alignment | Hybrid | 1.000 ± 0.007 | 1.000 ± 0.005 | 0.989 ± 0.032 |
| | Sequence | 0.917 ± 0.094 | 0.960 ± 0.045 | 0.679 ± 0.094 |
| CGR | Hybrid | 0.896 ± 0.122 | 0.924 ± 0.095 | 0.799 ± 0.151 |

Table CS13: **Supplementary Table CS6: Subfamily Specificity Breakdown.** Discrimination of Proton Pumps ($n = 310$) from phylogenetically related families (chloride pumps and sensory rhodopsins; $n = 296$) under random sequence-level cross-validation. Metrics are median ± fold SD.

| **Architecture** | **Modality** | **AUROC*** | **PR-AUC*** | **F1-Score*** |
|---|---|---|---|---|
| **Markov Model** | **Stacked** | **1.000 ± 0.157** | **1.000 ± 0.119** | **0.984 ± 0.165** |
| Markov Model | Structure | 0.489 ± 0.293 | 0.490 ± 0.241 | 0.618 ± 0.249 |
| Levenshtein | Structure | 1.000 ± 0.035 | 1.000 ± 0.024 | 0.944 ± 0.056 |
| Levenshtein | Sequence | 0.986 ± 0.058 | 0.991 ± 0.052 | 0.967 ± 0.091 |

*Large fold standard deviations for the Stacked Markov model reflect high between-fold variance in this small-sample subfamily setting, not values outside the metric range.

Table CS14: **Supplementary Table CS7: Evolutionary Convergence and Filter Dynamics.** Summary of the GA trajectory for blue-shifting a proton pump (Run 789). Convergence at generation 5 shows the population discovering a strongly blue-shifted candidate that passes the Markov sequence-plausibility threshold (filter $p \approx 0.27$).

| **Metric** | **Initial (G0)** | **Conv. (G5)** | **Final (G16)** | **Target** |
|---|---|---|---|---|
| Best $\hat{\lambda}_{\max}$ (nm) | 606.56 | 490.00 | 490.00 | 490.00 |
| Prediction Error (%) | 23.79% | 0.00% | 0.00% | 0.00% |
| Cumulative Mutations ($\Delta D_{Lev}$) | 0 | 26 | 26 | – |
| Markov Anomaly Score | -2.62 | -2.72 | -2.72 | $> -2.74$ |
| Filter Probability ($p$) | 0.32 | 0.27 | 0.27 | – |

Table CS15: **Supplementary Table CS8: Component-Wise Sensitivity Analysis.** ANOVA results ($F$-stat, $p$-value, $\eta^2$) for filter, target wavelength, and mutation-prior effects in the schema grid.

| **Component** | **Metric** | **F-stat** | **p-value** | $\eta^2$ | **Impact** |
|---|---|---|---|---|---|
| Sequence-Plausibility Filter | Predicted $\lambda_{\max}$ | 124.43 | $< 10^{-100}$ | 0.637 | Large |
| Sequence-Plausibility Filter | Convergence Gen. | 9.68 | $1.8 \times 10^{-8}$ | 0.164 | Large |
| Target Wavelength | Convergence Gen. | 33.45 | $1.3 \times 10^{-13}$ | 0.211 | Large |
| Mutation Prior | Convergence Gen. | 3.73 | 0.0014 | 0.083 | Medium |
| Mutation Prior | Predicted $\lambda_{\max}$ | 3.63 | 0.0016 | 0.042 | Small |

Table CS16: **Supplementary Table CS9: Evolutionary Success Rates.** Percentage of independent runs achieving the target phenotype (within 1% tolerance), aggregated across seven mutation priors and three seeds per filter-target pair ($n = 21$ runs).

| **Filter Strength** | **410** nm | **450** nm | **490** nm |
|---|---|---|---|
| 0.05 | 90.5% | 100.0% | 100.0% |
| 0.1 | 90.5% | 100.0% | 100.0% |
| 0.15 | 23.8% | 66.7% | 100.0% |
| 0.2 | 28.6% | 81.0% | 90.5% |
| 0.25 | 28.6% | 85.7% | 95.2% |
| 0.3 | 0.0% | 0.0% | 0.0% |
| 0.35 | 0.0% | 0.0% | 0.0% |
| 0.4 | 0.0% | 0.0% | 23.8% |

Table CS17: **Supplementary Table CS10: Proton-Pump Scaffold Composition.** Counts are wild-type scaffold annotations among proton-pump entries in the sequence database ($n = 310$). GPR and BR are the dominant mutagenesis series and together account for 223 of 310 proton-pump entries (71.9%).

| **Scaffold group** | **Count** ($n$) | **Proportion of proton pumps** |
|---|---|---|
| GPR | 126 | 40.6% |
| BR | 97 | 31.3% |
| GPR + BR | 223 | 71.9% |
| Other proton-pump scaffolds | 87 | 28.1% |
| **Total proton pumps** | **310** | **100.0%** |

[7] Trevor Hastie, Robert Tibshirani, and Jerome Friedman. *The Elements of Statistical Learning: Data Mining, Inference, and Prediction*. Springer, 2nd edition, 2009.

[14] Haris Saeed, Aidong Yang, and Wei Huang. Disentangling protein function via decoupled information theoretic selection of key tuning residues. *bioRxiv (Cold Spring Harbor Laboratory)*, 2025.

[15] Siméon Denis Poisson. *Recherches sur la probabilite des jugements en matiere criminelle et en matiere civile*. Bachelier, 1837.

[16] John Burkardt. The truncated normal distribution. Department of Scientific Computing, Florida State University, 2014.

[17] Johannes Ewald and Paul C. Ewald. Quantifying the acceptance probability of mutations in protein engineering. *PLoS Computational Biology*, 14(9):e1006469, 2018.
