## Supplementary material for "AI-enabled rhodopsin design for blue-light enhanced bacterial growth": Biological Supplementary Information

### Materials and methods

#### Molecular cloning for designed rhodopsins

Four machine-learning designed rhodopsins APR1, APR6, and APR7 were codon-optimised using codon usage frequencies for Cupriavidus necator (previously Ralstonia eutropha) obtained from the Kazusa Codon Usage Database (Kazusa DNA Research Institute, Japan; https://www.kazusa.or.jp/codon/), and were synthesised by Integrated DNA Technologies (IDT, UK) (Biological Supplementary Information Table S3). PCR amplification of gene inserts and vector backbones was performed using Q5 High-Fidelity DNA Polymerase (NEB, UK) with primers containing complementary overlap sequences (Biological Supplementary Information Table S6). PCR products were purified using the Monarch® PCR & DNA Cleanup Kit (NEB, UK) according to the manufacturer's protocol.

DNA fragments were assembled using NEBuilder® HiFi DNA Assembly Master Mix (NEB, UK) and transformed into E. coli DH5α High Efficiency Competent Cells (NEB, UK) via heat shock. Transformants were selected on LB agar plates containing tetracycline, and single colonies were cultured overnight. Plasmids were extracted using the QIAprep® Spin Miniprep Kit (Qiagen, UK) and verified by Nanopore sequencing.

#### Rhodopsin expression in *Escherichia coli* C41 (DE3)

Codon-optimised DNA fragments for *Escherichia coli* encoding APRs were chemically synthesised (Eurofins Genomics) and inserted into the NdeI-XhoI site of the pET21a(+) plasmid vector (Novagen). The plasmids were transformed into E. coli C41 (DE3) (Lucigen) cells, and colonies were inoculated into Luria-Bertani (LB) medium supplemented with 100 µg mL−1 ampicillin. Before protein expression, transformants were cultured at 37 °C in 2× YT medium (NaCl 5 g L−1, Bacto tryptone 16 g L−1, and Bacto yeast extract 10 g L−1, pH 7.0) with 100 µg mL−1 ampicillin until the OD660 reached 0.3. Protein expression was then induced at 37 °C for 4 h in the dark by adding 0.1 mM isopropyl β-D-1-thiogalactopyranoside (IPTG: Sigma-Aldrich) and 10 µM all-trans retinal (Sigma-Aldrich). Rhodopsin-expressing cells were collected by centrifugation at 4,400 × g for 3 min at 4 °C (MX-305, Tomy Seiko) and then washed twice in 100 mM NaCl.

#### Rhodopsin purification from *Escherichia coli* C41 (DE3)

For the APR protein purification, the plasmid was transformed into *E. coli* C43 (DE3) (Lucigen) cells. The protein expression was induced by 0.1 mM IPTG in the presence of 10 µM all-trans-retinal (Toronto Research Chemicals) for 4 h at 37 °C. The expressed proteins had a 6 × His-tag on the C-terminus. The expressed protein was purified using a 5 mL Co2+-NTA column (HiTrap TALON crude; Cytiva) on an ÄKTA go protein purification system (Cytiva). The rhodopsin-expressing cells were harvested and resuspended in a buffer containing 50 mM Tris-HCl (pH 8.0) and 5 mM MgCl2. The harvested cells were disrupted by sonication (Ultrasonic Homogenizer VP-300N, TAITEC). The membrane fraction was collected by ultracentrifugation (CP80NX, Eppendorf Himac Technologies) at 142,000 × g for 1 h. The proteins were solubilized in a buffer containing 50 mM Tris-HCl (pH 7.5), 300 mM NaCl, 10 µM all-trans-retinal and 3% DDM. Solubilized proteins were separated from insoluble fractions by ultracentrifugation at 142,000 × g for 1 h. After loading the solubilized proteins on the Co2+-NTA column, the column was washed with a buffer containing 50 mM Tris-HCl (pH 7.5), 300 mM NaCl, 10 mM imidazole, and 0.1% DDM. The His-tagged proteins were eluted using a 10–500 mM imidazole gradient with a buffer containing 50 mM Tris-HCl (pH 7.5), 300 mM NaCl, and 0.1% DDM. The eluted proteins were dialyzed using a buffer containing 20 mM

HEPES-NaOH (pH 7.0), 100 mM NaCl, 0.05% DDM to remove imidazole. Absorption spectra were recorded with a UV–vis spectrometer (V-750, JASCO).

**Transformation of *C. necator***

C. necator H16 competent cells were prepared following a protocol adapted from Pseudomonas aeruginosa 35. Briefly, strains were streaked on LB-agar and incubated for 48 h at 30 °C. Single colonies were used to inoculate 5 mL SOB medium and grown overnight (16-18 h). A 200 µL aliquot of saturated culture was transferred into 50 mL SOB and grown to $OD_{600}$ ≈ 0.8. Cells were pelleted at 4000 x g for 10 min, washed three times with a buffer containing 1 mM HEPES (pH 7.0), 1 mM $MgSO_4$, and 10% glycerol and resuspended in 0.5 mL of the same buffer. Sterile 50% glycerol was added to yield a final concentration of 15%.

For electroporation, 150 ng of plasmid DNA was mixed with 100 µL of competent cells on ice and transferred to a 0.2 cm cuvette. Electroporation was performed at 2.5 kV using a MicroPulser™ (Bio-Rad). Afterward, 0.95 mL of SOB + 20 mM D-fructose recovery medium was added, and cells were incubated for 2-4 h at 30 °C (150 rpm). Transformants were plated on selective LB-agar and incubated for up to 48 h at 30 °C.

**Rhodopsin expression and chromophore phenotype in *C. necator***

Four rhodopsin candidates were cloned into the same pLO11 plasmid backbone and named APR1, APR2, APR6, and APR7. As an initial indication that the spectral shift engineered into the APR variants was present upon expression in *C. necator*, cultures were grown with or without L-arabinose induction. Consistent with the blue shift in absorption from ~545 nm (GR) to ~450-500 nm (APR), induced APR-expressing cells displayed a distinctly yellow-orange colouration relative to both uninduced controls and GR-containing cells, which presented a deeper pink-red hue (Fig. 1).

**Rhodopsin-mediated phototrophic growth in engineered *C. necator***

Whether APR variants conferred a growth advantage over the previously characterised GR strain [1] was assessed using batch cultures grown in Chi.bio™ reactors, which enabled continuous $OD_{600}$ monitoring under defined illumination conditions. All cultures were normalised to $OD_{600}$ = 0.1 at the start of the experiment and grown in darkness for the first 8 h.

### Results

**Uninduced and dark-induced controls showed similar growth trajectories**

Before assessing strains under illumination, growth of induced cultures in the dark was first examined across all strains (Supplementary Fig. S1). As growth profiles were comparable in the absence of light, the pooled dark-induced condition was compared with uninduced controls grown under light or dark conditions (Supplementary Fig. S2). Both followed similar trajectories throughout the experiment, with only modest divergence at later time points, which was small relative to the effects subsequently observed in light-exposed strains. The dark-induced condition was therefore used as the principal reference for subsequent comparisons.

**APR strains exhibited significantly altered growth trajectories relative to the dark-induced control**

With the reference established, divergence between strains emerged after approximately 20–30 h (Fig 2.). Growth trajectories were compared using an ordinary least squares (OLS) regression model incorporating time (linear and quadratic terms) and strain × time interactions.

This analysis revealed a significant difference in growth kinetics between strains (likelihood ratio test, $p = 5.18 \times 10^{-8}$).

Relative to the dark-induced control, all APR strains exhibited significantly steeper increases in OD600 over time (APR1: $\beta = 0.0168$, $p = 2.12 \times 10^{-18}$; APR2: $\beta = 0.0162$, $p = 9.50 \times 10^{-12}$; APR6: $\beta = 0.0110$, $p = 4.39 \times 10^{-10}$; APR7: $\beta = 0.0153$, $p = 1.08 \times 10^{-15}$). In contrast, GR also showed a significant but smaller increase in growth rate ($\beta = 0.0058$, $p = 0.005$), indicating a weaker effect relative to the APR variants.

No major differences were observed in the curvature of the growth trajectories, as indicated by the absence of significant strain × time$^2$ interaction terms (Table S3). In contrast, significant strain × time interaction terms for all strains indicate that the observed differences were driven primarily by variation in growth rate during the exponential phase.

**Pairwise differences emerged from 20 h onward and were strongest for APR1 and APR7**

To determine when strains diverged, pairwise comparisons at each time point were performed using Welch's two-sample t-tests, with p-values adjusted using the Benjamini–Hochberg procedure to control the false discovery rate. No significant differences were detected during early growth before 20 h, indicating similar initial behaviour across all strains. At 20 h, APR1 was significantly higher than the dark induced control ($p = 0.00653$) (Fig. 3A).

By 25 h, APR1, APR6, and APR7 were significantly higher than the dark-induced control ($p = 1.85 \times 10^{-5}$, $2.47 \times 10^{-4}$, and $1.01 \times 10^{-3}$, respectively), whereas APR2 and GR did not differ significantly at this stage (Fig. 3B). By 40 h, this pattern had broadened: all APR variants were now significantly elevated relative to the dark-induced control (APR1: $p = 5.63 \times 10^{-4}$; APR2: $p = 6.26 \times 10^{-3}$; APR6: $p = 5.63 \times 10^{-4}$; APR7: $p = 5.25 \times 10^{-5}$), and APR1 and APR7 were also significantly higher than GR ($p = 0.0259$ and $p = 0.0166$, respectively), indicating a greater growth advantage for these variants during mid-to-late growth (Fig. 3C). These differences were maintained at 50 h, where all APR strains remained significantly above the control (APR1: $p = 1.87 \times 10^{-3}$; APR2: $p = 1.55 \times 10^{-3}$; APR6: $p = 1.16 \times 10^{-3}$; APR7: $2.47 \times 10^{-4}$). APR7 also continued to exceed GR ($p = 0.0232$), while other APR–GR comparisons were not significant after correction (Fig. 3D).

**Growth-rate analysis showed a stronger and more sustained light-dependent kinetic advantage in APR strains**

Time-resolved growth rates (μ) were calculated to examine how growth dynamics evolved over the 50 h experiment (Fig. 4). All strains exhibited an initial increase in growth rate during the early exponential phase, which peaked between 10–15 h and declined thereafter as cultures approached the stationary phase. Under illumination, APR-expressing strains consistently exhibited higher growth rates than the dark-induced control during the early-to-mid exponential phase (10–25 h). For example, APR1 reached a peak growth rate of $\mu = 0.139 \pm 0.013$ h$^{-1}$ at 15 h under light, compared with $\mu = 0.085 \pm 0.007$ h$^{-1}$ under dark conditions at the same time point. Similar illumination-dependent increases were observed for APR7 ($\mu = 0.136 \pm 0.023$ h$^{-1}$ under light vs $\mu = 0.061 \pm 0.011$ h$^{-1}$ under dark at 15 h) and APR6, which reached $\mu = 0.132 \pm 0.013$ h$^{-1}$ at 10 h under light compared with $\mu = 0.119 \pm 0.010$ h$^{-1}$ under dark.

APR2 showed a more modest but still detectable light-dependent effect, with peak growth rates of $\mu$ = 0.118 ± 0.019 $h^{-1}$ at 10–15 h but sustained higher values under illumination at later time points ($\mu$ = 0.073 ± 0.012 $h^{-1}$ at 25 h under light vs $\mu$ = 0.037 ± 0.004 $h^{-1}$ under dark). In contrast, GR displayed a comparatively weak and inconsistent response to illumination. Although GR reached a slightly higher peak under light ($\mu$ = 0.148 ± 0.020 $h^{-1}$ at 10 h) than under dark ($\mu$ = 0.119 ± 0.010 $h^{-1}$), this difference was not sustained, and growth-rate profiles converged thereafter ($\mu$ ≈ 0.041–0.052 $h^{-1}$ at 20–25 h under light vs $\mu$ ≈ 0.037–0.044 $h^{-1}$ under dark).

Growth rates were therefore averaged across the exponential phase (10–40 h) to provide a summary measure for each strain (Fig. 5). Mean growth rates were highest for APR2 ($\mu$ = 0.0596 ± 0.0072 $h^{-1}$), APR1 ($\mu$ = 0.0583 ± 0.0070 $h^{-1}$), and APR7 ($\mu$ = 0.0560 ± 0.0071 $h^{-1}$), differing by less than 0.004 $h^{-1}$. APR6 exhibited a lower mean growth rate ($\mu$ = 0.0508 ± 0.0054 $h^{-1}$), while GR showed the lowest growth rate among the light-exposed strains ($\mu$ = 0.0386 ± 0.0057 $h^{-1}$).

These differences were statistically significant (Welch one-way ANOVA, $F_{5,22.7}$ = 6.14, $p$ = 0.001). Pairwise Welch t-tests followed by Benjamini–Hochberg correction showed that all APR strains exhibited higher growth rates than the dark-induced control (adjusted $p < 0.05$), whereas GR did not differ significantly from the dark-induced condition (Table S4). In addition, GR was lower than APR1 and APR2 (adjusted $p < 0.05$), while differences among APR variants were not significant.

**APR strains exhibit an improved cellular energy state under illumination**

To determine whether the enhanced growth observed in APR strains was associated with changes in cellular energy status, ATP and ADP levels were quantified and expressed as ATP fraction and ADP/ATP ratio (Fig. 6). Illumination significantly altered both measures of cellular energy status across all strains. ATP fraction increased from 0.186 ± 0.004 under dark conditions to 0.252 ± 0.015 under illumination (two-way linear model, main effect of condition: $F_{1,19}$ = 18.49, $p$ = 0.0004), while the ADP/ATP ratio decreased from 4.42 ± 0.11 to 3.16 ± 0.24 ($F_{1,19}$ = 25.62, $p$ = 0.0001). Although all strains exhibited this light-dependent effect, the magnitude of the response did not differ significantly between strains (Genotype × Condition interaction: $F_{4,19}$ = 1.28, $p$ = 0.314 for ATP fraction; $F_{4,19}$ = 1.18, $p$ = 0.351 for ADP/ATP ratio). Within-strain comparisons between light and dark conditions were not statistically significant (Welch two-sample t-tests with Benjamini–Hochberg correction, all adjusted $p > 0.05$).

Taken together, these results demonstrate a consistent light-dependent growth advantage in APR-expressing strains, accompanied by a shift toward a more energy-rich cellular state.

### Supplementary tables

**Table S1**. Quantitative breakdown of the rhodopsin sequence database. Summary of functional classes, sequence counts (N = 884), and absorption wavelength (λmax ) statistics. Proton pumps dominate the dataset but are excluded from the blue extreme (< 500nm), which is occupied by channelrhodopsins.

| Functional Class | Count (n) | Prop. (%) | Median λmax (nm) | Mean λmax (nm) | Range (nm) |
|---|---|---|---|---|---|
| **Proton Pump** | 310 | 35.1 | 548.0 | 551.4 | 455-622 |
| **Proton Sensory** | 237 | 26.8 | 535.0 | 531.8 | 460-618 |
| **Sodium Pump** | 220 | 24.9 | 527.0 | 530.3 | 465-581 |
| **Chloride Pump** | 59 | 6.7 | 548.0 | 547.5 | 505-583 |
| **Channel** | 37 | 4.2 | 480.0 | 493.1 | 453-587 |
| **Unknown** | 21 | 2.4 | 534.0 | 525.1 | 436-568 |
| **Total Combined** | **884** | **100.0** | **537.0** | **539.9** | **436-622** |

**Table S2**. Pairwise Sequence and Structural Identity Statistics. Comparison of Levenshtein identity metrics in primary sequence versus secondary structure space for the unique proton pump lineage (n = 309). Primary identity shows high divergence, while secondary structure remains over 2.5-fold more conserved.

| Metric | Primary Sequence | Secondary Structure | Ratio (Sec/Pri) |
|---|---|---|---|
| Mean Identity | 47.50% | 85.44% | 1.80 |
| Median Identity | 27.17% | 82.06% | 3.02 |
| Standard Deviation | 31.75% | 8.50% | 0.27 |
| Min / Max Identity | 16.60% / 100.00% | 65.05% / 100.00% | 3.92 / 1.00 |
| E-Value < 0.01 | 23.9% | - | - |

**Table S3:** Strains of bacteria used in this study

| Strains | Comments |
|---|---|
| *E. coli* NEB5α | F- endA1 glnV44 thi-1 recA1 relA1 gyrA96 deoR nupG Φ80dlacZΔM15 Δ(lacZYA-argF)U169, hsdR17(rK- mK+), λ- (Commercial strain New England Biolabs, UK) |
| *Cupriavidus necator (Ralstonia eutropha* H16, ATCC 17699) | Wild-type strain |
| *Cupriavidus necator APR1* | Wild-type strain *C. necator* with APR1 plasmid |
| *Cupriavidus necator APR2* | Wild-type strain *C. necator* with APR2 plasmid |
| *Cupriavidus necator APR6* | Wild-type strain *C. necator* with APR6 plasmid |
| *Cupriavidus necator APR7* | Wild-type strain *C. necator* with APR7 plasmid |
| **Plasmids** | **Comments** |
| pLO11 ($Tc^r$, RK2 *ori*, $Mob^+$) | Expression vector for use in *R. eutropha* with $P_{BAD}$ promoter and downstream cloning sites (gift from Oliver Lenz, Technische Universität Berlin, Germany). |
| pLO11-GR | pLO11 containing the gene for GR rhodopsin from *Gloeobacter violaceus* PCC7421 |
| pLO11-APR1 | pLO11 containing ML designed rhodopsin APR1 |
| pLO11-APR2 | pLO11 containing ML designed rhodopsin APR2 |
| pLO11-APR6 | pLO11 containing ML designed rhodopsin APR6 |
| pLO11-APR7 | pLO11 containing ML designed rhodopsin APR7 |

**Table S4: Coefficients and p-values derived using the global growth-curve regression model**

| term | coef | std_err | stat | p_value | ci_lower | ci_upper |
|---|---|---|---|---|---|---|
| Intercept | 0.686 | 0.016 | 41.897 | 3.9E-219 | 0.654 | 0.719 |
| C(curve_group, Treatment(reference='Dark induced'))[T.APR1] | 0.406 | 0.041 | 9.963 | 2.6E-22 | 0.326 | 0.486 |
| C(curve_group, Treatment(reference='Dark induced'))[T.APR2] | 0.218 | 0.051 | 4.278 | 2.1E-05 | 0.118 | 0.318 |
| C(curve_group, Treatment(reference='Dark induced'))[T.APR6] | 0.256 | 0.038 | 6.755 | 2.5E-11 | 0.181 | 0.330 |
| C(curve_group, Treatment(reference='Dark induced'))[T.APR7] | 0.398 | 0.041 | 9.761 | 1.6E-21 | 0.318 | 0.478 |
| C(curve_group, Treatment(reference='Dark induced'))[T.GR] | 0.170 | 0.045 | 3.792 | 1.6E-04 | 0.082 | 0.258 |
| hours_c | 0.016 | 0.001 | 21.206 | 3.4E-82 | 0.015 | 0.017 |
| C(curve_group, Treatment(reference='Dark induced'))[T.APR1]:hours_c | 0.017 | 0.002 | 8.930 | 2.1E-18 | 0.013 | 0.020 |
| C(curve_group, Treatment(reference='Dark induced'))[T.APR2]:hours_c | 0.016 | 0.002 | 6.899 | 9.5E-12 | 0.012 | 0.021 |
| C(curve_group, Treatment(reference='Dark induced'))[T.APR6]:hours_c | 0.011 | 0.002 | 6.305 | 4.4E-10 | 0.008 | 0.014 |
| C(curve_group, Treatment(reference='Dark induced'))[T.APR7]:hours_c | 0.015 | 0.002 | 8.155 | 1.1E-15 | 0.012 | 0.019 |
| C(curve_group, Treatment(reference='Dark induced'))[T.GR]:hours_c | 0.006 | 0.002 | 2.805 | 5.1E-03 | 0.002 | 0.010 |
| hours_c2 | -3.72E-04 | 5.96E-05 | -6.247 | 6.3E-10 | -4.89E-04 | -2.55E-04 |
| C(curve_group, Treatment(reference='Dark induced'))[T.APR1]:hours_c2 | -2.29E-04 | 1.48E-04 | -1.544 | 1.2E-01 | -5.20E-04 | 6.21E-05 |
| C(curve_group, Treatment(reference='Dark induced'))[T.APR2]:hours_c2 | 1.21E-04 | 1.85E-04 | 0.655 | 5.1E-01 | -2.42E-04 | 4.85E-04 |
| C(curve_group, Treatment(reference='Dark induced'))[T.APR6]:hours_c2 | -1.30E-04 | 1.38E-04 | -0.941 | 3.5E-01 | -4.00E-04 | 1.41E-04 |
| C(curve_group, Treatment(reference='Dark induced'))[T.APR7]:hours_c2 | -2.04E-04 | 1.48E-04 | -1.373 | 1.7E-01 | -4.95E-04 | 8.75E-05 |
| C(curve_group, Treatment(reference='Dark induced'))[T.GR]:hours_c2 | -3.38E-05 | 1.63E-04 | -0.207 | 8.4E-01 | -3.54E-04 | 2.87E-04 |

The linear time term (hours_c) represents growth rate, and strain × time interactions capture differences in growth rate between strains. The quadratic term (hours_c²) describes curvature, while strain × time² interactions represent differences in curve shape.

**Table S5: Exponential-phase (10-40 h) growth rate across strains using pairwise Welch t-tests with Benjamini–Hochberg correction.**

| Group A | Group B | n(A) | n(B) | Mean A | Mean B | Mean diff. | t-stat | p-value raw | p-value adj. | Reject $H_0$ |
|---|---|---|---|---|---|---|---|---|---|---|
| GR | APR1 | 8 | 10 | 0.0386 | 0.0583 | -0.0197 | -2.7822 | 0.0181 | 0.0454 | TRUE |
| GR | APR2 | 8 | 6 | 0.0386 | 0.0596 | -0.0209 | -3.4798 | 0.0116 | 0.0430 | TRUE |
| GR | APR6 | 8 | 12 | 0.0386 | 0.0508 | -0.0122 | -2.4019 | 0.0295 | 0.0617 | FALSE |
| GR | APR7 | 8 | 10 | 0.0386 | 0.0560 | -0.0174 | -2.4457 | 0.0329 | 0.0617 | FALSE |
| GR | Dark Induced | 8 | 52 | 0.0386 | 0.0353 | 0.0033 | 1.0585 | 0.2992 | 0.4488 | FALSE |
| APR1 | APR2 | 10 | 6 | 0.0583 | 0.0596 | -0.0012 | -0.1418 | 0.8893 | 0.8893 | FALSE |
| APR1 | APR6 | 10 | 12 | 0.0583 | 0.0508 | 0.0075 | 0.9203 | 0.3707 | 0.5055 | FALSE |
| APR1 | APR7 | 10 | 10 | 0.0583 | 0.0560 | 0.0023 | 0.2446 | 0.8095 | 0.8674 | FALSE |
| APR1 | Dark Induced | 10 | 52 | 0.0583 | 0.0353 | 0.0230 | 3.2319 | 0.0078 | 0.0392 | TRUE |
| APR2 | APR6 | 6 | 12 | 0.0596 | 0.0508 | 0.0088 | 1.2061 | 0.2519 | 0.4198 | FALSE |
| APR2 | APR7 | 6 | 10 | 0.0596 | 0.0560 | 0.0036 | 0.4072 | 0.6901 | 0.7962 | FALSE |
| APR2 | Dark Induced | 6 | 52 | 0.0596 | 0.0353 | 0.0243 | 3.9980 | 0.0056 | 0.0392 | TRUE |
| APR6 | APR7 | 12 | 10 | 0.0508 | 0.0560 | -0.0052 | -0.6328 | 0.5356 | 0.6695 | FALSE |
| APR6 | Dark Induced | 12 | 52 | 0.0508 | 0.0353 | 0.0155 | 3.0210 | 0.0077 | 0.0392 | TRUE |
| APR7 | Dark Induced | 10 | 52 | 0.0560 | 0.0353 | 0.0207 | 2.8963 | 0.0143 | 0.0430 | TRUE |

**Table S6:** Oligonucleotides used in this study

| Name | Sequence | Type |
|---|---|---|
| pLO11_GA_F1 | CGGCCGCAGATCTGCTTG | Cloning |
| pLO11_GA_490_1_R1 | CGCATGGGGTCTCCTCCTTAGCTAGC | Cloning |
| Sch1_490_1_F | TAAGGAGGAGACCCCATGCG | Cloning |
| Sch1_490_1_R | CAAGCAGATCTGCGGCCGTCAGG | Cloning |
| pLO11_GA_490_2_R1 | AGTAATTTCATGGGGTCTCCTCCTTAGCTAGC | Cloning |
| Sch1_490_2_F | TAAGGAGGAGACCCCATGAAATTACT | Cloning |
| Sch1_490_2_R | CAAGCAGATCTGCGGCCGTTAGGC | Cloning |
| pLO11_GA_490_6_R1 | CAGTTTCATGGGGTCTCCTCCTTAGCTAGC | Cloning |
| Sch1_490_6_F | TAAGGAGGAGACCCCATGAAACTG | Cloning |
| Sch1_490_6_R | CAAGCAGATCTGCGGCCGTCAAG | Cloning |
| pLO11_GA_490_7_R1 | ACCCATGGGGTCTCCTCCTTAGCTAGC | Cloning |
| Sch1_490_7_F | TAAGGAGGAGACCCCATGGGT | Cloning |
| Sch1_490_7_R | CAAGCAGATCTGCGGCCGTCAGT | Cloning |
| Sch1_490_1_SeqF1 | CACCCTGGTGATGCTGATCG | Sequencing |
| Sch1_490_1_SeqR1 | TTGTTCACGAAGTCGGCCAG | Sequencing |
| Sch1_490_2_SeqF1 | GCATGGCCCGCATTTATCATC | Sequencing |
| Sch1_490_2_SeqR1 | CTTTCTTTGACAGCGACGTTCC | Sequencing |
| Sch1_490_6_SeqF1 | TAGTCGGGTCGCTGGTCATG | Sequencing |
| Sch1_490_6_SeqR1 | CCAAATGATCGTCCCAAACAGG | Sequencing |
| Sch1_490_7_SeqF1 | TCGCCGCATCGTTATTCAAG | Sequencing |
| Sch1_490_7_SeqR1 | TAAGATTCAGGGCAGACACG | Sequencing |
| PLO11_Flank_F1 | GGCTCTTCTCGCTAACCAAAC | Sequencing |
| PLO11_Flank_R1 | GAACGTGGCGAGAAAGGAAG | Sequencing |

### Figures

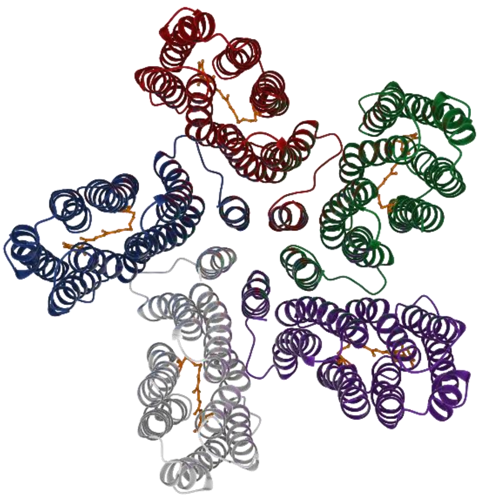

APR1

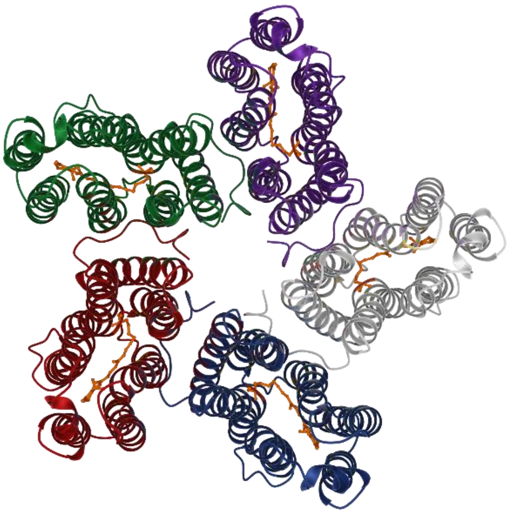

APR2

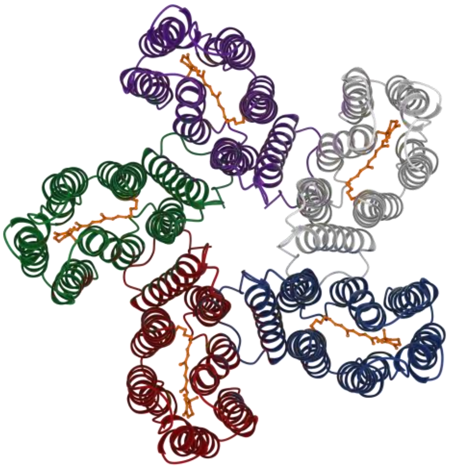

APR6

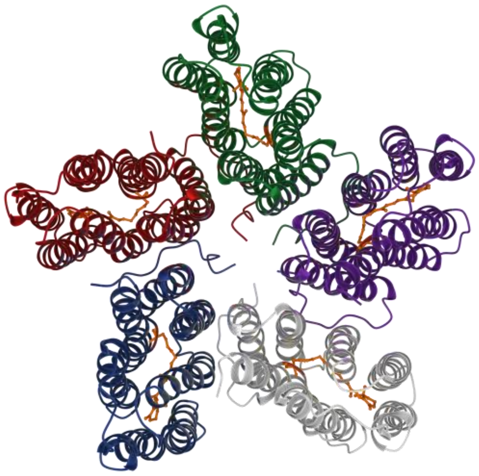

APR7

**Figure BS1.** AI predicted structures of APR1, APR2, APR6 and APR7.

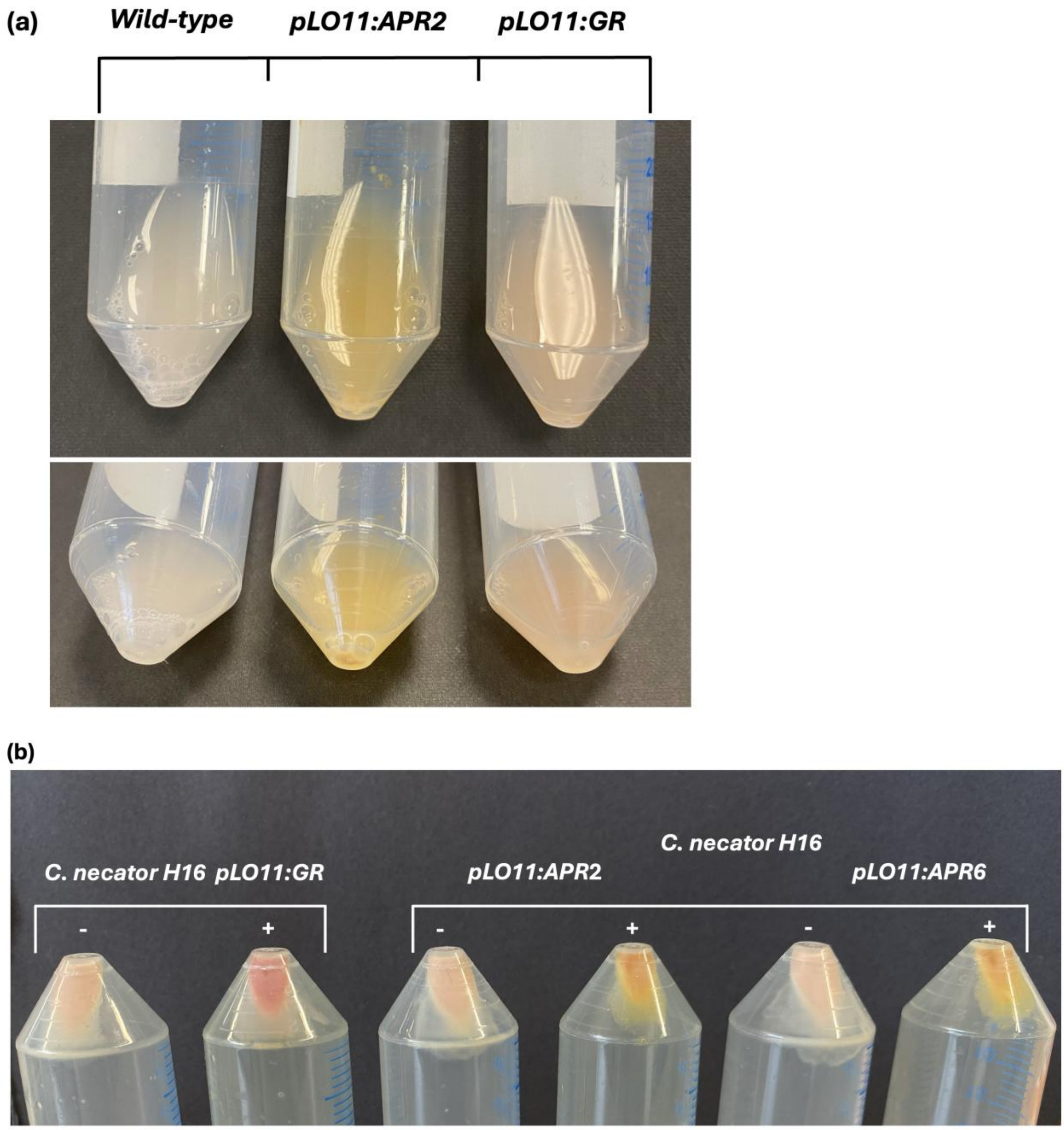


**Figure BS2.** Cell colouration confirming APR expression in *C. necator* H16. *C. necator* H16 strains harbouring *pLO11:GR, pLO11:APR2*, or *pLO11:APR6* were grown in minimal medium containing 5 µg $ml^{-1}$ all-*trans*-retinal and 80 mM formate. (a) Liquid cultures of wild-type, $GR^{+}$, and $APR2^{+}$ strains, showing the shift in colour from pink-red ($GR^{+}$) to yellow-orange ($APR2^{+}$), consistent with the engineered blue shift in rhodopsin absorption maxima from ~545 nm to ~450-500 nm and indicative of functional chromophore incorporation. (b) Centrifuged pellets from GR-induced (left), APR2-induced (centre), and APR6-induced (right) *C. necator* H16 cultures. Arabinose induction (+) and uninduced controls (−) are shown for each strain.

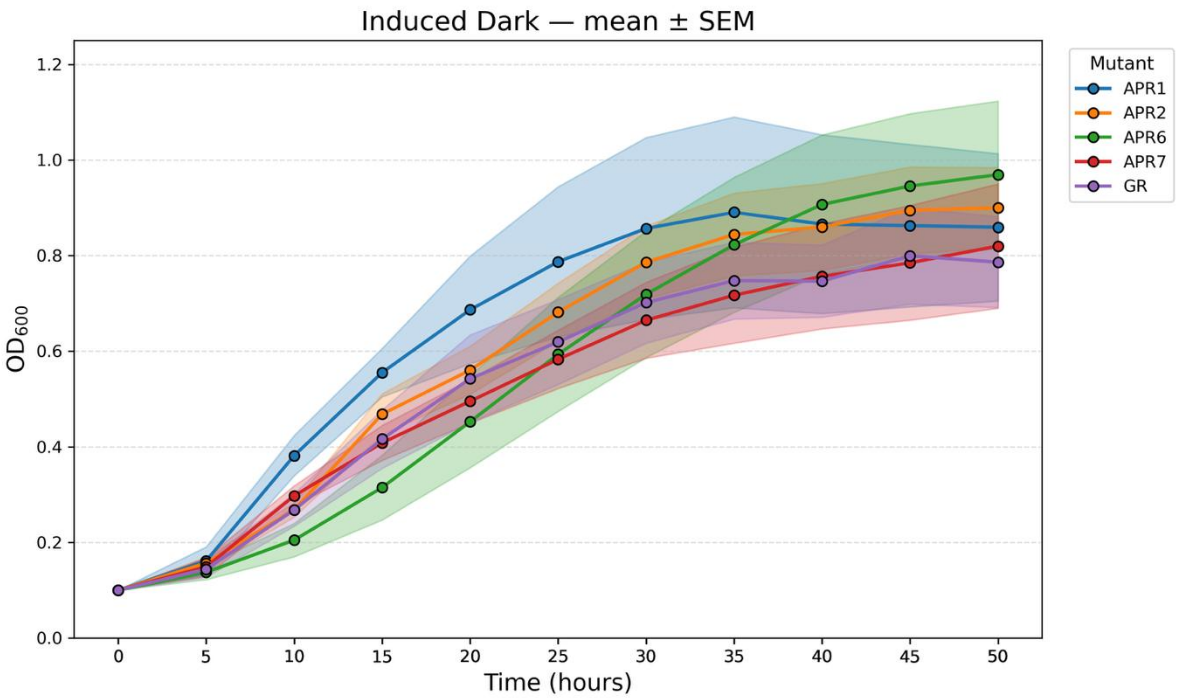


**Figure BS3. Growth of individual strains under induced dark conditions.** Growth curves for each strain grown under induced dark conditions. Data represent mean ± SEM (n = 4–18 biological replicates per strain).

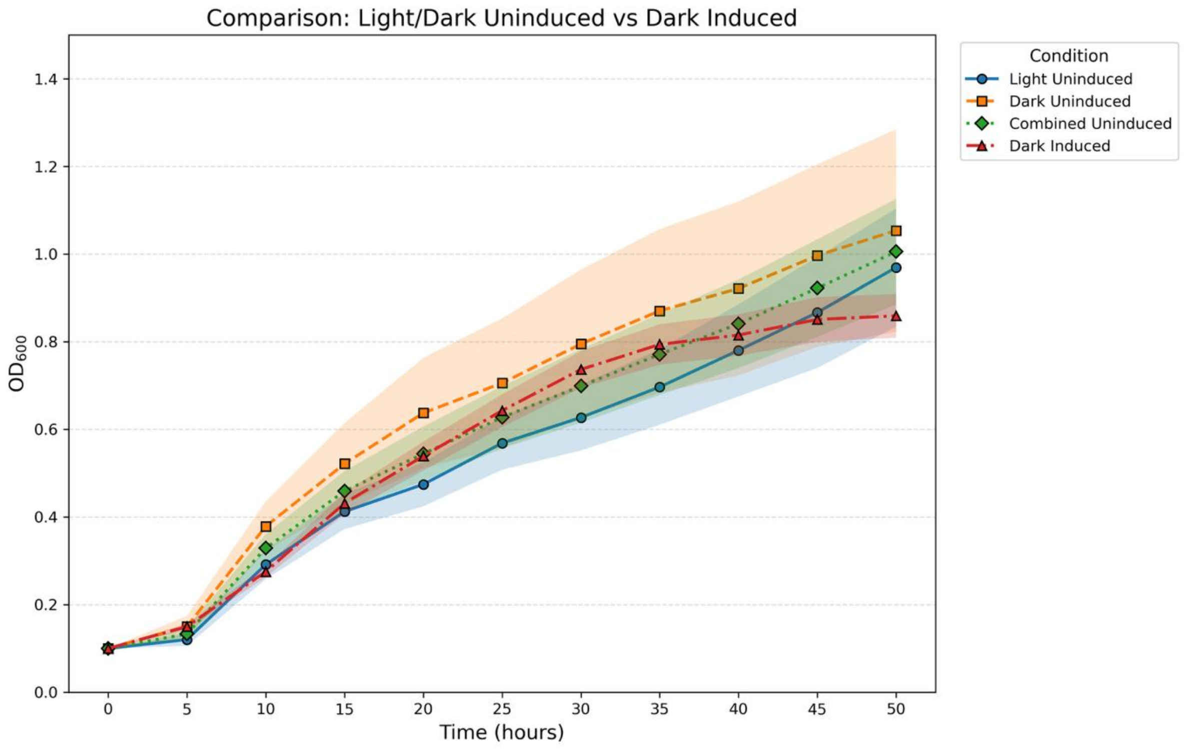


**Figure BS4. Comparison of the pooled uninduced controls with the pooled dark-induced control.** Growth curves (mean ± SEM) for Light Uninduced (n=8), Dark Uninduced (n=6), Combined Uninduced (Light and Dark, n=14), and Dark Induced (n=52) conditions across the 50 h cultivation period.

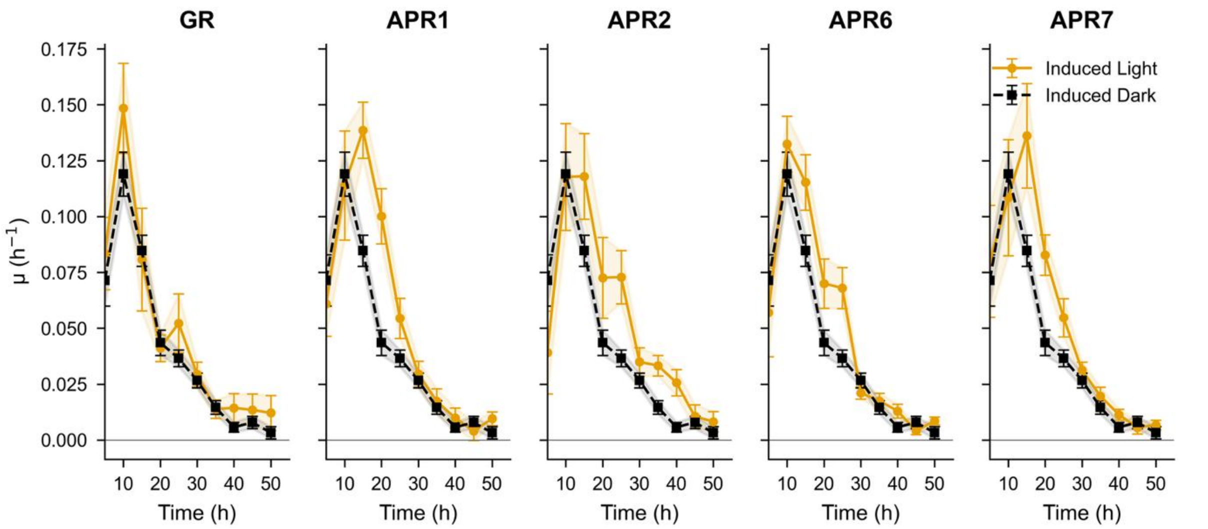


**Figure BS5. Time-resolved growth-rate profiles under induced light and induced dark conditions.** Specific growth rate (μ, $h^{-1}$) plotted over time for each strain under induced light and induced dark conditions. Data represent mean ± SEM in 5 hour intervals (n = 6–12 biological replicates per strain; n = 52 pooled control measurements)
